# First Generation Tools for the Modeling of Capicua (CIC) - Family Fusion Oncoprotein-Driven Cancers

**DOI:** 10.1101/2025.05.13.653825

**Authors:** Cuyler Luck, Kyle A. Jacobs, Julia Riad, Christopher D. Macaraig, Rovingaile Kriska M. Ponce, Ross A. Okimoto

**Affiliations:** Biomedical Sciences Graduate Program, University of California, San Francisco, CA, USA; Department of Medicine, University of California, San Francisco, CA, USA; Department of Cell and Tissue Biology, University of California, San Francisco, CA, USA; University of San Francisco, San Francisco, CA, USA; Boston University Chobanian and Avedisian School of Medicine, Boston, MA, USA; Helen Diller Comprehensive Cancer Center, University of California, San Francisco, CA, USA

**Keywords:** CIC::DUX4, CIC::NUTM1, CIC::LEUTX, ATXN1::DUX4, fusion, capicua, sarcoma, model, plasmid, synthetic

## Abstract

Clinical divergence between patients harboring CIC-rearrangements is frequently observed. For example, the prototypical CIC::DUX4 fusion associates with soft tissue tumors while CIC::NUTM1 fusions typically localize to the CNS (brain/spinal cord). The basis for these differences is poorly understood due to a lack of molecular tools. To address this need, we generated patient-informed, synthetic coding sequences for CIC::NUTM1, CIC::LEUTX, and ATXN1::DUX4 and validated them in structure-function studies. We found that CIC::NUTM1 drives a transcriptional program distinct from that of CIC::DUX4 due to a C-terminal NUTM1 functional domain, CIC::LEUTX weakly activates CIC target genes through LEUTX transactivation sequences, and ATXN1::DUX4 upregulates CIC target genes via the ATXN1 AXH domain. Our findings indicate that the CIC fusion binding partner may alter overall fusion oncoprotein activity. Thus, these first generation synthetic tools provide an unprecedented resource to study CIC-family fusions beyond CIC::DUX4 and allow for the dissection of this rare subgroup of cancers.

**Graphical Abstract:** 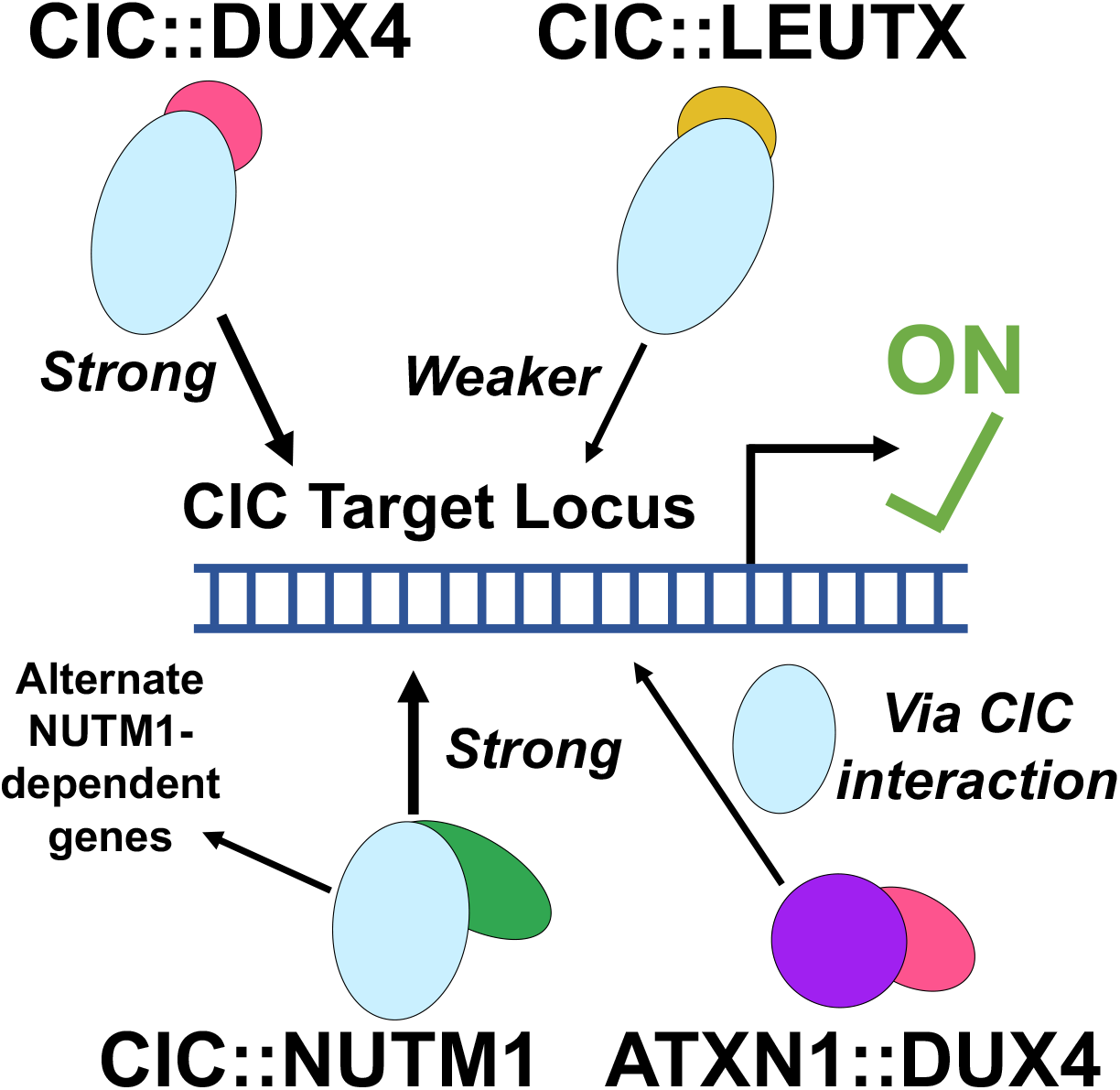

## Introduction

Transcription factor (TF) fusion oncoproteins often possess the ability to rewire cellular states to promote cellular transformation and malignant progression. It has become increasingly clear that TF fusion oncogenes and the diseases they cause are not strictly limited to a 1:1 pairing – highly similar or identical fusions may cause different cancers, and groups of fusions that vary in one or both partner genes may collectively drive a singular family of tumors (termed fusion gene promiscuity, reviewed here^1–3^). Our research focuses on sarcomas, where several types of pediatric and young adult cancers are driven by fusion oncoproteins (e.g. Ewing sarcoma – EWSR1::FLI1, alveolar rhabdomyosarcoma – PAX3/7::FOXO1, synovial sarcoma – SS18:SSX1/2). These tumors also consist of relatively promiscuous fusions, where there remains debate over whether differences in the partner genes comprising the driver fusion oncogene may have implications for tumor biology and patient outcomes. In alveolar rhabdomyosarcoma for example, PAX3::FOXO1 vs PAX7::FOXO1 prognostic data is quite mixed^4–6^, while recent isogenic modeling work in fibroblasts suggests that these two versions of the fusion may possess unique DNA binding sites and downstream gene activation^7^. As molecular diagnostics improve in the clinic, an improved understanding of whether such differences in gene fusion partners translates to divergent mechanisms of tumorigenesis and patient outcomes will be important to tailor treatment strategies.

Our group studies the gene capicua (*CIC*), which encodes a developmental transcription factor^8^ that acts as a default repressor and serves to negatively regulate the expression of cell cycle genes and MAPK pathway regulators^9,10^. As a tumor suppressor, both genetic and functional inactivation of wild-type CIC is associated with several types of cancer^11–15^. In a subset of undifferentiated round cell sarcoma, the C-terminal end of wild type (WT) CIC is replaced with another TF with transactivating capacity to generate a relatively promiscuous family of gene fusions. Mechanistically, the founding member of this family, CIC::DUX4, is thought to activate transcription through the recruitment of the histone acetyltransferases p300/CBP by the DUX4 transactivating domain to DNA binding sites determined by a largely intact CIC fragment^16–21^. Recent studies have expanded the spectrum of *CIC* rearrangements beyond those involving *DUX4*, with the most common alternative 3’ partner genes being *NUTM1*^22–28^, *LEUTX*^29–32^, and *FOXO4*^33–35^. These additional 3’ partners all possess evidence that they interact with p300/CBP^36–40^, lending support to the current consensus that *CIC*-rearrangements may recruit p300/CBP to activate CIC target genes. Fusions with a 5’ gene partner of *ATXN1* or *ATXN1L*, homologs which interact with CIC and modulate its stability^41,42^, have also been reported with 3’ partners of *DUX4*^43,44^, *NUTM1*^45^, or *NUTM2A*^46^. Where performed, the methylation profiles of tumors bearing these *ATXN1*-/*ATXN1L*-fusions have grouped with those harboring *CIC* rearrangements^43,44,46^, suggesting they together comprise a larger group perhaps more aptly termed “*CIC*-family” fusion oncogenes.

The presumed mechanism of action of *CIC*-fusion oncogenes (p300/CBP recruitment to CIC binding sites on DNA) would imply that all *CIC*-rearranged cancers should drive equivalent molecular programs and would present similarly in the clinic, regardless of 3’ fusion partner. However, the available data suggest that there may be differences between *CIC* fusions harboring different 3’ partner genes. Transcriptionally, the limited RNA-sequencing data from patients shows that while *CIC::DUX4, CIC::NUTM1*, and *CIC::FOXO4* positive tumors cluster away from other fusion-driven tumors, they also somewhat subdivide based on 3’ fusion partner^23^. Structurally, we have previously observed that *CIC::NUTM1* fusion sequences tend not to retain the CIC C1 domain, which is almost universally preserved in *CIC::DUX4* fusions and appears to be required for maximal function of CIC::DUX4^18^. Anatomically, *CIC::DUX4* tumors are largely a tumor of the soft tissue, while *CIC::NUTM1* tumors appear to predominantly present in the central nervous system (CNS) and spine^22–24^, as we have reviewed recently^47^. Furthermore, the rare tumors with *CIC*-family fusions in the brain have recently been divided into two diagnostic entities: sarcomas and high-grade neuroepithelial tumors (HGNET-CIC) which cluster separately by methylation profiling and have distinct morphological features^29,48^. Notably, while patient numbers are limited, the majority of HGNET-CIC tumors appear to carry the *CIC::LEUTX* rearrangement, while the distribution of fusion partners in CNS sarcoma with *CIC*-family rearrangement is much more varied^48^. These comparisons all struggle from small patient numbers and scarce data, but collectively suggest that the manifestation of *CIC*-family fused tumors may vary based on the fusion partners.

These data are likely explained by at least one of two hypotheses. The first is that different combinations of *CIC*-family fusion partners may be more likely to arise in various cell types, which could explain their distinct anatomical locations. This is a difficult hypothesis to test as the cell of origin is not known for any *CIC*-family fusion. A second possibility is that the molecular mechanism of action for *CIC*-family fusions could vary depending on the partner genes and may not simply be activation of *CIC* target genes by p300/CBP. In this scenario, the unique functionality imparted by different partner genes could be sufficient to modify the overall behavior of the fusion and enable it to transform different cell types. This too is difficult to investigate, as there are very few tools to study *CIC::DUX4* (a handful of plasmids^16,49^, patient-derived cell lines^50–52^, and transgenic mice^17^ have been developed), and none of any kind exist to model the other *CIC*- or *ATXN1*/*1L*-fusions in the lab. Waiting to encounter a patient with a fusion-bearing tumor, cloning that fusion and/or using the tumor to establish xenografts or cell lines, and using those tools for modelling is painfully slow when the tumor type in question is exceedingly rare and understudied, as is the case here. However, if there is existing data describing the breakpoint sequence of the fusion in patients, an alternative approach is to manually clone synthetic but patient-informed fusion oncogenes from fragments of the partner genes. Such an approach is certainly subject to caveats but is relatively quick and could allow for a head start on studying fusions that lack tools while traditional models are developed.

In this study we employ such an approach to create the first generation of molecular tools for studying non-*CIC::DUX4 CIC*-family fusions including *CIC::NUTM1, CIC::LEUTX*, and *ATXN1::DUX4*. We then employ these tools for structure-function studies to determine how the various partner genes contribute to the activity of their fusions, and how they may converge or diverge from each other. We find that CIC::NUTM1 fusions are capable of driving a slightly distinct transcriptional program from that of CIC::DUX4, and nominate a novel C-terminal NUTM1 functional domain which may explain this behavior. We additionally show CIC::LEUTX to be a relatively weak transactivator, with dependence on two small C-terminal motifs to execute this function. Finally, we show evidence to support the initial hypothesis^43^ that ATXN1::DUX4 may function by colocalizing the DUX4 moiety with CIC via the ATXN1 AXH domain, yielding a new mechanism as an “indirect *CIC* fusion”. Together, these tools enable the first modeling of these ultra-rare fusion oncogenes and yield mechanistic insights that may help to explain why *CIC*-family fusions can diverge in clinical settings. Looking forward, we aim to lower the barrier to entry into these emerging fields of research by making these tools publicly available in the hopes that we may collectively work to develop better models and new treatments for cancers driven by *CIC*-family fusion oncogenes.

## Results

### Generation of synthetic, patient-informed CIC-family fusions

The absence of molecular models for *CIC*-family fusions aside from *CIC::DUX4* poses a major roadblock to answering basic mechanistic questions about how different *CIC*-family fusions function. Direct cloning out of patient samples was not feasible due to the rarity of these tumors, and gene synthesis was not ideal due to the long and complex coding sequences of these fusions. Instead, we chose to leverage the fact that several studies have published trans-breakpoint Sanger sequencing of *CIC::NUTM1, CIC::LEUTX*, and *ATXN1::DUX4* identified in patients. This information allows for cloning fusion coding sequences from existing plasmids that contain the coding sequences for individual partner genes via a modified Gibson assembly approach (NEBuilder HiFi DNA Assembly). Using such a strategy, we chose four reported breakpoints^23,30,43^ and used them to generate four synthetic, patient-informed, in-frame N-terminally HA-tagged CIC-family fusions in a CMV-driven transfection backbone: *HA-CIC(ex18)::NUTM1(ex3), HA-CIC(ex20)::NUTM1(ex6), HA-CIC::LEUTX, and HA-ATXN1::DUX4* (Figure 1, Supplemental Figures S1-S2, further sequence details are available in the Methods section). These constructs, in concert with an existing *HA-CIC::DUX4* plasmid^16^, allow for expression of various *CIC*-family fusions in cells.

**Figure 1:**
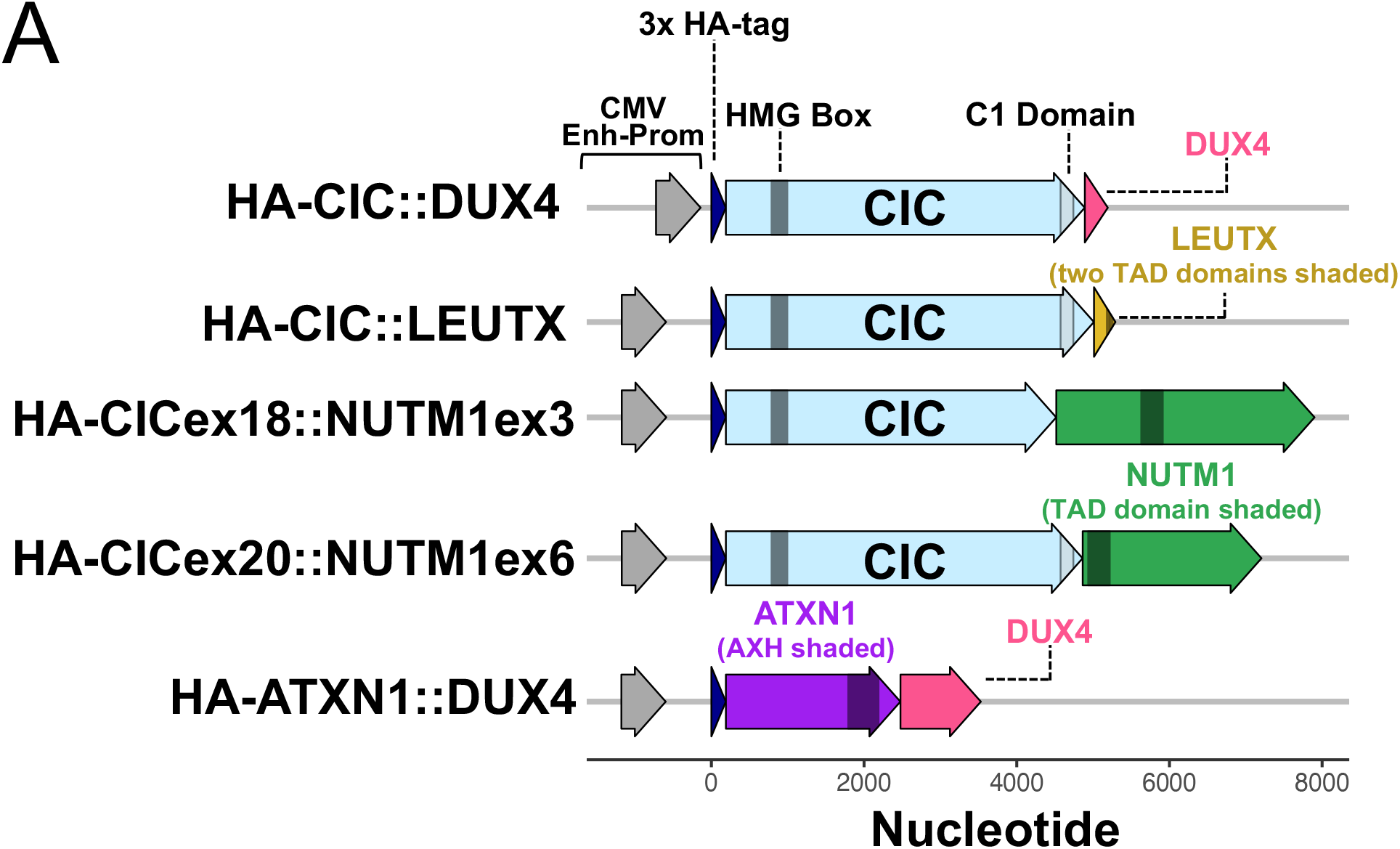
Structure of plasmids encoding *CIC*-family fusions. (A) To-scale representation of key regulatory and coding sequences in the original HA-CIC::DUX4 plasmid^16^ and the four synthetic *CIC*-family fusions. All plasmids are driven by a CMV enhancer-promoter region (“CMV Enh-Prom”) though the exact sequence varies slightly between the HA-CIC::DUX4 plasmid and the others, owing to slightly different backbones (a more complete description is available in the Methods). Shaded regions for the LEUTX and NUTM1 fragments indicate previously identified or nominated transactivating motifs that interact with p300/CBP. The shaded region for ATXN1 indicates the AXH domain, which mediates interaction with CIC. The CIC HMG-box (dark gray) and C1 (light gray) DNA-binding domains are indicated.

For *CIC::NUTM1*, we cloned two versions due to our prior interest in the CIC C1 domain, which works with the CIC HMG box to bind DNA^53^. We previously found that this functional domain is retained in almost all CIC::DUX4 patient fusions, and is necessary for maximal transcriptional activity of the fusion oncoprotein^18^. However, we also noted that in our previous breakpoint dataset^18^, the majority of CIC::NUTM1 sequences identified in patients did not retain the C1 domain. This led us to clone two CIC::NUTM1 constructs based on two independent patient breakpoints: one that does retain the C1 domain (CIC(ex20)::NUTM1(ex6)) and one that does not (CIC(ex18)::NUTM1(ex3)) (Figure 1).

### Molecular and functional characterization of the CIC::NUTM1 fusion oncoprotein

To characterize the behavior of these fusion oncoproteins we chose to perform cell-based experiments using the 293T cell line. Derivatives of HEK293 cells, including 293T, are a frequent model of choice in structure-function studies of fusion oncogenes^37,39,54–57^ and have the advantage of enabling rapid isogenic comparisons while being easy to work with. Additionally, HEK293 derivatives express wild type CIC and consequently do not express high levels of many known CIC target genes, yielding an easily measurable increase in CIC target gene expression when wild type *CIC* is perturbed or when *CIC*-fusions are overexpressed^10,18^.

When overexpressed in 293T cells, both CIC::NUTM1 isoforms markedly increased ETV5 expression, a known CIC^10,53,58^ and CIC::DUX4^16^ target gene, at levels comparable to that seen by CIC::DUX4 and they both demonstrated nuclear localization (Figure 2A and 2B). We tested whether CIC::NUTM1 is a *bona-fide* fusion that requires both partner genes for activity, as opposed to acting as a truncation mutant of one or both partners, by independently expressing the fractional partner genes and comparing their ability to upregulate ETV5 relative to the full-length (FL) fusion. These studies indicate that while the FL CIC::NUTM1 fusion could induce ETV5 expression, neither truncated partner alone was sufficient, suggesting that CIC::NUTM1 is dependent on both partners to upregulate ETV5 (Figure 2C).

**Figure 2:**
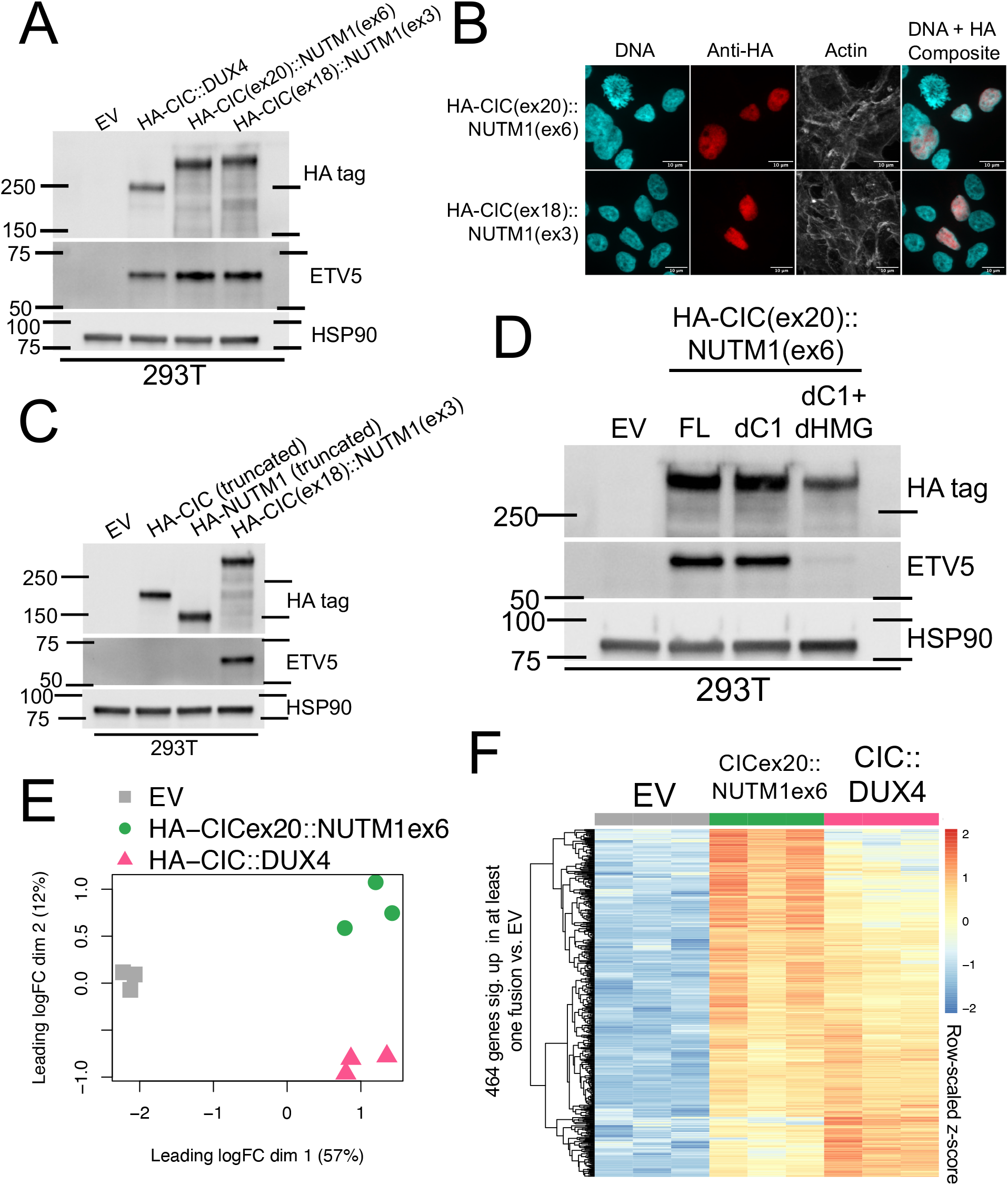
Functional validation of the synthetic CIC::NUTM1 fusion oncoproteins. (A) Immunoblot of 293T cells approximately 48 hours after transfection with empty vector (EV) or the indicated constructs, representative of three independent experiments. (B) Immunofluorescence microscopy of 293T cells approximately 48 hours after transfection with the indicated constructs and stained with DAPI (DNA), rhodamine-phalloidin (Actin), or an anti-HA antibody. Scale bars indicate 10 μm, representative cells are shown from one experiment. (C) Immunoblot of 293T cells approximately 48 hours after transfection with empty vector (EV), full length HA-CIC(ex18)::NUTM1(ex3), or truncated versions of HA-CIC(ex18)::NUTM1(ex3) that are comprised only of the HA tag motif and the fractional coding sequence of the indicated partner, split at the breakpoint. Representative of three independent experiments. (D) Immunoblot of 293T cells approximately 48 hours after transfection with empty vector (EV) or one of three versions of HA-CIC(ex20)::NUTM1(ex6): full length (FL), CIC C1 domain deleted (dC1), or CIC C1 domain and HMG box deleted (dC1 + dHMG). Representative of three independent experiments. (E) edgeR multi-dimensional scaling plot of RNA-seq data from 293T cells 48 hours after transfection with empty vector (EV) or the indicated fusion-expressing constructs. (F) Row-scaled RNA-seq expression heatmap of 464 significantly upregulated genes (log_2_ fold change > 1.5, q-value < 0.001) in at least one of the fusion-transfected 293T conditions vs EV.

We unexpectedly observed that both CIC::NUTM1 isoforms could increase ETV5 to a similar extent (Figure 2A) despite CIC(ex18)::NUTM1(ex3) lacking the C1 domain. This was surprising since we previously found that when the CIC::DUX4 fusion lacked the C1 domain its ability to transcriptionally upregulate *ETV5* was impaired^18^. Thus, we directly tested the necessity of the C1 domain for CIC::NUTM1 to activate ETV5 by deleting it alone or together with the HMG box from CIC(ex20)::NUTM1(ex6), and found that loss of the C1 domain alone had little impact on ETV5 induction, while loss of the C1 domain and HMG box together abrogated expression (Figure 2D).

These molecular findings coupled with the observed clinical differences between CIC::NUTM1 and CIC::DUX4 suggested that these fusions may potentially regulate unique gene targets. To evaluate this, we performed bulk RNA-sequencing (RNA-seq) of 293T cells expressing either CIC::DUX4 or CIC(ex20)::NUTM1(ex6). Our data indicated primary separation on a multidimensional scaling plot based on the presence or absence of the fusion oncoprotein, but we additionally observed a clear secondary separation between CIC::DUX4 and CIC(ex20)::NUTM1(ex6) expressing cells (Figure 2E). Indeed, a heatmap of all genes significantly upregulated in at least one of the fusion-transfected conditions vs. control showed that while many genes were significantly increased by both CIC::DUX4 and CIC(ex20)::NUTM1(ex6), there were gene clusters that were differentially regulated by one fusion relative to the other (Figure 2F). Together, these results suggest that CIC::NUTM1 is a *bona-fide* fusion oncoprotein that overlaps with but may in part differ in functional domain requirements and transcriptional output from CIC::DUX4.

### Differential target gene regulation by CIC::NUTM1 and CIC::DUX4

To further investigate which genes were preferentially activated by CIC(ex20)::NUTM1(ex6) vs CIC::DUX4, we first validated that expression of the canonical CIC targets *ETV1, ETV4*, and *ETV5* were strongly upregulated by both fusions, regardless of the 3’ partner gene (Figure 3A). Next, we noted upregulation of multiple forkhead box TF family members by CIC::DUX4 and CIC(ex20)::NUTM1(ex6), which are broadly implicated in development. Of these genes, we were particularly struck by the presence of *FOXD3, FOXB1*, and *FOXG1*, which have been previously implicated in the development of various neural structures and cell types^59^. Importantly, both *FOXD3* and *FOXB1* were specifically upregulated in the CIC(ex20)::NUTM1(ex6) condition but not in the CIC::DUX4 expressing cells, and *FOXG1* met the significance threshold in the CIC(ex20)::NUTM1(ex6) but was slightly under the log_2_ fold change cutoff (while meeting neither cutoff in the CIC::DUX4 condition) (Figure 3A). These were striking to us because of the predilection for CIC::NUTM1 tumors to present in the CNS and spine of patients, whereas CIC::DUX4 tumors typically arise in the soft tissue. We also noted the shared upregulation of *FOXC2* (which is involved in the epithelial to mesenchymal transition^60^) by both fusions, as well as the upregulation of the stemness factor *SOX2* specifically in the CIC(ex20)::NUTM1(ex6) condition (Figure 3A).

**Figure 3:**
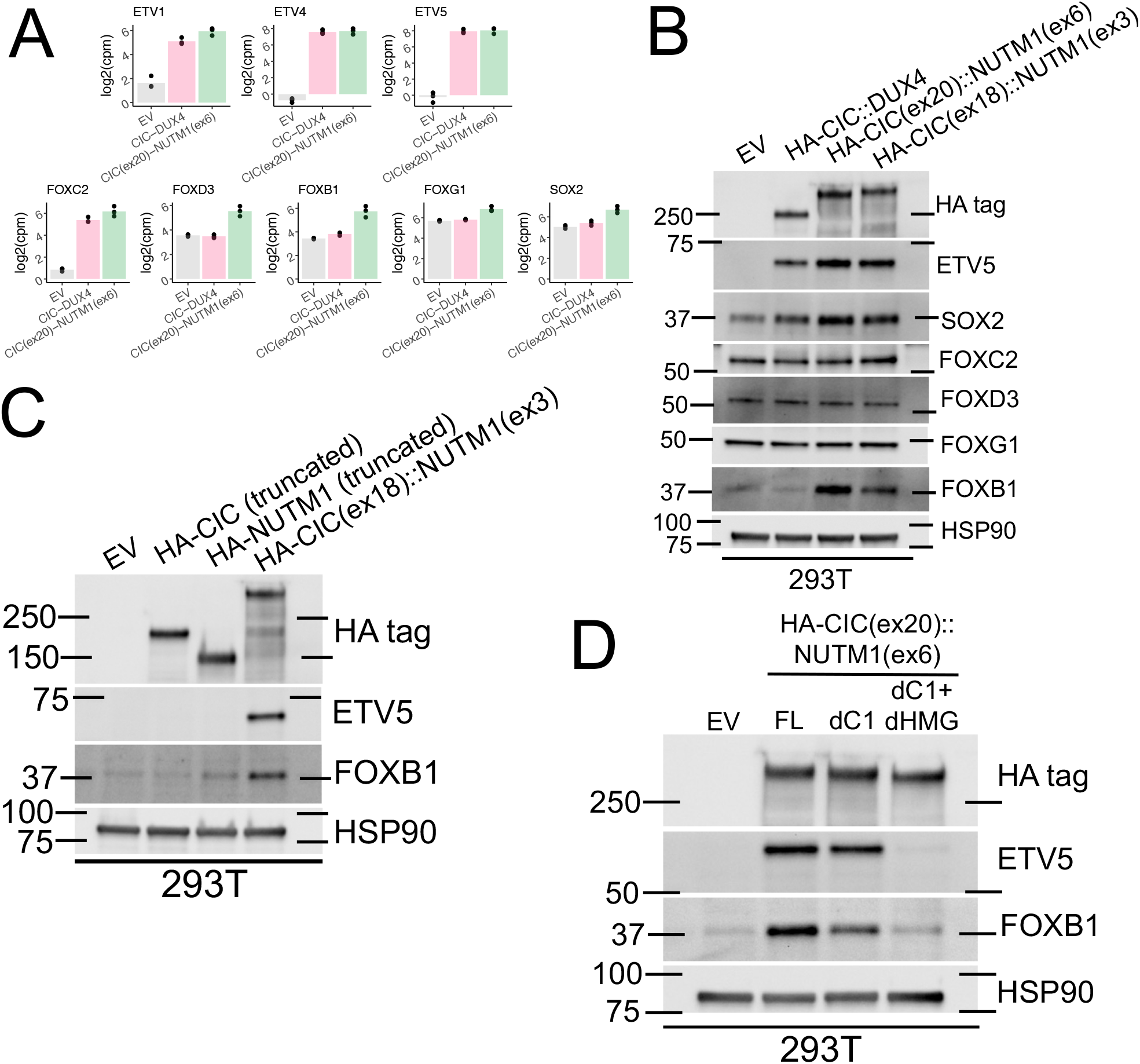
Differential gene expression analysis identifies *FOXB1* as a CIC::NUTM1 fusion specific target. (A) log_2_ counts per million plots of the indicated genes in 293T cells 48 hours after transfection with empty vector (EV) or the indicated fusion-expressing constructs. Per edgeR quasi-likelihood F-test results, *ETV1, ETV4, ETV5*, and *FOXC2* were significantly activated (log_2_ fold change > 1.5, q-value < 0.001) in both the HA-CIC::DUX4 and HA-CIC(ex20)::NUTM1(ex6) conditions vs EV. However, FOXD3, FOXB1, and SOX2 were only significantly activated in the HA-CIC(ex20)::NUTM1(ex6) condition vs EV. In the HA-CIC(ex20::NUTM1(ex6) vs EV comparison, FOXG1 met the significance threshold but was slightly under the log_2_ fold change threshold (value: 1.17), but was included due to its role in brain development. FOXG1 did not meet either significance cutoff in HA-CIC::DUX4 vs EV. (B) Immunoblot of 293T cells approximately 48 hours after transfection with empty vector (EV) or the indicated constructs, representative of the same three independent experiments as in Figure 2A but with a different replicate shown. (C) Immunoblot of 293T cells approximately 48 hours after transfection with empty vector (EV), full length HA-CIC(ex18)::NUTM1(ex3), or truncated versions of HA-CIC(ex18)::NUTM1(ex3) that are comprised only of the HA tag motif and the fractional coding sequence of the indicated partner, split at the breakpoint. Representative of the same three independent experiments as in Figure 2C but with a different replicate shown. (D) Immunoblot of 293T cells approximately 48 hours after transfection with empty vector (EV) or one of three versions of HA-CIC(ex20)::NUTM1(ex6): full length (FL), CIC C1 domain deleted (dC1), or CIC C1 domain and HMG box deleted (dC1 + dHMG). Representative of the same three independent experiments as in Figure 2D but with a different replicate shown.

Of these genes, we validated that FOXB1 protein levels were specifically elevated following CIC::NUTM1 but not CIC::DUX4 expression (Figure 3B). We then revisited our structure-function experiments and observed that upregulation of FOXB1 is observed following expression of FL CIC(ex18)::NUTM1(ex3) (Figure 3C). We also observed that unlike the response of ETV5, FOXB1 protein levels were mildly impaired when the C1 domain was deleted from CIC(ex20)::NUTM1(ex6) and lost upon deletion of both CIC DNA binding domains (Figure 3D). In summary, these data indicate that FOXB1 transcript and protein levels are preferentially elevated by CIC::NUTM1 compared to CIC::DUX4 expression and provide rationale to further explore the role of FOXB1-mediated neurogenic development of CIC::NUTM1 sarcomas.

### Defining the mechanism of CIC::NUTM1-mediated transcriptional activation

Since CIC::DUX4 is thought to gain activating capacity through its interaction with p300 and since NUTM1 is a known p300 interactor^61^, we hypothesized that p300 recruitment was mediated by the addition of NUTM1 to CIC in the context of CIC::NUTM1. Prior studies of *BRD3/4-NUTM1* fusions have localized the minimal region within NUTM1 that binds to p300 to a pair of transactivating domains (TADs) within a roughly 100 amino acid region^39,40^. Importantly, this region is retained in both of the CIC::NUTM1 constructs that we engineered (Figure 1, TAD domain). To test if this region is necessary for the activation of CIC target genes by CIC::NUTM1, we genetically deleted this domain from both of our CIC::NUTM1 constructs (termed dTAD mutants) and evaluated their ability to regulate ETV5 levels. We observed that even following TAD domain deletion, both CIC::NUTM1 versions still displayed moderate activation of ETV5 (Figure 4A). This suggested that while p300 does play a role in the activation of ETV5 by CIC::NUTM1, there may be alternative mechanisms that facilitate a CIC::NUTM1 mediated increase in ETV5 levels.

**Figure 4:**
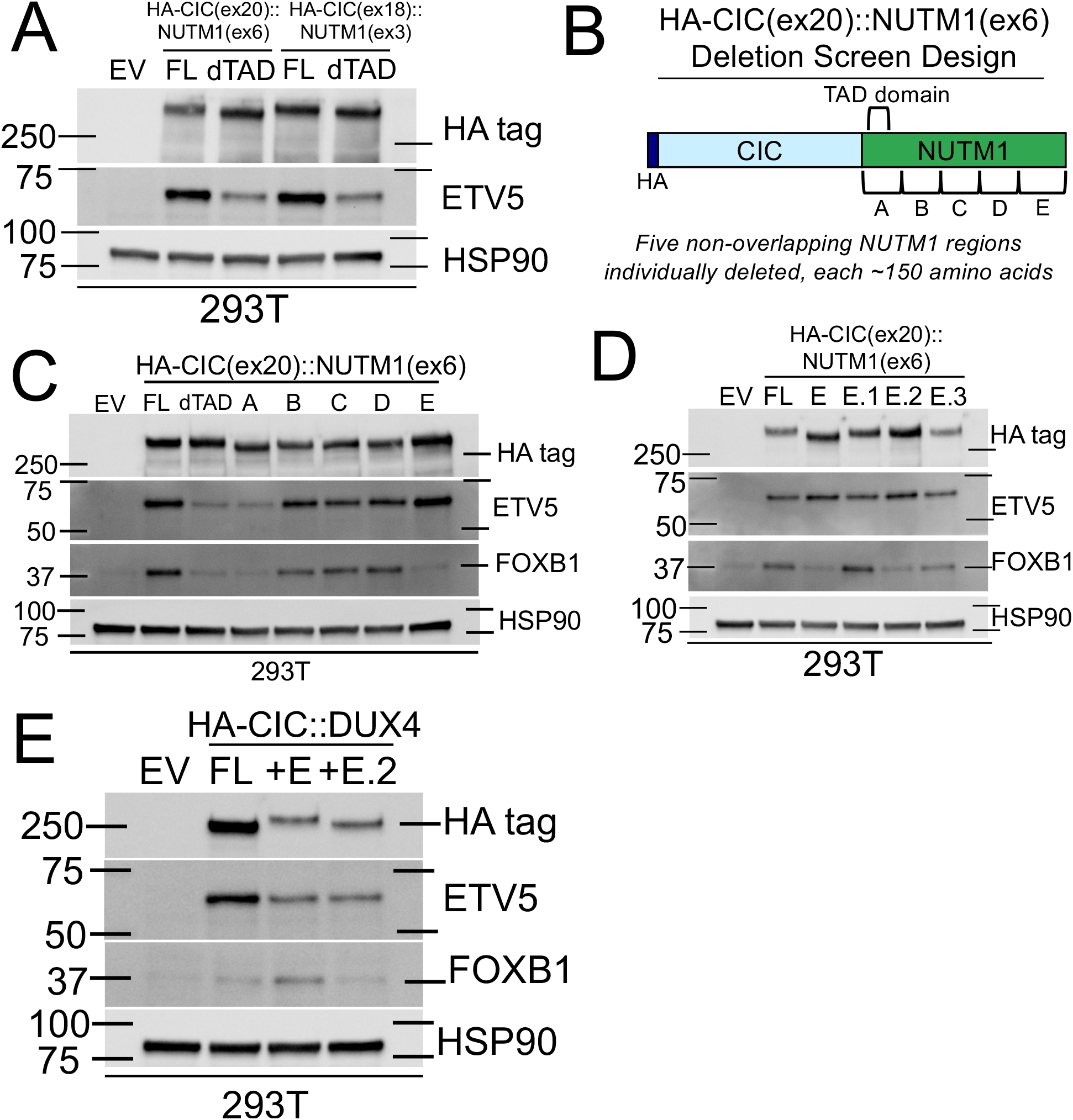
Genetic dissection of the CIC::NUTM1 fusion oncoprotein reveals a key NUTM1 domain that is necessary and sufficient to drive *FOXB1* expression. (A) Immunoblot of 293T cells approximately 48 hours after transfection with empty vector (EV) or one of the indicated versions of HA-CIC(ex20)::NUTM1(ex6) or HA-CIC(ex18)::NUTM1(ex3): full length (FL) or with the p300-interacting domain deleted (dTAD). Representative of three independent experiments. (B) Diagram of the strategy for the CIC(ex20)::NUTM1(ex6) deletion screen. Note that the dTAD mutant has a deletion that is a subset of that for the deletion A mutant. (C) Immunoblot of 293T cells approximately 48 hours after transfection with empty vector (EV) or one of the indicated versions of HA-CIC(ex20)::NUTM1(ex6): full length (FL), dTAD as in panel A, or a deletion of one of the regions depicted in panel B. Representative of two independent experiments. (D) Immunoblot of 293T cells approximately 48 hours after transfection with empty vector (EV) or one of the indicated versions of HA-CIC(ex20)::NUTM1(ex6): full length (FL), the E deletion from panels B and C, or one of three smaller approximately 60 amino acid sub deletions within the E region, described in the Methods section. Representative of three independent experiments. (E) Immunoblot of 293T cells approximately 48 hours after transfection with empty vector (EV) or one of the indicated versions of HA-CIC::DUX4: full length (FL), or with the E region or E.2 region from panels B-D cloned onto the C-terminus of the HA-CIC::DUX4 coding sequence. Representative of three independent experiments.

### A C-terminal domain of NUTM1 is necessary and sufficient to drive FOXB1 activation by CIC(ex20)::NUTM1(ex6) or CIC::DUX4

We considered two explanations for the residual increase in ETV5 levels: first, that the remaining NUTM1 moiety could provide dominant negative activity to de-repress ETV5 in a manner not seen with truncated CIC alone, or second, the possible presence of an unidentified NUTM1 functional domain that contributes to ETV5 activation. We chose to test the second of these hypotheses by undertaking a deletion screen of the NUTM1 fragment of CIC(ex20)::NUTM1(ex6) that deleted five non-overlapping roughly 150 amino acid stretches of the NUTM1 fractional coding sequence (Figure 4B). We expressed these in 293T cells and used expression of both ETV5 and FOXB1 as readouts, where ETV5 serves as a reporter on general CIC-fusion activity and FOXB1 levels indicate CIC::NUTM1-specific activity. While the deletion of region A phenocopied the dTAD mutants (as the TAD region is inside of the A deletion), deletions B, C, and D displayed no change in ETV5 and FOXB1 induction (Figure 4C). However, the deletion of the E region towards the C-terminal end of NUTM1 led to complete abrogation of FOXB1 induction while ETV5 elevation was completely intact (Figure 4C). Deletion of the E region also led to complete loss of FOXB1 induction and a weaker level of ETV5 activation in the CIC(ex18)::NUTM1(ex3) construct (Supplementary Figure 3A). Co-deletion of the E region with the TAD region of NUTM1 did not completely eliminate ETV5 activation by CIC(ex20)::NUTM1(ex6) (Supplementary Figure 3B), indicating this region is not responsible for the residual ETV5 activation observed without the p300-interacting domain of NUTM1.

A further sub-screen of the E region using smaller ∼60 amino acid non-overlapping deletions indicated that the minimal sequence necessary for this phenotype was likely in the middle of the domain (E.2 mutant), possibly with some contribution from the very C-terminal sequence (E.3 mutant) (Figure 4D). To test if this moiety was sufficient to confer FOXB1 activation to CIC::DUX4, which we previously observed not to strongly activate FOXB1, we then cloned the E region or the E.2 subregion onto the C-terminus of CIC::DUX4 and evaluated ETV5 and FOXB1 expression. We observed a moderate increase in FOXB1 activation only by the broader E region (Figure 4E), with the E.2 subregion not sufficient to elevate FOXB1 levels on its own or with a short flexible linker sequence (Figure 4E, Supplementary Figure 3C). These results support the identification of the E region as sufficient to activate FOXB1 by CIC::NUTM1 in a manner uncoupled from the fusion’s ability to activate ETV5 expression.

### The NUTM1 E region permits activation of a gene program beyond core CIC/CIC::DUX4 target genes

To determine if this phenomenon might help to explain the differential regulation of genes beyond *FOXB1* by CIC::NUTM1 vs CIC::DUX4, we performed another bulk RNA-seq experiment in 293T cells expressing several *CIC*-family fusions as well as the dTAD and E-deletion mutants of CIC(ex20)::NUTM1(ex6). Multidimensional scaling analysis reproduced our previous observation that CIC::NUTM1 samples clustered separately from CIC::DUX4 samples (Supplementary Figure 4A). However, we also observed that while the TAD deletion generally seemed to make CIC(ex20)::NUTM1(ex6) samples weaker (i.e. closer to empty vector control), the E-deletion samples still clustered far from control but were now located much closer to the CIC::DUX4 samples (Supplementary Figure 4A).

Heatmap analysis of genes upregulated in the FL CIC(ex20)::NUTM1(ex6) condition revealed two groups of genes: those that were activated regardless of whether or not the E region was present (termed “E ignorers”), and those whose activation was dependent on the presence of the E region (termed “E responders”) (Figures 5A and 5B). We defined lists of these genes (Figure 5B, Supplementary Datasets S1 and S2) and employed gene set enrichment analysis with gProfiler2^62^ (Figure 5C). Among the terms enriched for each group, we noted that those describing general development were significant for both groups, while regulation of ERK1/2 signaling was a hallmark of the E ignorer genes, consistent with prior knowledge that CIC and CIC::DUX4 (which do not have the E region) regulate the MAPK cascade^9,19^. We additionally noted several terms related to CNS development that were specifically enriched for the E responder group, suggesting their activation was associated with the presence of the E region.

**Figure 5:**
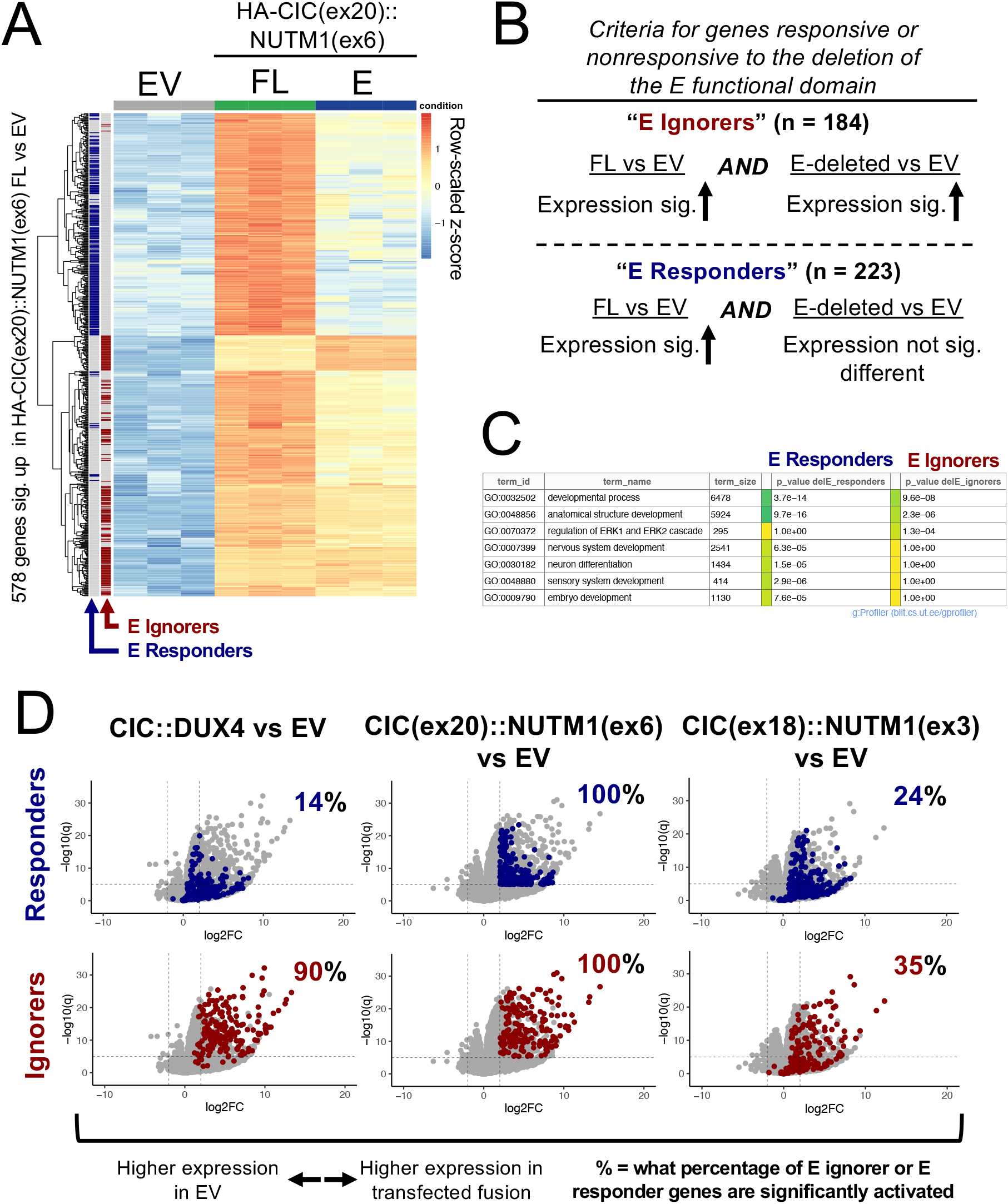
The NUTM1 E region permits activation of a gene program beyond core CIC/CIC::DUX4 target genes. (A) Row-scaled RNA-seq expression heatmap of 578 genes significantly activated (log_2_ fold change > 2, q-value < 0.00001) in the HA-CIC(ex20)::NUTM1(ex6) transfected 293T cells vs EV comparison. Values are shown for cells transfected with EV, full length HA-CIC(ex20)::NUTM1(ex6) (FL), and HA-CIC(ex20)::NUTM1(ex6) with the E region deleted (E). E ignorers and responders as defined in panel B are shaded in on the left. (B) Description of how the E ignorer and E responder gene sets were defined. FL and E-deleted refer to the HA-CIC(ex20)::NUTM1(ex6) full length or E region deletion constructs, respectively. For both gene sets, significant activation in the FL vs empty vector (EV) comparison used cutoffs of log_2_ fold change > 2, q-value < 0.00001. For the E ignorer gene set, significant activation in the E-deleted vs EV comparison was also defined as log_2_ fold change > 2, q-value < 0.00001. For the E responder gene set, no significant change in the E-deleted vs EV comparison was decided using stricter cutoffs of an absolute value log_2_ fold change < 1, and q-value > 0.01. (C) Selected gene set enrichment analysis output using gProfiler2 on the E responder and E ignorer gene sets. (D) Depiction of where the E responder and E ignorer gene sets fall on RNA-seq volcano plots of comparisons between 293T transfected with the indicated fusion-expressing constructs vs. empty vector (EV). The percentages indicate what proportion of the specified gene set meets the thresholds for being significantly activated in a given volcano plot (log_2_ fold change > 2, q-value < 0.00001).

We next mapped the expression of these genes to the CIC(ex18)::NUTM1(ex3) samples (Figure 5D), and observed that while in general fewer of them met the significance threshold for activation (consistent with an overall weaker fusion compared to CIC(ex20)::NUTM1(ex6) as seen in Supplementary Figure 4A), a similar amount of both E ignorers (35%) and E responders (24%) were significantly activated in these samples. However, in the CIC::DUX4 samples, 90% of the E ignorer genes were significantly activated while just 14% of E responder genes met the same criteria (Figure 5D). This suggested that the E ignorer genes are likely a hallmark of both CIC::DUX4 and CIC::NUTM1 gene signatures, while the E responder genes are specifically biased towards CIC::NUTM1 expression due to the NUTM1 E region. Comparison of the dTAD mutants with the E-deletion mutants of CIC(ex20)::NUTM1(ex6) further demonstrated that TAD deletion generally led to a reduction of target gene expression in general, while loss of the E region retained high expression levels of many genes including those known to be CIC and CIC::DUX4 targets (Supplementary Figure 4 B-D). Together, these results support the notion that the NUTM1 E region enables CIC::NUTM1 to activate a gene program separate from core CIC and CIC::DUX4 target genes, which may begin to explain the divergent behavior between CIC::NUTM1 and CIC::DUX4 bearing tumors.

### CIC::LEUTX is a relatively weak activator of CIC target genes in a manner dependent on its two C-terminal transactivation sequences

We next turned to CIC::LEUTX, which in our synthetic patient-informed construct is comprised of almost the entire *CIC* coding sequence with a small *LEUTX* fragment at the C-terminus, similar to *CIC::DUX4* (Figure 1). Expression of this construct led to a milder ETV5 activation compared to the response seen from CIC::DUX4 or CIC::NUTM1, with no accompanying increase in FOXB1 expression (Figure 6A, Supplemental Figure 5A). RNA-seq data similarly showed that CIC::LEUTX was relatively weak compared to the other FL fusions (i.e. clustered closer to the empty vector control samples, Supplementary Figure 4A).

**Figure 6:**
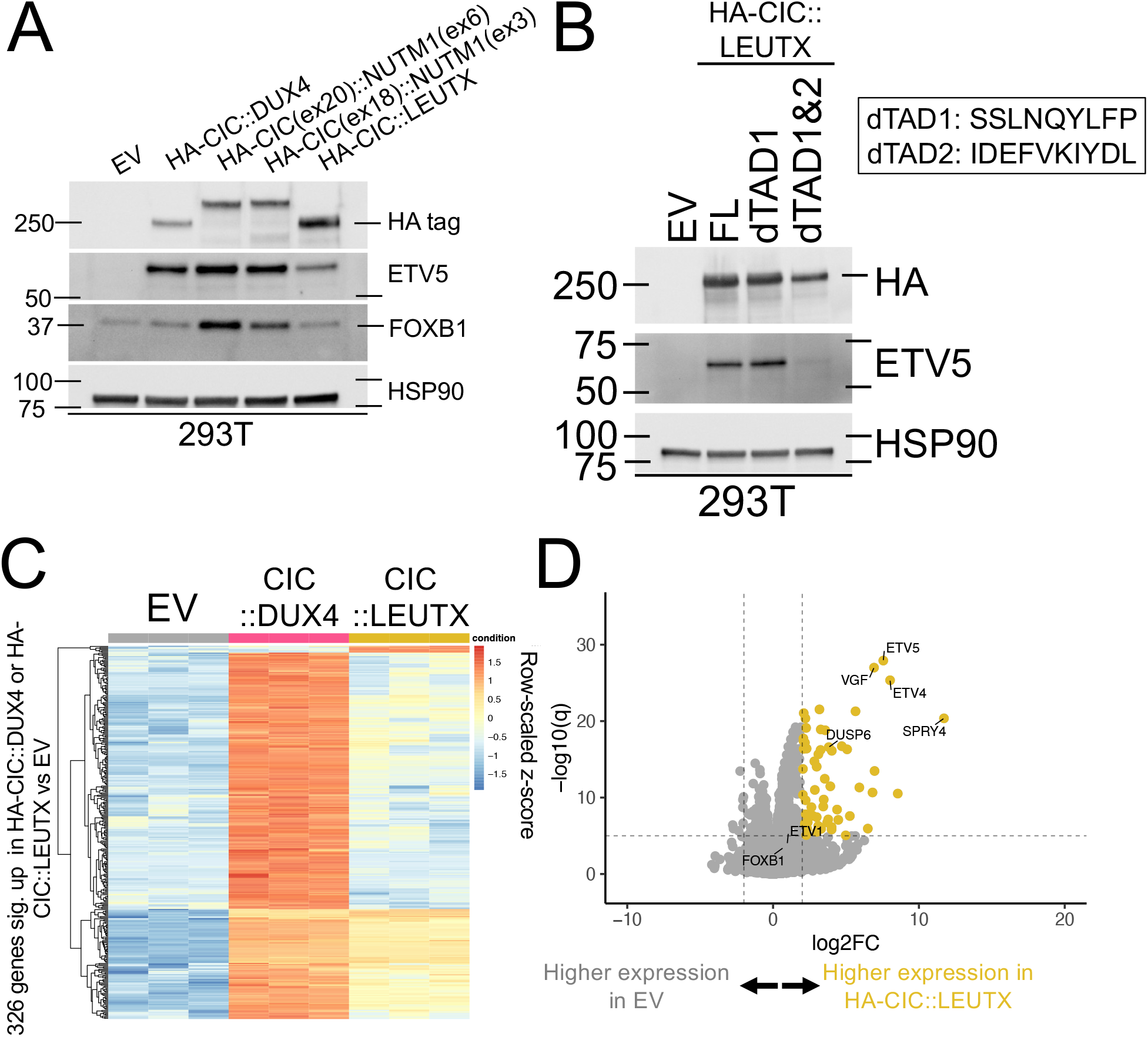
CIC::LEUTX is a relatively weak activator of CIC target genes in a manner dependent on its two C-terminal transactivation sequences. (A) Immunoblot of 293T cells approximately 48 hours after transfection with empty vector (EV) or constructs expressing the indicated fusion oncoproteins. Experiment was performed once, but replicated in a similar experiment in Supplementary Figure 5. (B) Immunoblot of 293T cells 48 hours after transfection with empty vector (EV) or constructs expressing the following HA-CIC::LEUTX sequences: full length (FL), dTAD1, and dTAD1&2, where the dTAD mutants have deletion of the indicated LEUTX C-terminal amino acids. Representative of three independent experiments. (C) Row-scaled RNA-seq expression heatmap of 326 genes significantly activated (log_2_ fold change > 2, q-value < 0.00001) in at least one of the HA-CIC::DUX4 or HA-CIC::LEUTX transfected 293T cells vs EV comparisons. (D) Volcano plot of RNA-seq expression data from 293T cells 48 hours after transfection with empty vector (EV) or a construct encoding HA-CIC::LEUTX. Select CIC/CIC::DUX4/CIC::NUTM1 target genes are indicated.

WT LEUTX has been shown to interact with p300/CBP through at least one of two short transactivating motifs towards its C-terminus^38,63^, both of which are retained in our construct. We deleted one or both of these motifs and tested the mutants for their ability to activate ETV5, which indicated that loss of the SSLNQYLFP motif alone (previously suggested to abrogate the p300/CBP-LEUTX interaction^38^) is not sufficient to block ETV5 induction (Figure 6B). However, deletion of both motifs almost entirely eliminated ETV5 induction, suggesting that CIC::LEUTX does indeed activate ETV5 in a manner dependent on these motifs and the recruitment of p300/CBP.

We compared the transcriptional activation signature driven by CIC::LEUTX to that generated by CIC::DUX4 using our RNA-seq data and noted that CIC::LEUTX drove the activation of a smaller set of genes than CIC::DUX4, but still significantly increased the expression of several core CIC target genes (Figure 6C-D). Taken together, these data indicate that our CIC::LEUTX fusion operates as an attenuated version of CIC::DUX4 that is capable of activating core CIC target genes through two C-terminal transactivation motifs, likely in a p300/CBP-dependent manner.

### ATXN1::DUX4 activates the expression of select CIC target genes in a manner dependent on the AXH domain

Finally, we aimed to investigate *ATXN1::DUX4*, which is interesting because it is thought to indirectly activate CIC target genes due to the physical interaction between ATXN1 and CIC, which is mediated by the ATXN1 AXH domain^43^. If proven true, *ATXN1/ATXN1L* fusions would be the first example of fusions that drive tumors with characteristics of *CIC*-rearrangements without directly altering the *CIC* locus. Notably, the construct that we generated was based on a breakpoint that does retain the ATXN1 AXH domain (Figure 1), making this mechanism testable with our model. We expressed both the FL ATXN1::DUX4 and a mutant lacking the AXH domain and observed that ETV5 levels were increased only in the context of the FL fusion (Figure 7A).

**Figure 7:**
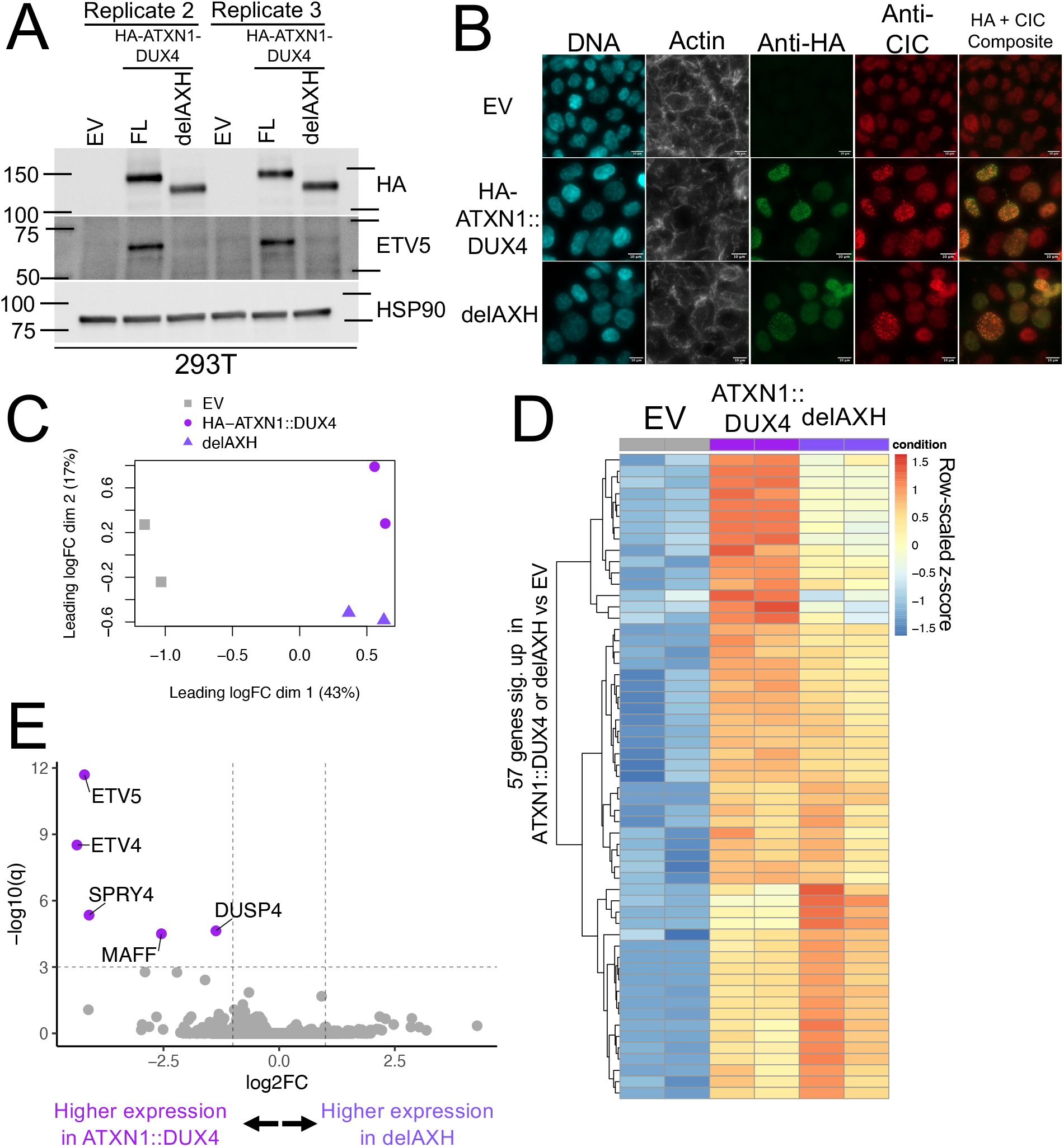
ATXN1::DUX4 activates select CIC target genes in an AXH domain-dependent manner. (A) Immunoblot of 293T cells approximately 48 hours after transfection with empty vector (EV) or constructs expressing the full length (FL) or AXH domain deleted (delAXH) versions of HA-ATXN1::DUX4. Two representative replicates of three independent experiments are shown. (B) Immunofluorescence of 293T cells approximately 48 hours after transfection with empty vector (EV) or the indicated constructs and stained with DAPI (DNA), rhodamine-phalloidin (Actin), an anti-HA antibody, or an anti-CIC antibody. Scale bars indicate 10 μm, representative cells are shown from one of two experiments. (C) edgeR multi-dimensional scaling plot of RNA-seq data from 293T cells 48 hours after transfection with empty vector (EV) or the indicated fusion-expressing constructs. (D) Row-scaled RNA-seq expression heatmap of 57 genes significantly activated (log_2_ fold change > 1, q-value < 0.001) in at least one of the indicated comparisons. Values are shown for cells transfected with EV, HA-ATXN1::DUX4, and AXH-deleted HA-ATXN1::DUX4 (delAXH). (E) Volcano plot of RNA-seq expression data from 293T cells approximately 48 hours after transfection with the indicated constructs.

To test if the ATXN1::DUX4 fusion impacts endogenous CIC localization we performed immunofluorescence microscopy on 293T cells transfected with the full length or AXH-deleted versions of ATXN1::DUX4. Interestingly, regardless of the presence of the AXH domain we observed some cells with puncta of ATXN1::DUX4 that co-localized with CIC, whereas in cells not expressing the fusion, CIC was diffuse in the nucleus (Figure 7B). However, we also noted that some transfected cells clearly expressed ATXN1::DUX4 but did not show these puncta, again regardless of the presence or absence of the AXH domain. Consequently, it is difficult to determine if these puncta are biologically relevant or technical artifacts. However, we have not observed puncta formation when using similar constructs to overexpress CIC::DUX4 or CIC::NUTM1 in our current (Figure 2B) or in our prior studies^18^, suggesting this is a phenomenon specific to the ATXN1::DUX4 fusion.

Regardless of its impact on co-localization with CIC, deletion of the AXH domain clearly impacted ETV5 activation (Figure 7A), which led us to perform RNA-seq on 293T cells expressing the full length and AXH-deleted ATXN1::DUX4 constructs. We observed that samples generally segregated by whether ATXN1::DUX4 was present or not, with separation on the second dimension based on the presence or absence of the AXH domain (Figure 7C). Heatmap analysis showed surprisingly few genes whose activation was significantly lost when the AXH domain was deleted (Figure 7D), and a volcano plot revealed that all five of these genes are known or likely CIC target genes^10,64^ (Figure 7E). We also noticed activation of *ZSCAN4* following expression of either form of ATXN1::DUX4, which is a known DUX4 target gene^65^ and likely due to the fact that in this particular fusion the DUX4 breakpoint is positioned early in the coding sequence allowing for retention of the second of two DUX4 DNA-binding homeobox domains (Supplementary Figure 6A-C). This data strongly suggests that the ATXN1::DUX4 fusion does indeed activate select CIC target genes in a manner dependent on the AXH domain.

## Discussion

Of the plethora of fusion oncogene-driven cancers that arise in patients, the majority are rare entities that will only affect a small number of individuals each year. Within these families of rare tumors, it can be even more difficult to identify patients harboring variants of promiscuous fusions with varying partner genes that alter their activity. While learning about the minute differences between fusions with differing partner gene composition can be biologically and clinically informative, the scarcity of patient samples makes them difficult to study through the traditional approaches of cloning from tumor tissue. In this study, we used an alternative approach of cloning synthetic, patient-informed fusions designed from real breakpoint sequences to study convergent and divergent biology among *CIC*-family fusion oncogenes.

While limited clinical and transcriptional data has suggested that CIC::NUTM1 may somewhat diverge in its functional capabilities from CIC::DUX4, we provide here the first evidence that they do differ in functional domain requirements and transcriptional programs. We followed up on our prior observation that CIC::NUTM1 breakpoints often exclude the C1 domain^18^ to show that these fusions indeed seem less reliant on the C1 domain to activate ETV5, a major difference from CIC::DUX4. We then localized a poorly defined 150 amino acid C-terminal domain of NUTM1 that through an as of yet unknown mechanism confers the ability of NUTM1 to activate a gene program including *FOXB1*. This program is largely not driven by CIC::DUX4 and appears to be enriched for genes involved in neural development, suggesting the possibility that this neomorphic behavior may in part explain why CIC::NUTM1 tumors have preferential tropism for the CNS and spine, though this is beyond the scope of our study. We anticipate that future studies will be able leverage these newly engineered molecular tools to further understand how FOXB1 or its broader gene program may mediate CIC::NUTM1 tumor development.

The E region falls on a part of the NUTM1 protein where few annotated functional domains exist. It appears to contain both putative nuclear localization and nuclear export sequences^66^, and closely abuts a defined but unstudied “TAD2” region^67^. It also overlaps a Uniprot annotation for a long disordered region at the C-terminus of NUTM1 (Uniprot Q86Y26). We noted that in addition to its effects on the transcriptional program exerted by CIC::NUTM1, the deletion of the E region appeared to consistently increase fusion protein expression (Figure 4C-D, Supplementary Figure 3A-B), and its addition to CIC::DUX4 decreased expression of those fusions (Figure 4E, Supplementary Figure 3C). We tend to be conservative about making conclusions on protein stability using plasmid models because our system does not precisely control expression levels, but the reproducibility of this phenomenon across several mutants and fusion constructs suggests the E region may impact protein stability or expression.

We think at least one of three explanations is likely to explain how the E region allows for the activation of new target genes in CIC::NUTM1. The first is that it may alter the DNA binding properties of CIC::NUTM1 compared to those of CIC::DUX4, allowing fusion binding at new sites and activation of nearby genes. The second is that the E region may amplify overall transactivation by CIC::NUTM1, increasing the expression of genes that are normally only very weakly regulated by CIC::DUX4. The third is that it may recruit different protein complexes to the fusion and drive additional chromatin remodeling separate from p300/CBP activity, allowing for the induction of new genes not activated by CIC::DUX4. Regardless of the mechanism, perhaps the most interesting future questions will be whether or not the gene program driven by the E region is reproducible in more patient-relevant models and is sufficient to explain the CNS and spine tropism observed in CIC::NUTM1 tumors. This is an important question which requires improvement and refinement of our current models and the discovery of the cell-of-origin for these tumors to dissect.

Given the prevailing hypothesis for how *CIC*-fusions mechanistically drive transcription, we also took the opportunity to study the role of p300/CBP in activation of CIC target genes using our *CIC::NUTM1* and *CIC::LEUTX* constructs. While we found that CIC::LEUTX does appear to be dependent on its two C-terminal LEUTX transactivation domains to activate ETV5, likely through recruitment of p300/CBP, we observed a substantial increase in known CIC target genes by CIC::NUTM1 even when its putative p300-recruiting domain was deleted. The mechanism explaining this behavior in CIC::NUTM1 is not immediately clear, as it is not explained by the E region and a truncating CIC mutation did not result in the same level of ETV5 activation. One potential explanation is that the addition of a bulky NUTM1 moiety, even if incapable of recruiting p300/CBP, is sufficient to generate a stronger dominant-negative mutant and increase CIC target gene expression. Deciphering this mechanism is worth pursuing because recent work has nominated using p300/CBP inhibitors and degraders in CIC::DUX4-bearing tumors to block target gene activation^21,68^. While our data suggests this approach may also be useful in blocking CIC::LEUTX-mediated signaling, it might not prove efficacious in CIC::NUTM1-bearing tumors which may operate through a p300/CBP independent manner to activate CIC targets.

The recent rise in reports of tumors bearing fusions of *ATXN1* or *ATXN1L* with 3’ partner genes including *DUX4, NUTM1*, and *NUTM2A* has been of special interest because ATXN1 and ATXN1L are known to functionally interact with CIC. The observation that *ATXN1*/*ATXN1L*-fused tumors tend to cluster with CIC-rearranged tumors by methylation profiling has raised the idea that these fusions likely activate CIC target genes by colocalizing a p300-interacting domain to CIC target sites.^43^ Our model provides data that suggests ATXN1::DUX4 is indeed capable of activating certain CIC target genes in a manner dependent on the AXH domain, consistent with a model of indirect CIC-target gene activation. Additionally, our results suggest that ATXN1::DUX4 may be able to form nuclear puncta which include CIC, but the mechanism and biological relevance remain unclear. Future studies are necessary to confirm that activation of CIC target genes is required to drive oncogenesis in ATXN1-rearranged tumors since it remains possible that other as of yet undefined functions of ATXN1::DUX4 underly tumorigenesis.

We chose to generate synthetic, patient-informed constructs of understudied CIC-family fusions to enable rapid structure-function studies and to overcome a barrier for studying these ultra-rare tumor types. While their synthetic nature and our choice to use them in 293T cells make them far from perfect models, they faithfully recapitulate known transcriptional signatures and are easily mutated to investigate functional domain requirements. In this study, we used these tools to identify a new functional domain of NUTM1 that influences CIC::NUTM1 activity, characterize CIC::LEUTX as a weak p300-dependent activator of CIC target genes, and provide evidence for the indirect-fusion theory of CIC target gene activation by ATXN1::DUX4. We view these tools as first generation constructs that allow us as a community to start exploring fusion promiscuity in CIC-rearranged tumors. We thus fully intend to distribute these constructs as a key resource (Addgene) and invite the scientific community to partake in future studies that utilize this *CIC* fusion molecular tool kit. We hope that these constructs will stimulate active investigation into the basic mechanisms that underly these understudied *CIC*-family fusion members and provide a foundation for the development of future CIC-rearranged model systems.

### Limitations of the Study

The primary limitation of this work is that our experiments were conducted in 293T cells, which are likely not particularly biologically relevant to patient-derived tumors. However, we consistently observed the proper regulation of key transcriptional responses matching those in tumors. Moreover, we largely performed structure-function experiments, which are unlikely to be cell type dependent. The main area where caution should be applied is in extending our results to explain tumorigenesis, for example whether FOXB1 and the E region may impact the anatomical location of CIC::NUTM1 tumors. While we can hypothesize about such a relationship, our data do not permit any conclusions of this nature and answering such questions will require more robust models.

While our constructs are based on breakpoints derived from actual patients, they are not cloned directly out of patient tumors and thus are potentially subject to caveats based on the sequences used to clone them. To be transparent, we have included complete details about the sequences used for cloning and what (if any) mutations they have compared to reference partner gene sequences in the Methods section.

Finally, our approach of transiently transfecting plasmids into 293T cells and evaluating response after a short period of time (typically 48 hours) enables rapid experimentation but has several limitations. First, despite being driven by similar promoter-enhancer elements, our constructs do not tightly regulate expression levels and thus there may be some impacts on readouts like target gene activation that could be dependent on transgene expression levels. Where relevant, we have tried to only draw conclusions that seem not to be affected by this (e.g. CIC::LEUTX expresses at higher protein levels than other CIC-fusions, but yields lower activation of ETV5) or we designed experiments such that mutants were compared to FL constructs of the same fusion. However, caution should be taken when directly comparing readouts from different fusions to each other. Second, the short timeframe at which we typically assayed for results could mean that some of our outputs (particularly gene programs) could be relevant at short timescales but not longer ones. For example, we did not find *FOXB1, FOXD3*, or *FOXG1* as differentially expressed genes for *CIC*-rearranged tumors in the only patient-derived transcriptomic dataset including *CIC::NUTM1* (Watson et al. 2018^23^, see their Supporting Information Table S3). This could imply a limitation of using 293T cells, or it could suggest that these genes play a transient role in tumorigenesis. Addressing this caveat will require the development of stable, longer-term models that sustain expression of these fusions over long time periods and permit time-based study of tumor development. Third, a related issue is that we use a strong CMV promoter-enhancer element to drive expression of these fusions, which may not directly relate to endogenous expression levels observed in tumors. Thus, future studies are required in order refine our initial interpretations and overcome these limitations.

## Supporting information

Supplementary Figures and Supplementary Dataset Descriptions

## Lead Contact

Requests for further information and resources should be directed to and will be fulfilled by the lead contact, Ross Okimoto.

## Materials Availability

The pCMV3 empty vector and the full-length pCMV3 expression constructs of HA-CIC(ex20)::NUTM1(ex6), HA-CIC(ex18)::NUTM1(ex3), HA-CIC::LEUTX, and HA-ATXN1::DUX4 are in the process of being deposited with Addgene as plasmids number 238135 through 238139 and will be available upon request until the deposit is finalized. Other constructs described in this paper are available upon request.

## Data and Code Availability

- RNA-seq data have been deposited at GEO: GSE295623, GEO: GSE295624, and GEO: GSE295625 and are publicly available as of the date of publication.
- Uncropped western blot images and Ponceau S loading control images are available in Supplemental Dataset S3.
- All original code is available at https://github.com/cuylerluck/CICfamily_models and is publicly available as of the time of publication.
- Any additional information required to reanalyze the data reported in this paper is available from the lead contact upon request.

## Acknowledgements

C. Luck acknowledges funding from the University of California, San Francisco BMS Graduate Program Training Grant T32GM136547-01 (National Institute of General Medical Sciences), the University of California, San Francisco Discovery Fellows Program, and an NCI F31 (CA287493). The content is solely the responsibility of the authors and does not necessarily represent the official views of the National Institutes of Health. K.A. Jacobs acknowledges funding from Tobacco-Related Disease Research Program Predoctoral Fellowship T33DT6442. This material is based upon work supported by the National Science Foundation Graduate Research Fellowship Program under grant no. 2038436 (to K.A. Jacobs). Any opinions, findings, and conclusions or recommendations expressed in this material are those of the author(s) and do not necessarily reflect the views of the National Science Foundation. R.A. Okimoto acknowledges funding from an NCI grant (R37 CA255453), the Children’s Cancer Research Fund, and Cookies for Kids’ Cancer. We want to thank Takuro Nakamura, Chris French, Huda Zoghbi, and Stephen Tapscott for sharing their plasmids either directly or through Addgene. C. Luck wishes to extend special thanks to Kevin Shannon, Alejandro Sweet-Cordero, and Dave Toczyski for their support and input during the development of this project. We are particularly grateful to the patients in the original studies that guided design of our constructs, who chose to donate their samples for scientific research, and the authors of those studies for sharing their data via their publications.

## Author Contributions

Conceptualization, C.L., R.A.O.; Data curation, C.L.; Formal analysis, C.L.; Funding acquisition, C.L., R.A.O.; Investigation, C.L., K.A.J., J.R., C.D.M., R.K.M.P.; Methodology, C.L.; Project administration, C.L., R.A.O.; Resources, C.L., K.A.J.; Software, C.L.; Supervision, C.L., R.A.O.; Validation, C.L.; Visualization, C.L.; Writing – original draft, C.L., R.A.O.; Writing – review & editing, C.L., K.A.J., J.R., C.D.M., R.K.M.P., R.A.O.

## Methods Cell Culture

293T cells (female) were purchased from ATCC (CRL-3216, RRID: CVCL_0063) in September 2021 and secondarily short tandem repeat verified with the FTA Human (135-XV) Authentication Service (ATCC). Cells were managed as described previously^18^. Briefly, cell lines were grown in humidified incubators at 37°C with 5% CO_2_ in DMEM with high glucose, L-glutamine, and sodium pyruvate (SH30243.02, Cytiva) supplemented with 10% FBS, 100 U/mL penicillin, and 100 ug/mL streptomycin. Cells were routinely screened for *Mycoplasma* using the e-Myco plus Mycoplasma PCR Detection Kit (Boca Scientific); for any given thawed cell line this typically included testing after use for the final experiment prior to disposal. Thawed cells were allowed to recover for approximately 48 hours prior to use in experiments. Cells were typically grown for no longer than one month before being replaced with a lower-passage vial.

### Cloning

The pcDNA3.1-HA-CIC::DUX4 plasmid was a gift from Takuro Nakamura^16^. All primers used for cloning are described in Supplementary Dataset S4. The pCMV3 empty plasmid, which served as a backbone for the cloned CIC- and ATXN1-synthetic fusion constructs, was derived from Sino Biologicals AG13105-CF by replacing the GFP-FLAG insert with a START-STOP motif via deletion PCR with the Q5 Site-Directed Mutagenesis Kit (NEB E0554). NEB 10-beta *E. coli* cells (NEB C3019) were used for most cloning, as in our hands they gave better efficiencies when cloning large plasmids. Full plasmid sequencing was performed by Plasmidsaurus using Oxford Nanopore Technology with custom analysis and annotation. Sanger sequencing was performed by Quintara Biosciences.

The synthetic fusions were modeled after the following patient reports (identifiers & references given): CIC(ex20)::NUTM1(ex6) – RNA012_16_073^23^; CIC(ex18)::NUTM1(ex3) – SARC084 / RNA012_16_074^23^; CIC::LEUTX – AS1^30^; ATXN1::DUX4 – the patient described in Satomi et al 2022^43^. The synthetic fusions were cloned with N-terminal HA tags into the pCMV3 empty backbone using NEB HiFi (NEB E2621), and an overview of the strategies is described in Supplementary Figure 2. The two CIC::NUTM1 fusions are comprised of CIC sequences from the pcDNA3.1-HA-CIC::DUX4 plasmid and NUTM1 sequences from pcDNA5 frt/to N-bioTAP-C-BRD4-NUT, a gift from Chris French (Addgene #171630, RRID:Addgene_171630)^69^. The CIC::LEUTX fusion is comprised of a CIC sequence from Origene RC215209 (contains CIC [NM_015125] C-terminally Myc-DDK tagged in a pCMV6 backbone) and a LEUTX sequence synthesized by Twist Bioscience that was derived from NM_001382345.1. The ATXN1::DUX4 fusion is comprised of an ATXN1 sequence from Flag-ATXN1, a gift from Huda Zoghbi (Addgene #48189, RRID:Addgene_48189)^70^ and a DUX4 sequence from pCW57.1-DUX4-WT, a gift from Stephen Tapscott (Addgene #99282, RRID:Addgene_99282)^71^.

The pCMV3 empty control vector and the four synthetic fusion pCMV3 expression plasmids will be available from Addgene (plasmids #238135 through #238139), including annotated full-plasmid sequencing files, when the deposit is finalized. Until then, these and all mutant constructs plus some undescribed lentiviral/retroviral constructs are available upon request.

Because the synthetic fusions were cloned from various sources, the sequences of their partner genes do not always exactly match the respective reference transcript sequences. This is also true of the CIC::DUX4 coding sequence, as it was originally cloned from a patient tumor. The amino acid differences are described below as compared to reference sequences for CIC (NM_015125.5), DUX4 (NM_001306068.3), NUTM1 (NM_001284292.2), LEUTX (NM_001382345.1), and ATXN1 (NM_001128164.2):

1. CIC::DUX4 – CIC fragment has a silent mutation at C270 and a micro deletion of the reference G1443-T1444.
2. CIC(ex20)::NUTM1(ex6) – CIC fragment has a silent mutation at C270 and a micro deletion of the reference G1443-T1444. NUTM1 fragment has a silent mutation at G867.
3. CIC(ex18)::NUTM1(ex3) - CIC fragment has a silent mutation at C270. NUTM1 fragment has a P50L mutation, a silent mutation at D103, a series of silent mutations involving A366-K368 and R370-P372, and a silent mutation at G867.
4. CIC::LEUTX – no differences from reference sequences.
5. ATXN1::DUX4 – ATXN1 fragment has a silent mutation at V118, only two Q residues in the CAG trinucleotide repeat region (reference Q197-Q225), and a silent mutation at L233.

Since the CIC::LEUTX fusion was cloned with a different CIC donor than that used for CIC::NUTM1 (which derived their CIC sequences from CIC::DUX4), there are minor differences in the CIC sequences between the two, of which the only coding difference is that CIC::LEUTX does not have deletion of G1443-T1444. Please also note that the pcDNA3.1-HA-CIC::DUX4 plasmid backbone has minor differences from the pCMV3 backbone used for the other fusions, specifically that it has AmpR instead of KanR, NeoR instead of HygR, and a slightly different CMV promoter-enhancer element. The pCMV3 backbone also possesses a chimeric intron element between the CMV promoter and the start of the coding sequences which the pcDNA3.1 vector does not. Despite these differences, we routinely observed comparable expression of all fusions regardless of their backbone or their mutations, though we cannot rule out plasmid-intrinsic effects. The empty vector control for most experiments was the empty pCMV3 construct, but was a previously used backbone^18^ more similar to the pcDNA3.1 backbone for some experiments only involving CIC::DUX4.

All deletion mutants were made with the Q5 Site-Directed Mutagenesis Kit (NEB E0554). The HA-CIC::DUX4 +E and +E.2 mutants were made with NEB HiFi (NEB E2621). The HA-CIC::DUX4 +linker+E.2 construct was made with the Q5 Site-Directed Mutagenesis Kit (NEB E0554), with a (GGSG)x3 linker whose sequence was derived from iGEM, part number BBa_K243006.

The amino acid coordinates for specific mutants and functional domains that were mutated are listed below, with the syntax of… (mutant/domain name/description): [first_residue]-[last_residue], (plasmid), {notes on the source of domain annotation or mutant identity}

… where the number of the residues refers to their position in the fusion coding sequence described in the given plasmid sequence, where the first residue is the initiating methionine at the start of the 3x HA-linker motif. Full plasmid sequences for the expression vectors of the full-length synthetic fusions will be provided on Addgene or on request until the Addgene deposit is finalized as described above. Where applicable, the source of the domain annotation is given.

1. HA-CIC (truncated): M1-T1505, pCMV3-HA-CIC(ex18)::NUTM1(ex3), i.e. removes the NUTM1 fragment
2. HA-NUTM1 (truncated): M1-A63 + A1506-Q2632, pCMV3-HA-CIC(ex18)::NUTM1(ex3), i.e. removes the CIC fragment
3. CIC C1 domain: R1525-M1580, pCMV3-HA-CIC(ex20)::NUTM1(ex6), sequence originally from Forés et al 2017^53^
4. CIC HMG box: I263-K331, pCMV3-HA-CIC(ex20)::NUTM1(ex6), sequence from UniProt (RRID: SCR_002380) entry Q96RK0
5. NUTM1 TAD region: W1645-R1738, pCMV3-HA-CIC(ex20)::NUTM1(ex6), sequence from Yu et al 2023^39^ (the ΔF1c mutant)
6. NUTM1 A region: P1625-D1771, pCMV3-HA-CIC(ex20)::NUTM1(ex6)
7. NUTM1 B region: G1772-K1921, pCMV3-HA-CIC(ex20)::NUTM1(ex6)
8. NUTM1 C region: Q1922-C2071, pCMV3-HA-CIC(ex20)::NUTM1(ex6)
9. NUTM1 D region: S2072-Q2221, pCMV3-HA-CIC(ex20)::NUTM1(ex6)
10. NUTM1 E region: E2222-Q2400, pCMV3-HA-CIC(ex20)::NUTM1(ex6)
11. NUTM1 E.1 region: E2222-E2281, pCMV3-HA-CIC(ex20)::NUTM1(ex6)
12. NUTM1 E.2 region: D2282-G2341, pCMV3-HA-CIC(ex20)::NUTM1(ex6)
13. NUTM1 E.3 region: K2342-Q2400, pCMV3-HA-CIC(ex20)::NUTM1(ex6)
14. LEUTX TAD1: S1743-P1751, pCMV3-HA-CIC::LEUTX, sequence from Gawriyski et al 2023^38^
15. LEUTX TAD2: I1727-L1736, pCMV3-HA-CIC::LEUTX, sequence from Katayama et al 2018^63^
16. ATXN1 AXH domain: S598-G729, pCMV3-HA-ATXN1::DUX4, sequence from Uniprot entry P54253

### Plasmid Transfections

Typically, 1.5 μg of each plasmid was reverse transfected into either 300,000 or 500,000 293T cells (depending on the experiment) in one well of a 6-well plate. Plasmid mixes were prepared with FuGENE 6 (Promega) at a 2:1 FuGENE:DNA ratio in Opti-MEM (Gibco). Transfected cells were usually used for downstream experiments approximately 48 hours after transfection.

### Western Blot

Western blots were performed in line with previous work^18^. At the time of protein harvest, adherent cell-containing plates were placed on ice, media was aspirated, and cells were gently washed 3 times with ice-cold Dulbecco’s Phosphate-Buffered Saline (DPBS). RIPA buffer supplemented with Halt protease and phosphatase inhibitors (Thermo Fisher Scientific) was then added, and cells were mechanically disrupted with cell scrapers. Cell suspensions were then incubated on ice for at least 15 minutes before sonication and centrifugation, after which the supernatant was stored at −20°C or −80°C until analysis.

Protein lysates were quantitated with the Pierce BCA kit (Thermo Fisher Scientific), normalized, boiled, and separated by denaturing electrophoresis on Criterion TGX 4-15% gels (Bio-Rad), with Precision Plus Protein Dual Color Standards (Bio-Rad) used as ladders. Proteins were transferred to nitrocellulose membranes by the Trans-Blot Turbo system (Bio-Rad), evaluated for transfer by Ponceau S staining (Sigma-Aldrich), and blocked in 5% Bovine Serum Albumin (BSA) in Tris Buffered Saline with 0.1% Tween 20 (TBS-T) for at least 1 hour. Membranes were cut horizontally for each target of interest and incubated in primary antibody diluted in blocking buffer on a cold room rotator overnight. Separate blots from identically loaded samples run at the same time were used to visualize targets of similar sizes. These and other per-blot details and full Ponceau S loading controls are shown in Supplementary Dataset S3. The next day, blots were washed 3x for 5 minutes in TBS-T, incubated in secondary antibody diluted in blocking buffer for 1 hour on a room temperature rotator, and again washed 3x for 5 minutes in TBS-T. Blots were imaged on a Bio-Rad ChemiDoc Touch using ECL Prime reagent (Millipore Sigma) by briefly drying the blot, submerging the blot in ECL Prime mixture, dabbing excess solution off, and imaging. When required, brightness/contrast was adjusted either on the ChemiDoc or using Image Lab (Bio-Rad). Please note that while HSP90 is shown for most blots as a presence/absence loading control, it was not optimized for quantitation. For better analysis of quantitation between lanes, please refer to the matched Ponceau S stains in Supplementary Dataset S3.

Antibodies and dilutions or concentrations used were: anti-HA clone C29F4 (Cell Signaling Technology 3724, 1:1000-1:5000), anti-HSP90 (Cell Signaling Technology 4874, 1:1000), anti-ETV5 clone E5G9V (Cell Signaling Technology 16274, 1:1000), anti-FOXB1 (Invitrogen PA5-28134, 1:1000), anti-SOX2 (Cell Signaling Technology 2748, 1:1000), anti-FOXC2 (Proteintech 23066-1-AP, 1:1000), anti-FOXD3 (Abcam ab64807, 1:1000), anti-FOXG1 (Abcam ab18259, 1 ug/mL), and horseradish peroxidase-linked anti-Rabbit IgG (Cell Signaling Technology 7074, 1:3000).

### Immunofluorescence

Immunofluorescence was performed similarly to that in previous work^18^. 293T cells were transfected as described above with the appropriate constructs. The next day, transfected cells were transferred onto poly-L-lysine (0.01%, Sigma-Aldrich P4707-50mL) coated coverslips and allowed to adhere overnight. Then, coverslips were briefly and gently rinsed once with DPBS, fixed with 4% paraformaldehyde in PBS at 37°C for 10 minutes, washed 1x with DPBS, quenched with 100 mM glycine at RT for 30 minutes, washed 2x with DPBS, and permeabilized with 0.2% Triton X-100 for 15 minutes at RT. Coverslips were than washed 3x for 5 minutes with DPBS and blocked for one hour in 2% BSA in DPBS for one hour at RT. Next, coverslips were inverted onto solutions of primary antibody diluted in blocking buffer for 1-2 hours at RT and were washed 3x for 5 minutes with DPBS. Coverslips were then again inverted onto solutions containing secondary antibody, DAPI (Invitrogen D1306, 1-5 mg/mL, diluted 1:1000-1:2000), and Rhodamine-Phalloidin (Invitrogen R415, 1:400) in blocking buffer for one hour in the dark. After a final set of 3x 5 minute DPBS washes, coverslips were mounted onto slides with Prolong Glass mounting media (Invitrogen P36984) and allowed to cure overnight at RT in the dark. Cells from the CIC::NUTM1 experiment were imaged with a Yokogawa CSU-X1 spinning disk confocal on a Nikon Ti-E microscope equipped with a Photometrics cMYO cooled CCD camera using a 60x/1.45NA lens (Nikon) with NIS-Elements AR (v5.21.03, Nikon) software at consistent exposure times and laser powers. Cells from the ATXN1::DUX4 experiment were imaged with a ZEISS Axio Imager 2 with ZEISS ZEN 2 (blue edition, version 2.0.0.0) software and a 40x/1.4 Plan-Apochromat oil objective (ZEISS) at consistent exposure times and laser powers. Images were processed using FIJI/ImageJ (https://github.com/fiji/fiji). Briefly, multiple planes were converted to single images by maximum intensity projection, and brightness/contrast was consistently adjusted between all images.

For the CIC::NUTM1 cells, the primary antibody and dilution used was: anti-HA-tag clone C29F4 (Cell Signaling Technology 3724, dilution 1:300). The accompanying secondary antibody and dilution used was: anti-Rabbit IgG Alexa 647 (Invitrogen A-21245, dilution 1:300). For the ATXN1::DUX4 cells, the primary antibodies used were: anti-CIC (Invitrogen PA5-83721, dilution 1:200) and anti-HA-tag clone 6E2 with conjugated Alexa 488 (Cell Signaling Technology 2350, dilution 1:1000). The accompanying secondary antibody and dilution used was: anti-Rabbit IgG Alexa 647 (Invitrogen A-21245, dilution 1:500).

### RNA Sequencing and Analysis

The RNA sequencing data in this study was performed as three separate experiments, which are designated by the month and year they were executed. These designations delineate which supplemental files correspond to their respective experiments. The experiments and the plasmid transfections they included were:

1. April 2023 – EV, HA-CIC::DUX4, and HA-CIC(ex20)::NUTM1(ex6). These data are shown in Figures 2 and 3.
2. May 2024 – EV, HA-CIC::DUX4, HA-CIC(ex20)::NUTM1(ex6), HA-CIC(ex20)::NUTM1(ex6) dTAD, HA-CIC(ex20)::NUTM1(ex6) E, HA-CIC(ex18)::NUTM1(ex3), HA-CIC::LEUTX. These data are shown in Figures 5 and 6, and Supplemental Figure 4.
3. February 2025 – EV, HA-ATXN1::DUX4, HA-ATXN1::DUX4 delAXH. These data are shown in Figure 7 and Supplemental Figure 6.

All experiments were performed in 293T cells, with plasmids separately reverse transfected as described above into either 300,000 (April 2023, May 2024) or 500,000 (February 2025) 293T cells per transfection. For each experiment, three replicate sets of transfections were performed on different days. After approximately 48 hours, RNA was harvested from transfected cells with the RNeasy Mini kit (Qiagen) including an on-column DNase digest (Qiagen). The April 2023 experiment also included an extra, short Buffer RPE wash. The May 2024 and February 2025 experiments were then additionally purified with the Monarch Spin RNA Cleanup Kit (NEB). RNA from all experiments was checked for purity with a TapeStation (Agilent) by either ourselves, Novogene, or both, with RIN values of 9.5 or higher. RNA was then submitted to Novogene for library preparation and sequencing as described previously^18^. Samples were polyA enriched and libraries were generated using the NEBNext Ultra II Directional RNA Library Prep Kit for Illumina (NEB). Libraries were PE150 sequenced on a NovaSeq 6000 (April 2023) or NovaSeq X Plus (May 2024, February 2025). At least 6G of data (20 million paired-end 150bp reads) were generated for all samples.

The April 2023 experiment initially included an HA-CIC(ex18)::NUTM1(ex3) condition, which was excluded due to poor QC metrics in one of the replicates and not deposited in the NCBI Gene Expression Omnibus (GEO, RRID: SCR_005012)^72^. FASTQ files for all remaining data showed no concerning quality indications per both the Novogene report and our own FastQC analysis (RRID: SCR_014583)^73^. Full code describing the processing and analysis of the data from each experiment is available at https://github.com/cuylerluck/CICfamily_models. Processing and analysis was similar to previous work^18^. Briefly, for a given experiment, FASTQ files were aligned to the human GRCh38 genome with STAR (RRID: SCR_004463)^74^, including quantitation of gene counts using the option quantMode GeneCounts. Uniquely mapped read rates were between 87-94% for samples across all experiments. Custom R scripts (version 4.2.2)^75–84^, also available at the same GitHub repository as above, were used to separately perform differential gene expression analysis for each experiment. Briefly, for a given experiment, column 4 from STAR GeneCount output was merged between all samples. Then, edgeR (version 3.40.2, RRID: SCR_012802)^85–88^ was used for differential expression analysis with a GLM method and quasi-likelihood F-tests, with the resulting P-values then FDR corrected. Log_2_(counts per million) values were extracted from the trimmed mean of M-values (TMM)-normalized datasets using the cpm function. Where applicable, gene list functional profiling was performed using the gProfiler2^62^ package with multiple testing correction using the built-in gSCS method. The third replicate of the HA-ATXN1::DUX4 sample from the February 2025 experiment was found to be an outlier on an MDS plot, so we excluded the whole replicate and analyzed those data as duplicates using replicates 1 and 2. Consequently, the third replicate data was not deposited in GEO. For all analyzed data, differential expression results and lists of the E-ignorer and E-responder genes are available in Supplementary Datasets S1-S2 and S5-S13. The raw FASTQ files, log2(cpm) data, and counts data have been deposited in NCBI GEO^72^ and are accessible through GEO Series accession numbers GSE295624, GSE295623, and GSE295625 for the April 2023, May 2024, and February 2025 datasets, respectively.

