## Supplementary Figures and Supplementary Dataset Descriptions for "First Generation Tools for the Modeling of Capicua (CIC) - Family Fusion Oncoprotein-Driven Cancers"

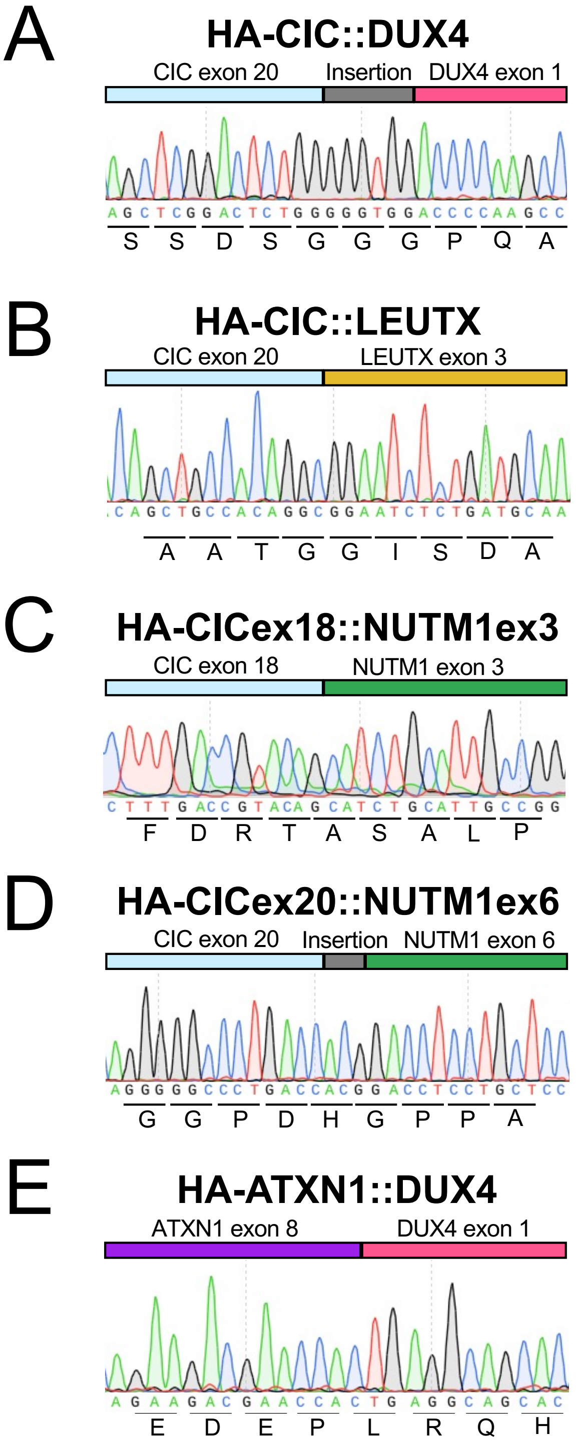


Supplementary Figure 1: Sanger sequencing of fusion breakpoints.
(A) – (E) Trans-breakpoint Sanger sequencing of fusion oncogene coding sequences in the plasmids described in Figure 1, with nucleotides and amino acid sequences (bottom) indicated. Exon numbering is based off of the reference sequences described in the Methods section.

A

| Construct | 5' Partner Source | 3' Partner Source | Cloning Approach |
|---|---|---|---|
| **HA-CIC::LEUTX** | Origene # RC215209 | Synthesized coding sequence of NM_001382345.1 (Twist Bioscience) | NEB HiFi |
| **HA-CICex18:: NUTM1ex3** | HA-CIC::DUX4 plasmid (PMID: 16717057) | Addgene #171630 | NEB HiFi |
| **HA-CICex20:: NUTM1ex6** | HA-CIC::DUX4 plasmid (PMID: 16717057) | Addgene #171630 | NEB HiFi |
| **HA-ATXN1::DUX4** | Addgene #48189 | Addgene #99282 | NEB HiFi |

*All: Backbone derived from Sino Biologicals pCMV3-GFP-FLAG, catalog # AG13105-CF. 3xHA tag/linker motif derived from HA-CIC::DUX4 plasmid.*

Supplementary Figure 2: Cloning approaches for synthetic *CIC*-family fusions.
(A) Description of the sequences used to derive the 5' and 3' fusion partners as well as the cloning approach used to generate each of the synthetic *CIC*-family fusion coding sequences. A more in-depth description of cloning approaches and sequence minutiae is available in the Methods section.

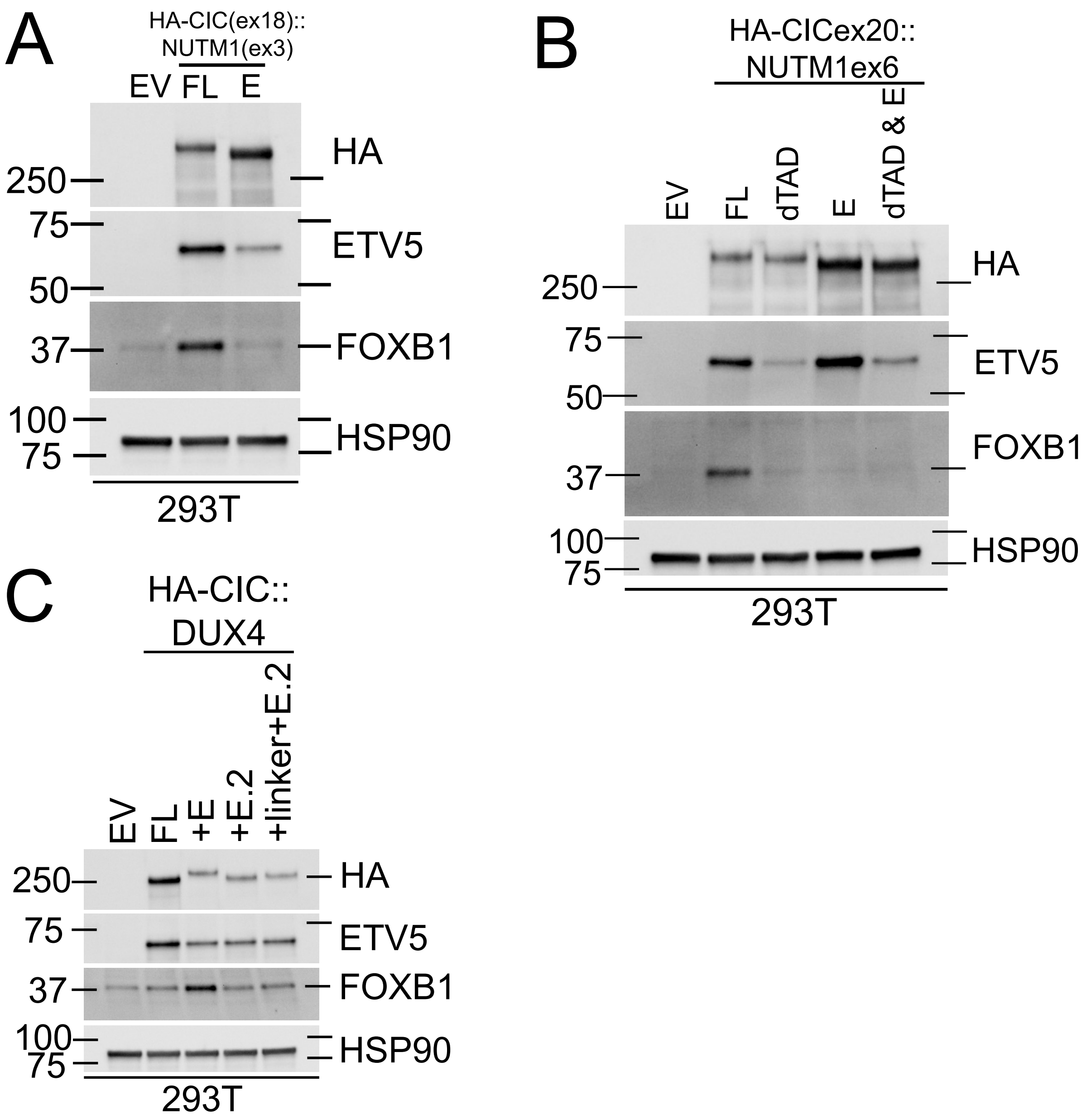


Supplementary Figure 3: Structure-function studies of the NUTM1 E region in CIC::NUTM1 and CIC::DUX4. (A) Immunoblot of 293T cells approximately 48 hours after transfection with empty vector (EV) or the indicated versions of HA-CIC(ex18)::NUTM1(ex3): full length (FL) or with the E region deleted. Representative of three independent experiments.
(B) Immunoblot of 293T cells approximately 48 hours after transfection with empty vector (EV) or the indicated versions of HA-CIC(ex20)::NUTM1(ex6): full length (FL), p300-interacting domain deleted (dTAD), E region deleted (E), or both p300-interacting and E domains deleted (dTAD & E). Experiment performed once.
(C) Immunoblot of 293T cells approximately 48 hours after transfection with empty vector (EV) or the indicated versions of HA-CIC::DUX4: full length (FL), or with the E region or E.2 region or a $(GSSG)_3$ linker plus the E.2 region from panels B-D cloned onto the C-terminus of the HA-CIC::DUX4 coding sequence. Experiment performed once.

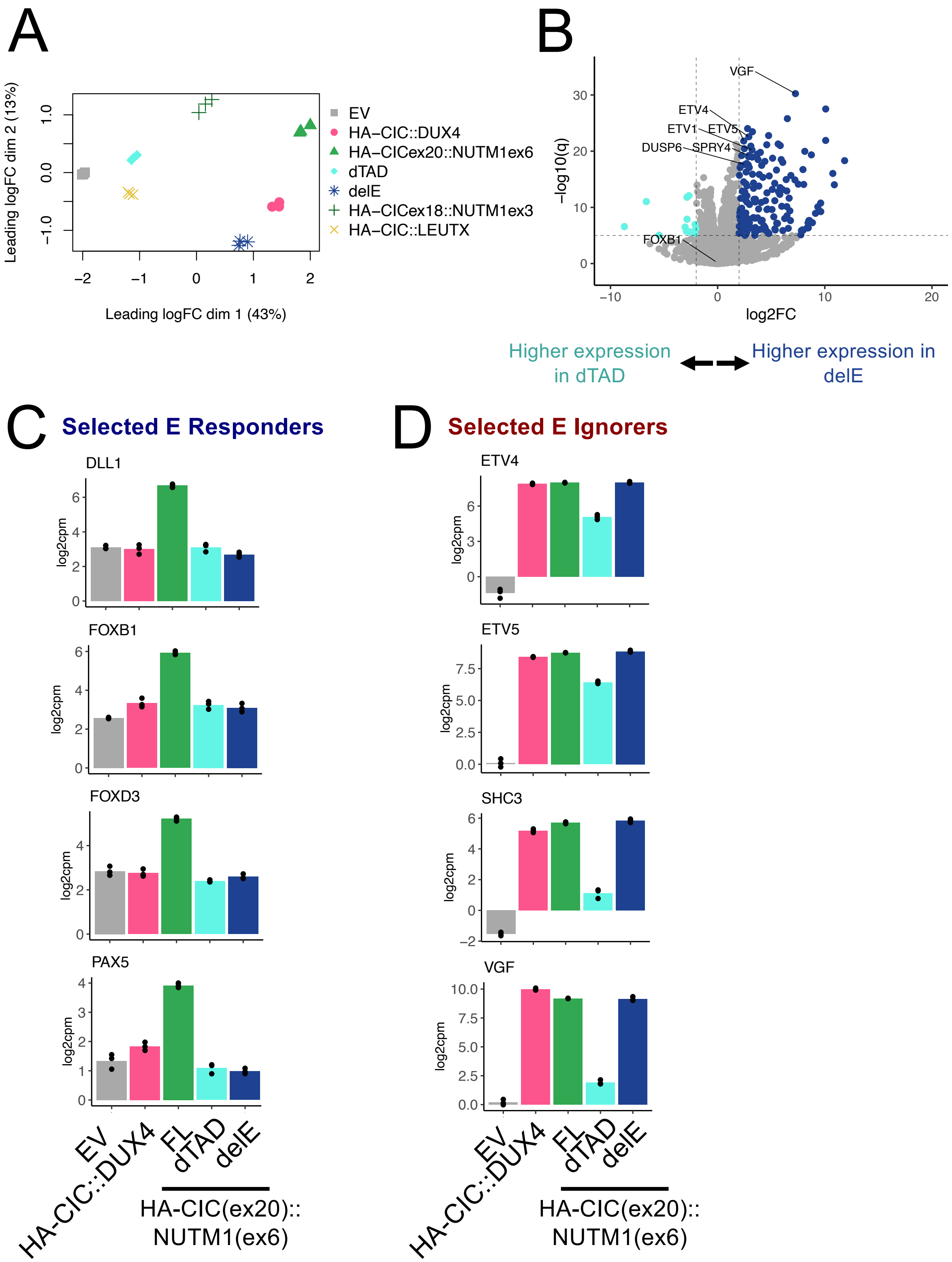


Supplementary Figure 4: Analysis of the NUTM1 E domain regulated transcriptional program. (A) edgeR multi-dimensional scaling plot of RNA-seq data from 293T cells 48 hours after transfection with empty vector (EV) or the indicated fusion-expressing constructs. dTAD and delE refer to deletion of the p300-interacting region or deletion of the E region in HA-CIC(ex20)::NUTM1(ex6), respectively.

(B) Volcano plot of RNA-seq data comparing gene expression in 293T transfected with the E-deleted mutant (delE) of HA-CIC(ex20)::NUTM1(ex6) vs the p300-interacting region deletion mutant (dTAD) of the same fusion. Several known CIC/CIC::DUX4/CIC::NUTM1 target genes are labelled.

(C) – (D) RNA-seq $log_2$(counts per million) values for selected genes from the E responder (C) and E ignorer (D) gene sets. FL, dTAD, and delE refer to the full length, p300-interacting region deletion mutant, and E region deletion mutant versions of HA-CIC(ex20)::NUTM1(ex6), respectively.

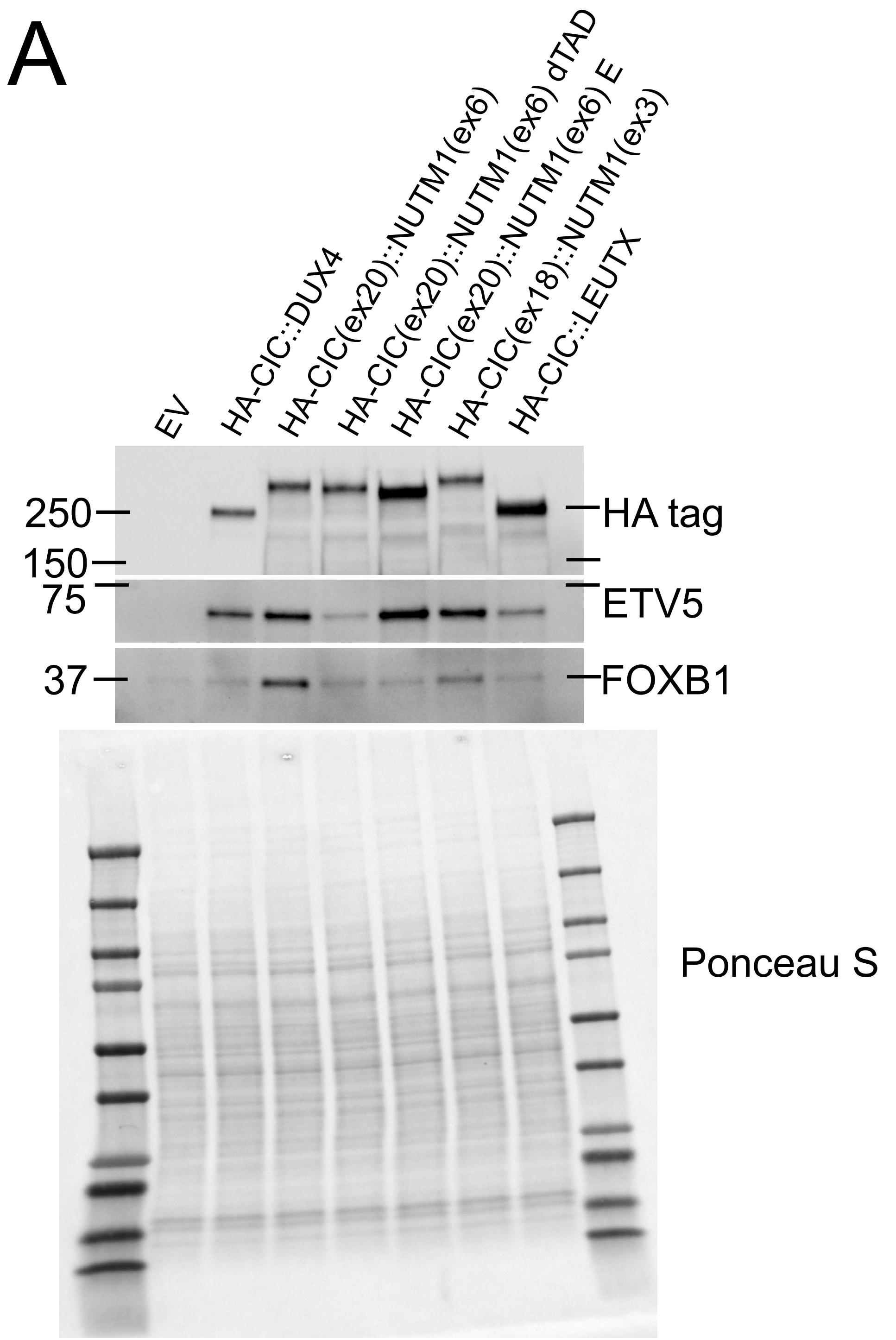


Supplementary Figure 5: Validation of expression for selected *CIC*-family fusions and mutants used for RNA-seq experiments.

(A) Immunoblot of 293T cells approximately 48 hours post transfection with empty vector (EV) or the indicated constructs. dTAD refers to deletion of the NUTM1 p300-interacting region and E refers to deletion of the NUTM1 E domain. Ponceau S staining of total protein is shown below. Representative of three independent experiments.

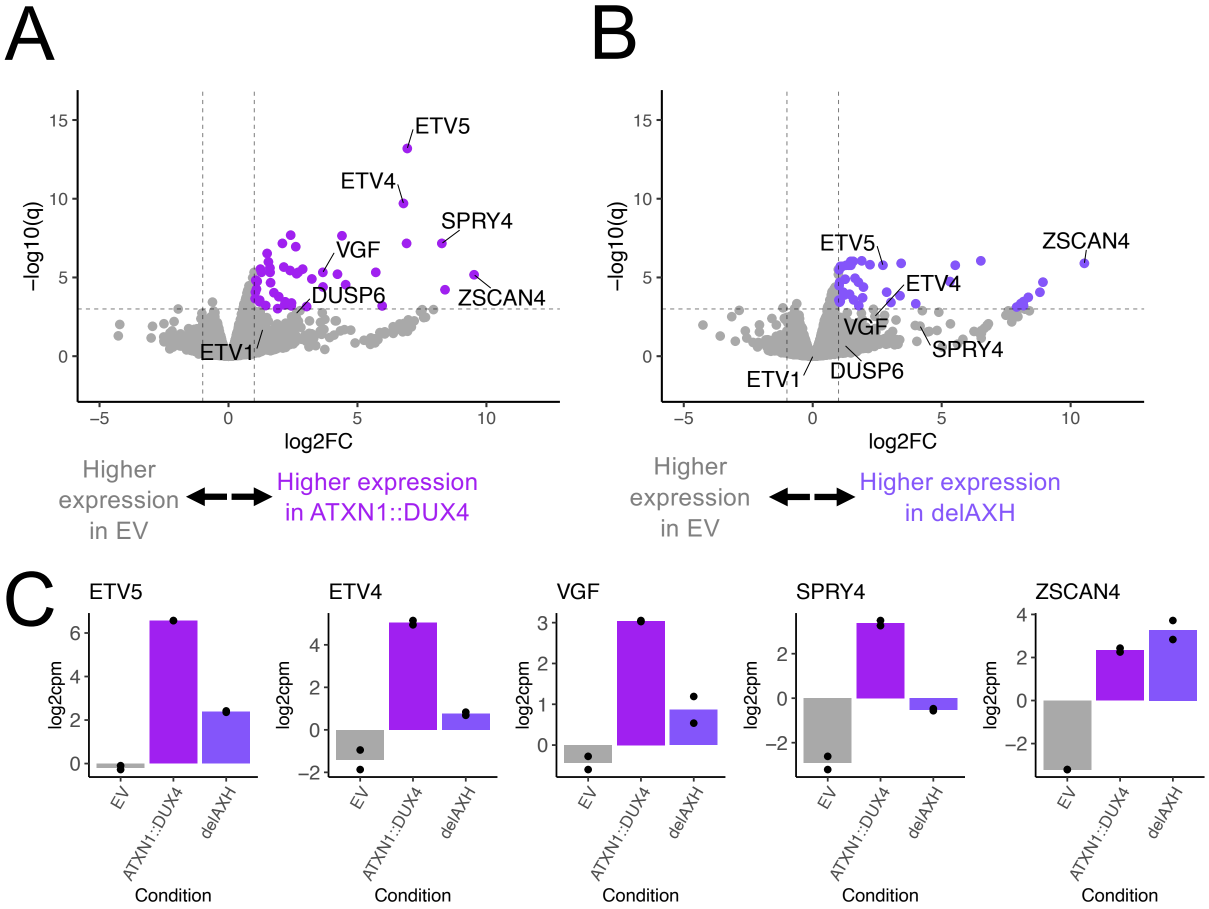


Supplementary Figure 6: Analysis of differential gene induction by ATXN1::DUX4 possessing or lacking the AXH domain.

(A) – (B) Volcano plots of RNA-seq expression data from 293T cells approximately 48 hours after transfection with the indicated constructs. (A) shows the full length HA-ATXN1::DUX4 vs. EV comparison, while (B) shows the AXH-deleted HA-ATXN1::DUX4 vs. EV comparison. Select CIC and DUX4 target genes are indicated.

(C) $\log_2$(counts per million) RNA-seq expression of selected CIC and DUX4 target genes in 293T cells approximately 48 hours after transfection with the indicated constructs.

**Supplementary Dataset Descriptions**

Supplementary Dataset 1: .txt file, list of E ignorer genes.

Supplementary Dataset 2: .txt file, list of E responder genes.

Supplementary Dataset 3: .pdf file, uncropped blots and Ponceau S staining for all western blots.

Supplementary Dataset 4: .xlsx file, list of primer sequences used for cloning.

Supplementary Dataset 5: tab-delimited .txt file, differential expression results of CIC::DUX4 vs EV comparison from April 2023 data, larger $log_2$ fold change value indicates higher expression in CIC::DUX4 condition.

Supplementary Dataset 6: tab-delimited .txt file, differential expression results of CIC(ex20)::NUTM1(ex6) vs EV comparison from April 2023 data, larger $log_2$ fold change value indicates higher expression in CIC(ex20)::NUTM1(ex6) condition.

Supplementary Dataset 7: tab-delimited .txt file, differential expression results of CIC::DUX4 vs EV comparison from May 2024 data, larger $log_2$ fold change value indicates higher expression in CIC::DUX4 condition.

Supplementary Dataset 8: tab-delimited .txt file, differential expression results of CIC(ex20)::NUTM1(ex6) vs EV comparison from May 2024 data, larger $log_2$ fold change value indicates higher expression in CIC(ex20)::NUTM1(ex6) condition.

Supplementary Dataset 9: tab-delimited .txt file, differential expression results of CIC(ex18)::NUTM1(ex3) vs EV comparison from May 2024 data, larger $log_2$ fold change value indicates higher expression in CIC(ex18)::NUTM1(ex3) condition.

Supplementary Dataset 10: tab-delimited .txt file, differential expression results of CIC::LEUTX vs EV comparison from May 2024 data, larger $log_2$ fold change value indicates higher expression in CIC::LEUTX condition.

Supplementary Dataset 11: tab-delimited .txt file, differential expression results of ATXN1::DUX4 vs EV comparison from February 2025 data, larger $log_2$ fold change value indicates higher expression in ATXN1::DUX4 condition.

Supplementary Dataset 12: tab-delimited .txt file, differential expression results of AXH-deleted ATXN1::DUX4 vs EV comparison from February 2025 data, larger $log_2$ fold change value indicates higher expression in AXH-deleted ATXN1::DUX4 condition.

Supplementary Dataset 13: tab-delimited .txt file, differential expression results of AXH-deleted ATXN1::DUX4 vs full length ATXN1::DUX4 comparison from February 2025 data, larger $log_2$ fold change value indicates higher expression in AXH-deleted ATXN1::DUX4 condition.
