## Supplementary Dataset S3 for "First Generation Tools for the Modeling of Capicua (CIC) - Family Fusion Oncoprotein-Driven Cancers"

### Figure 2 Panel A

Left = Chemiluminescence, Right = composite of chemi + visual light

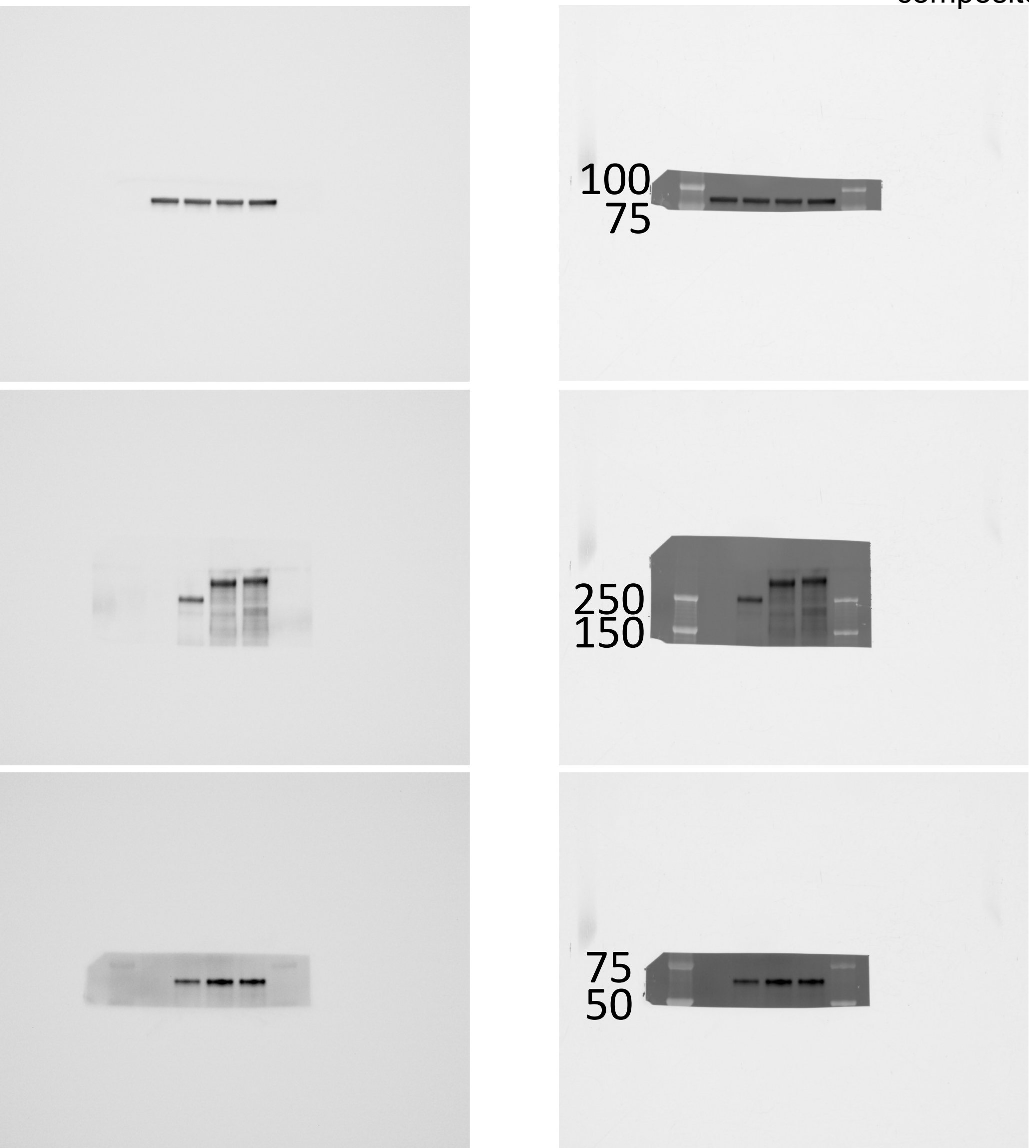


HSP90

HA tag

ETV5

Ponceau S

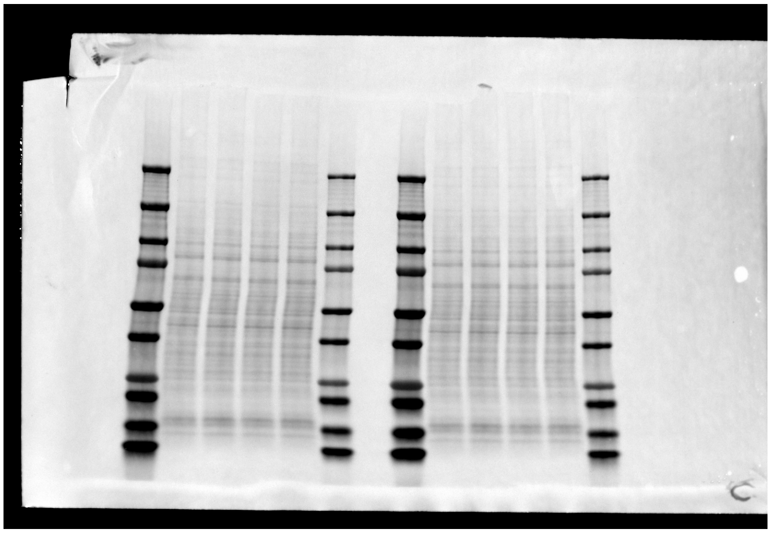

Sample loading order, left to right:
7 uL ladder
EV
HA-CIC::DUX4
HA-CIC(ex20)::NUTM1(ex6)
HA-CIC(ex18)::NUTM1(ex3)
3 uL ladder

Note: HA and ETV5 were cut from the right side samples, HSP90 from the left.

### Figure 2 Panel C

Left = Chemiluminescence, Right = composite of chemi + visual light

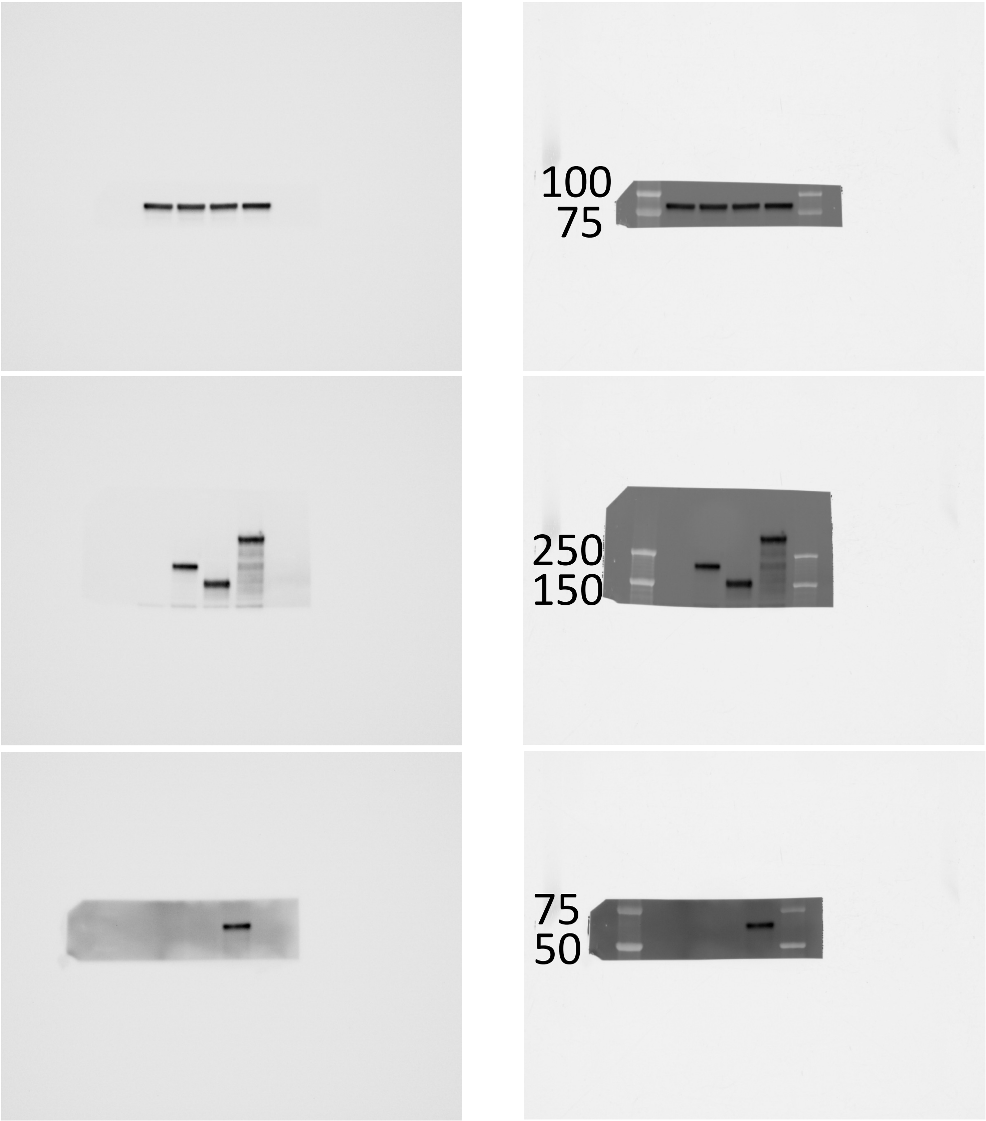


HSP90

HA tag

ETV5

Ponceau S

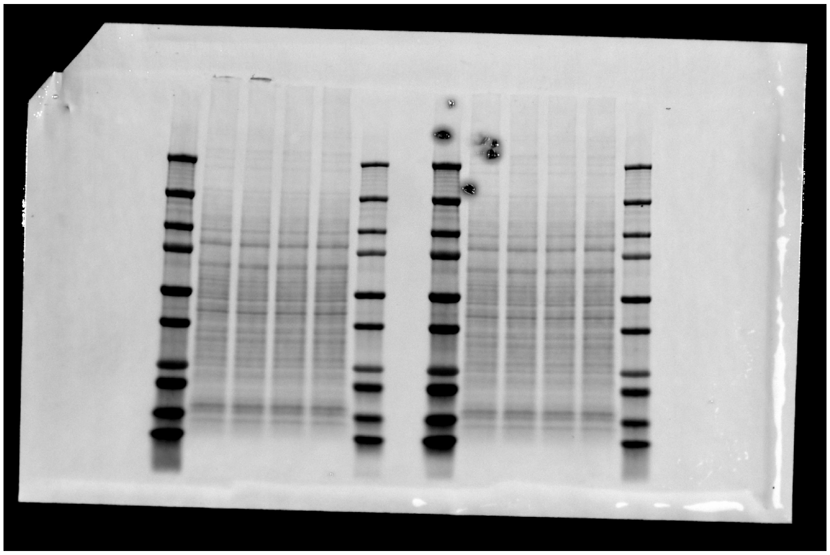

Sample loading order, left to right:
7 uL ladder
EV
HA-CIC (truncated)
HA-NUTM1 (truncated)
HA-CIC(ex18)::NUTM1(ex3)
3 uL ladder

Note: HA and ETV were cut from the left side samples, HSP90 from the right.

### Figure 2 Panel D

Left = Chemiluminescence, Right = composite of chemi + visual light

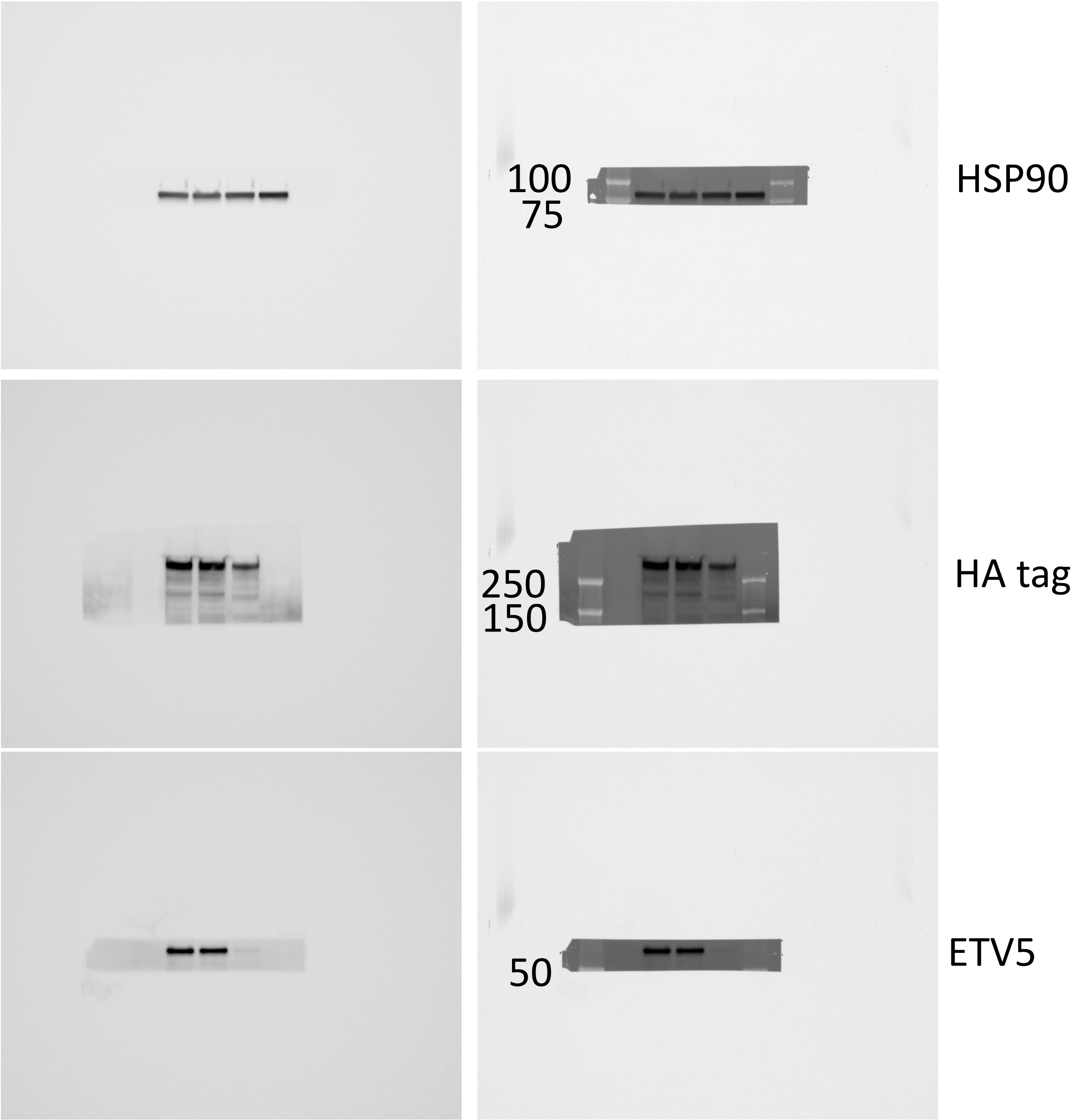


Ponceau S

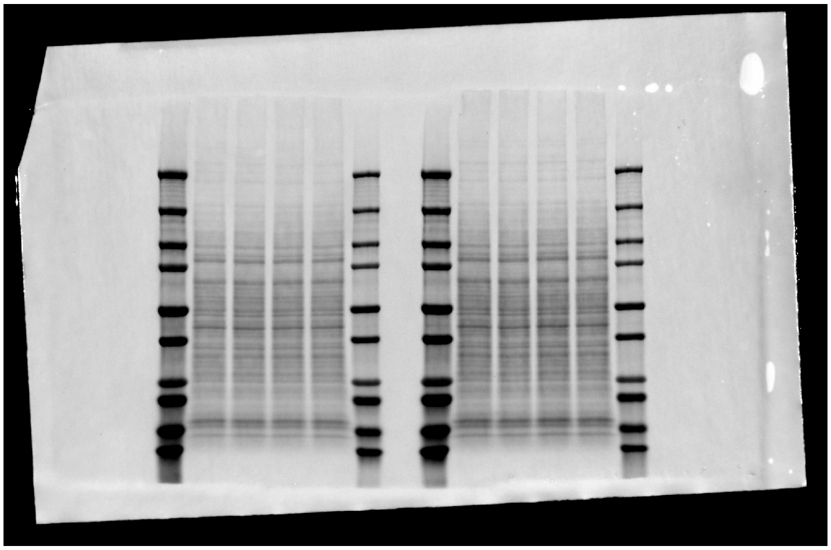

Sample loading order, left to right:
7 uL ladder
EV
HA-CIC(ex20)::NUTM1(ex6)
HA-CIC(ex20)::NUTM1(ex6) dC1
HA-CIC(ex20)::NUTM1(ex6) dC1/dHMG
3 uL ladder

Note: all blots taken from the right sided samples, the left is a different replicate.

### Figure 3 Panel B

Chemi + composite blots on following slides

Ponceau S

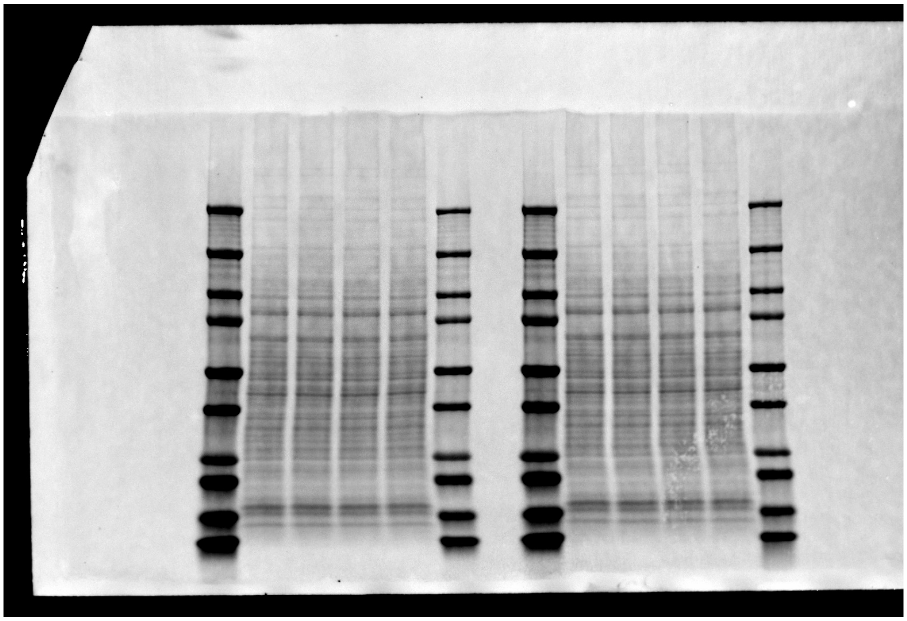

Note: SOX2 and FOXC2 were cut from the right side samples, HSP90 and FOXG2 from the left.

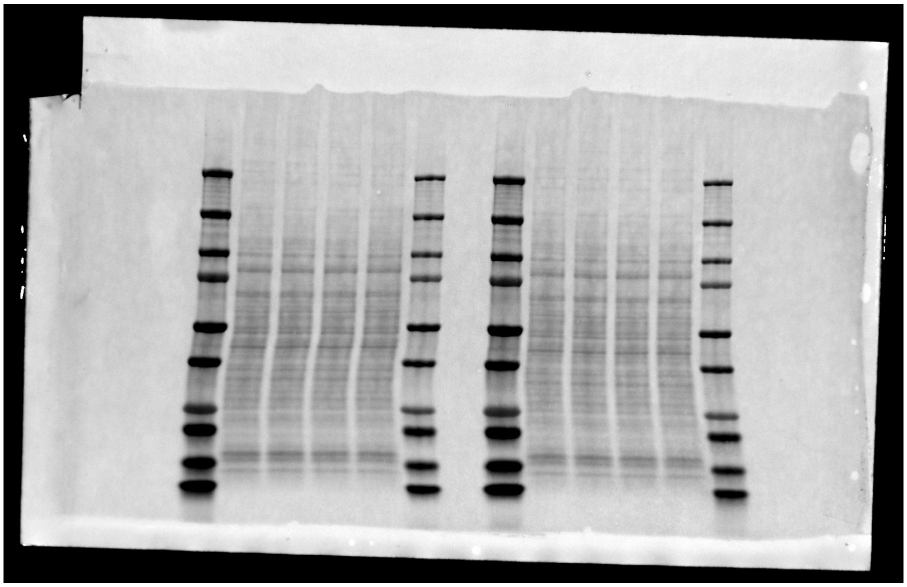

Note: HA tag, ETV5, and FOXB1 were cut from the left side samples, FOXD3 from the right.

Sample loading order, left to right:
7 uL ladder
EV
HA-CIC::DUX4
HA-CIC(ex20)::NUTM1(ex6)
HA-CIC(ex18)::NUTM1(ex3)
3 uL ladder

### Figure 3 Panel B (continued)

Left = Chemiluminescence, Right = composite of chemi + visual light

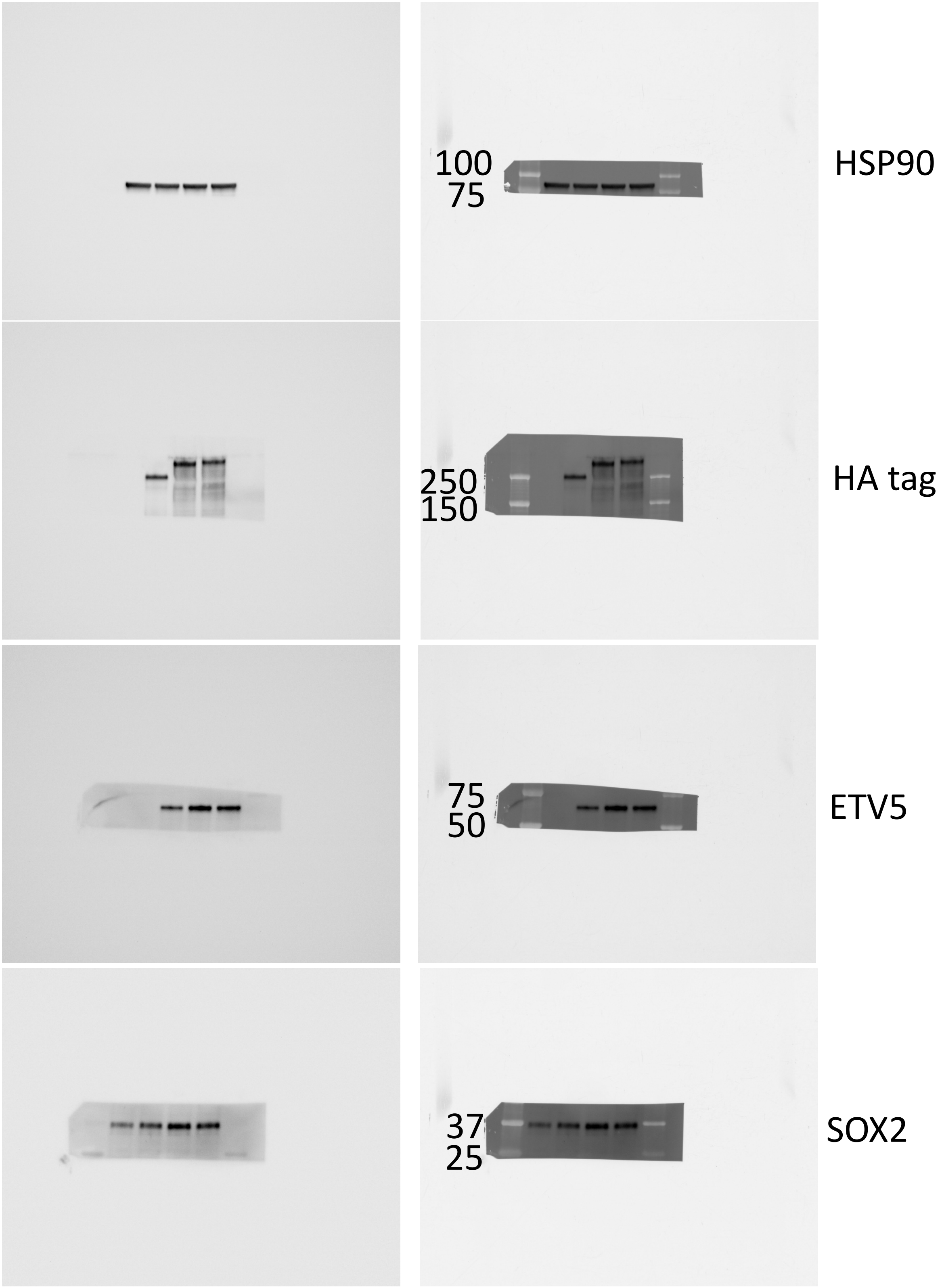


Sample loading order, left to right:
7 uL ladder
EV
HA-CIC::DUX4
HA-CIC(ex20)::NUTM1(ex6)
HA-CIC(ex18)::NUTM1(ex3)
3 uL ladder

### Figure 3 Panel B (continued)

Left = Chemiluminescence, Right = composite of chemi + visual light

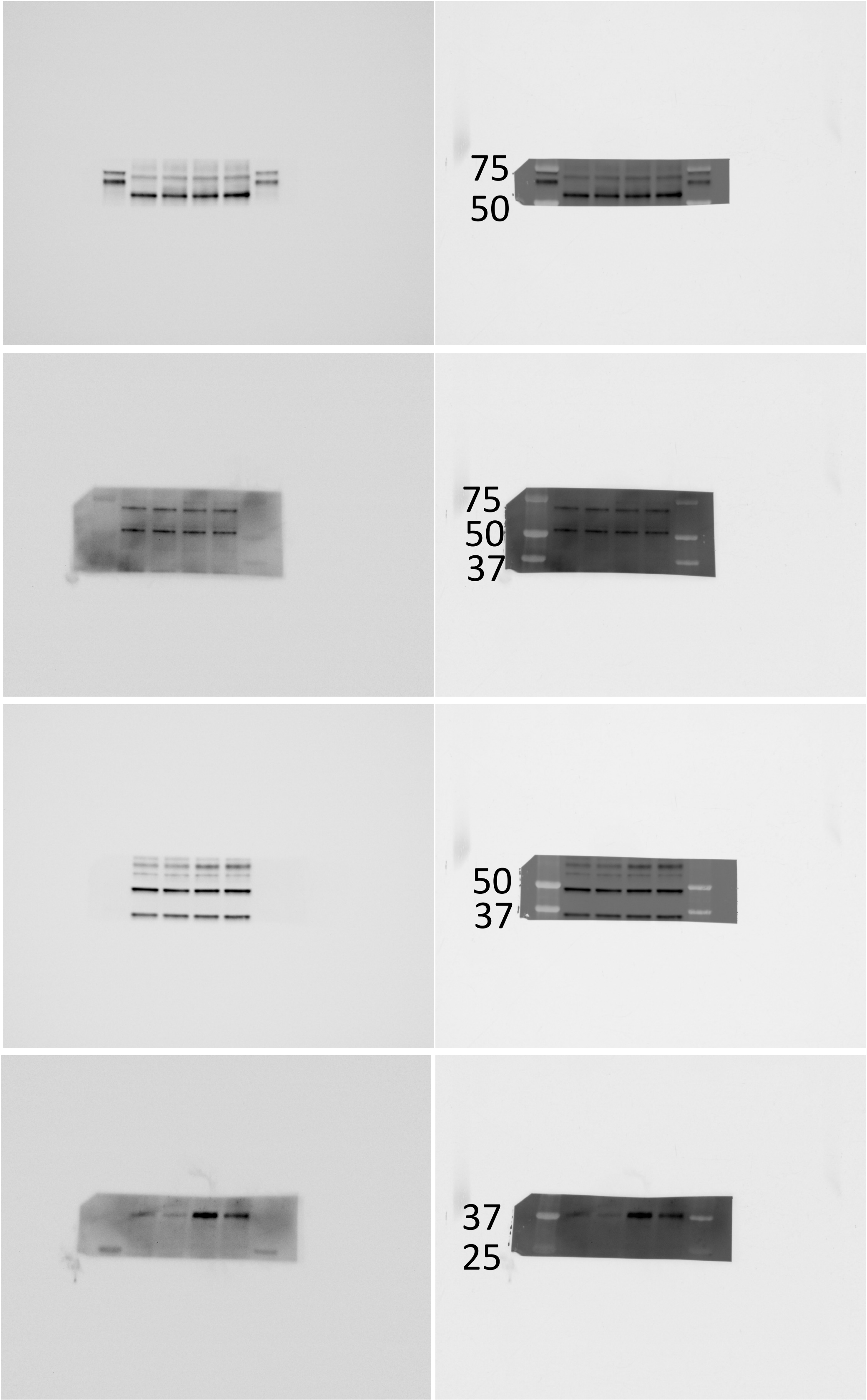


FOXC2

FOXD3

FOXG1

FOXB1

Sample loading order, left to right:
7 uL ladder
EV
HA-CIC::DUX4
HA-CIC(ex20)::NUTM1(ex6)
HA-CIC(ex18)::NUTM1(ex3)
3 uL ladder

### Figure 3 Panel C

Left = Chemiluminescence, Right = composite of chemi + visual light

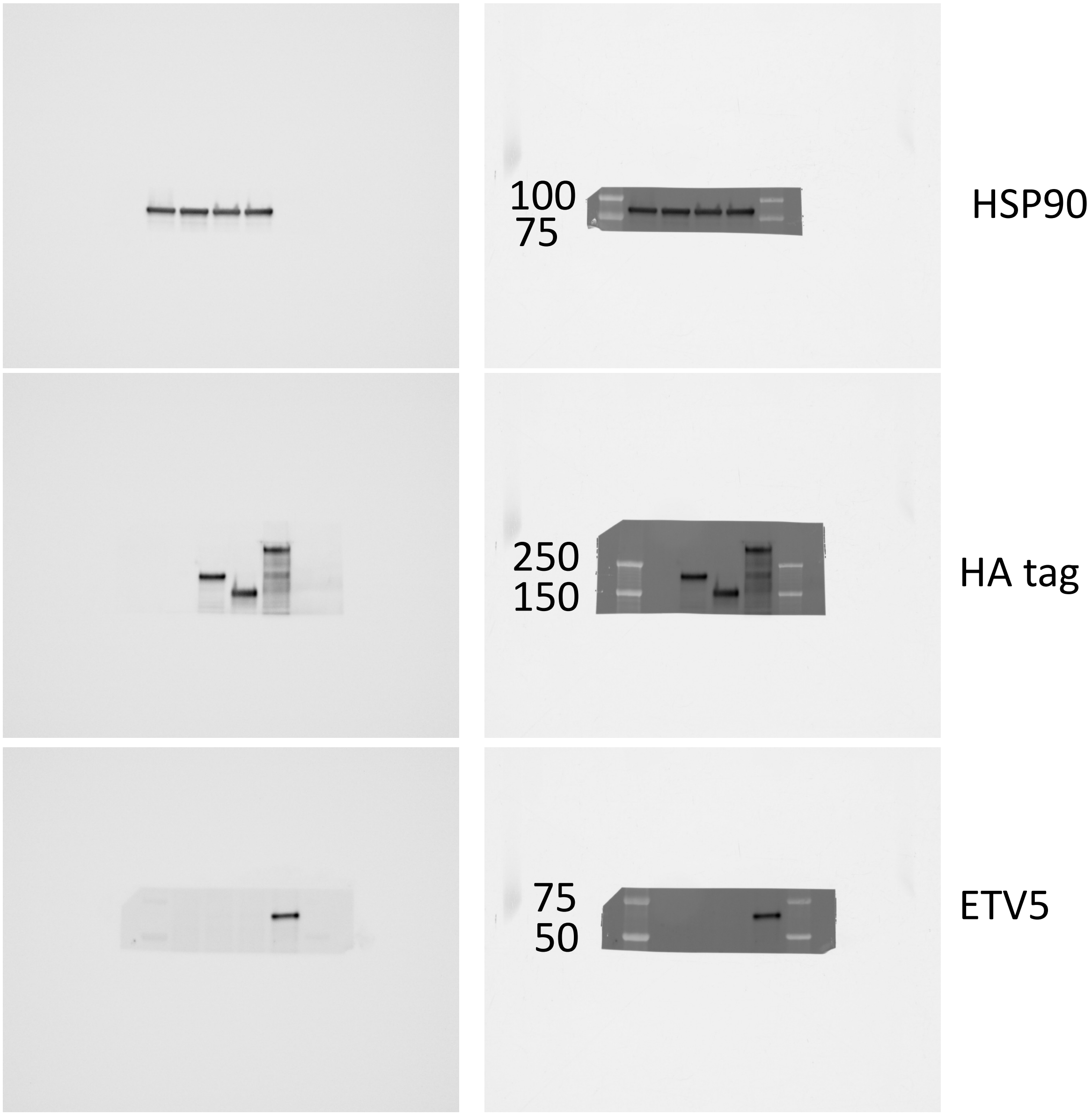


Ponceau S

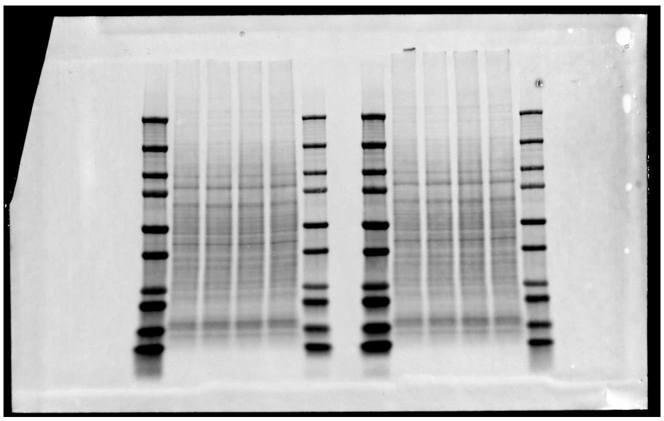

Sample loading order, left to right:
7 uL ladder
EV
HA-CIC (truncated)
HA-NUTM1 (truncated)
HA-CIC(ex18)::NUTM1(ex3)
3 uL ladder

Note: HA tag and ETV5 were cut from the left side samples, HSP90 and FOXB1 from the right.

### Figure 3 Panel C (continued)

Left = Chemiluminescence, Right = composite of chemi + visual light

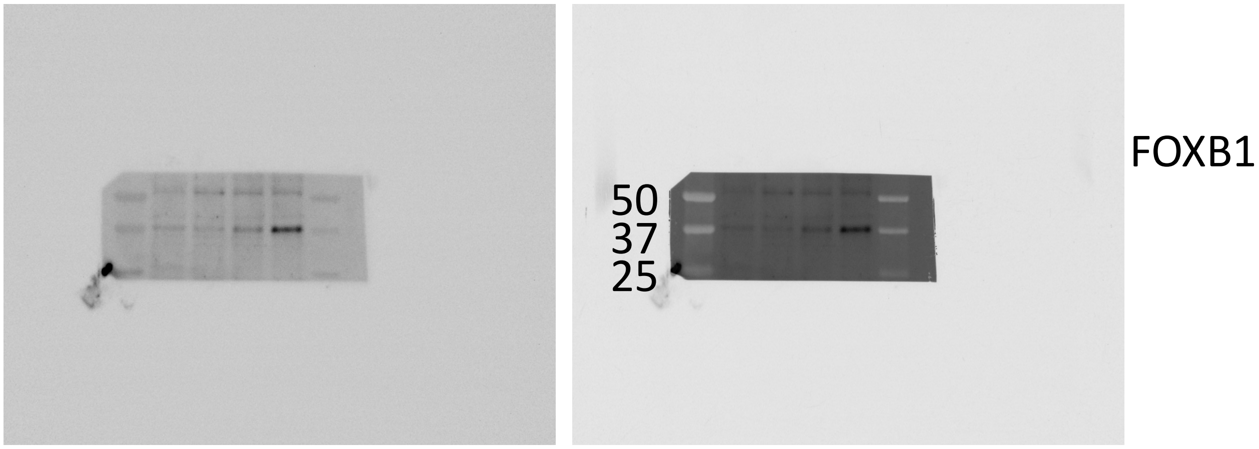


Sample loading order, left to right:
7 uL ladder
EV
HA-CIC (truncated)
HA-NUTM1 (truncated)
HA-CIC(ex18)::NUTM1(ex3)
3 uL ladder

### Figure 3 Panel D

Left = Chemiluminescence, Right = composite of chemi + visual light

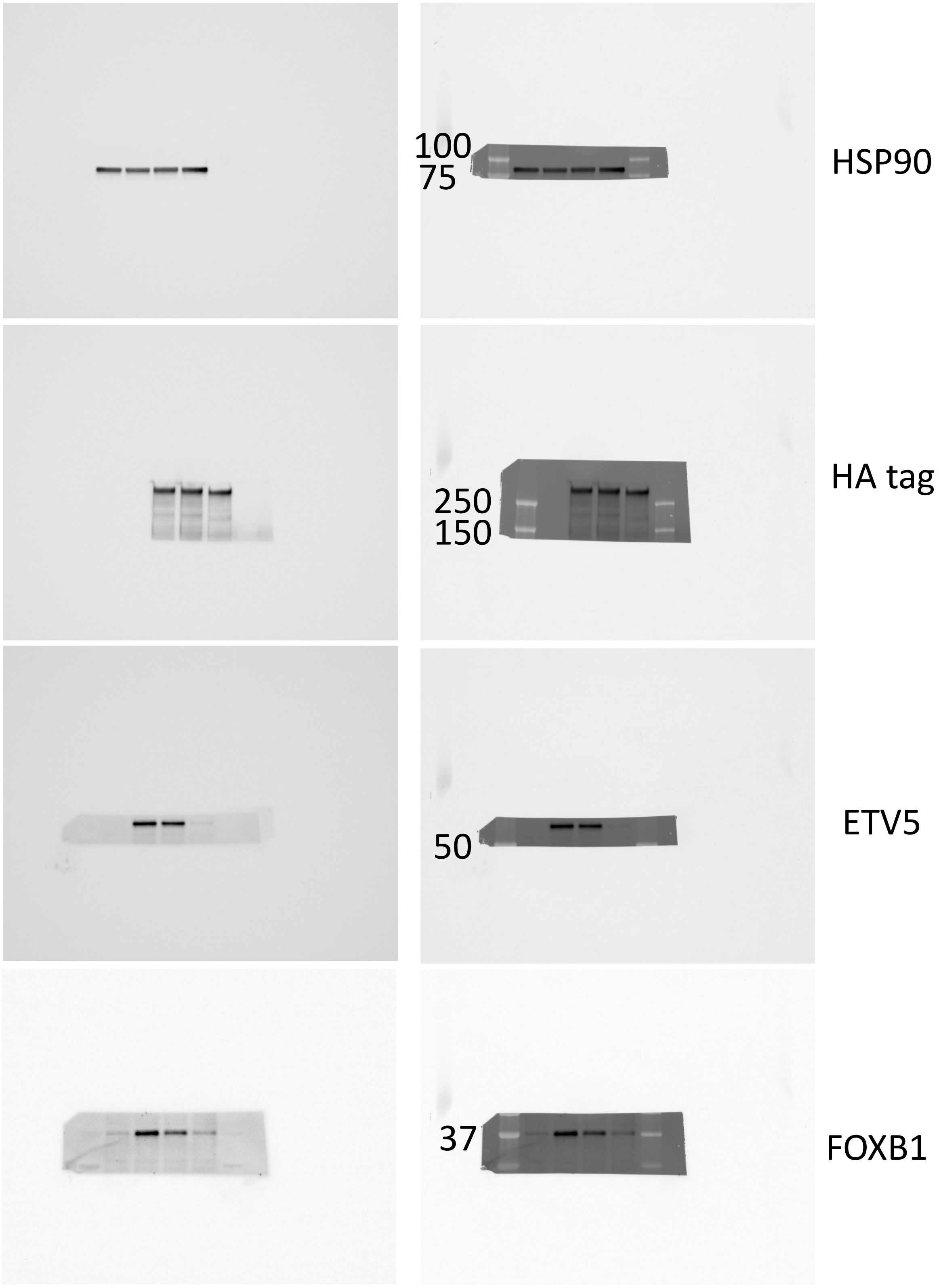


Ponceau S

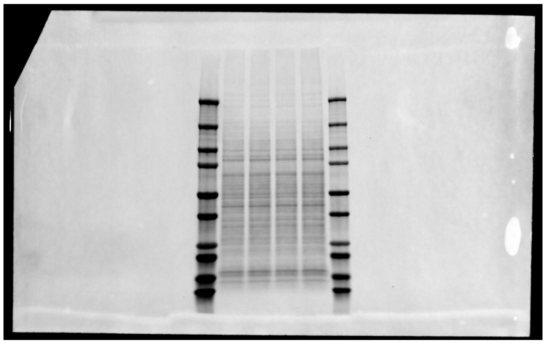

Sample loading order, left to right:
7 uL ladder
EV
HA-CIC(ex20)::NUTM1(ex6)
HA-CIC(ex20)::NUTM1(ex6) dC1
HA-CIC(ex20)::NUTM1(ex6) dC1/dHMG
3 uL ladder

### Figure 4 Panel A

Left = Chemiluminescence, Right = composite of chemi + visual light

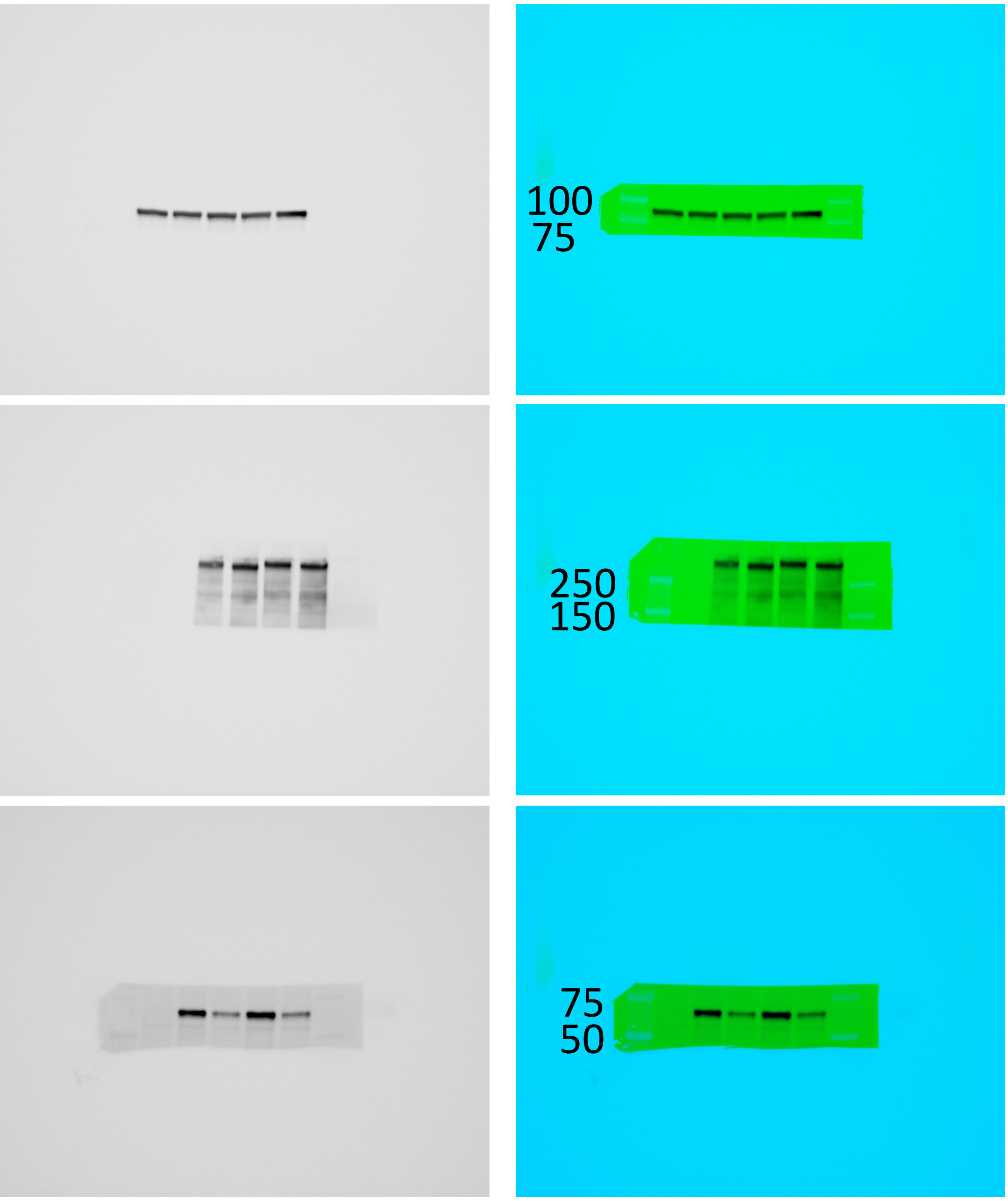


HSP90

HA tag

ETV5

Ponceau S

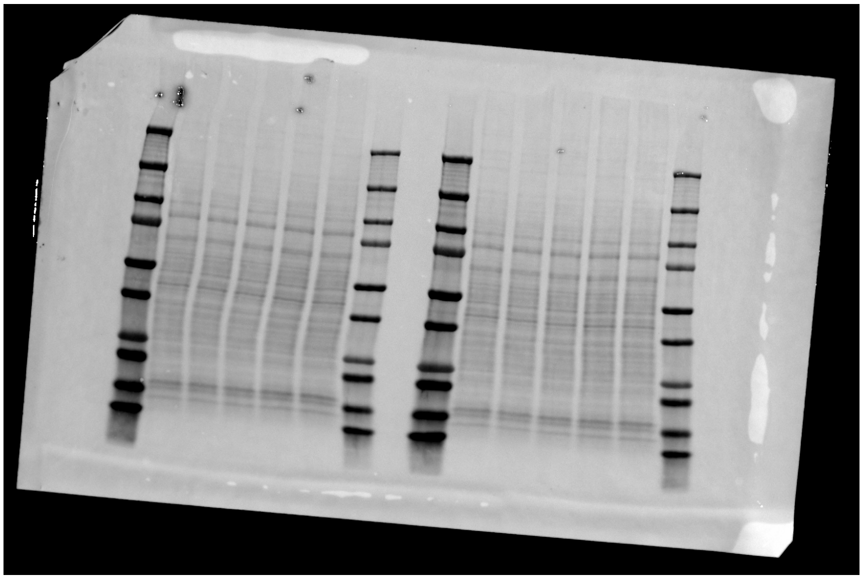

Sample loading order, left to right:

7 uL ladder

EV

HA-CIC(ex20)::NUTM1(ex6)

HA-CIC(ex20)::NUTM1(ex6) dTAD

HA-CIC(ex18)::NUTM1(ex3)

HA-CIC(ex18)::NUTM1(ex3) dTAD

3 uL ladder

Note: HA and ETV5 were cut from the left side samples, HSP90 from the right.

### Figure 4 Panel C

Chemi + composite blots on following slides

Ponceau S

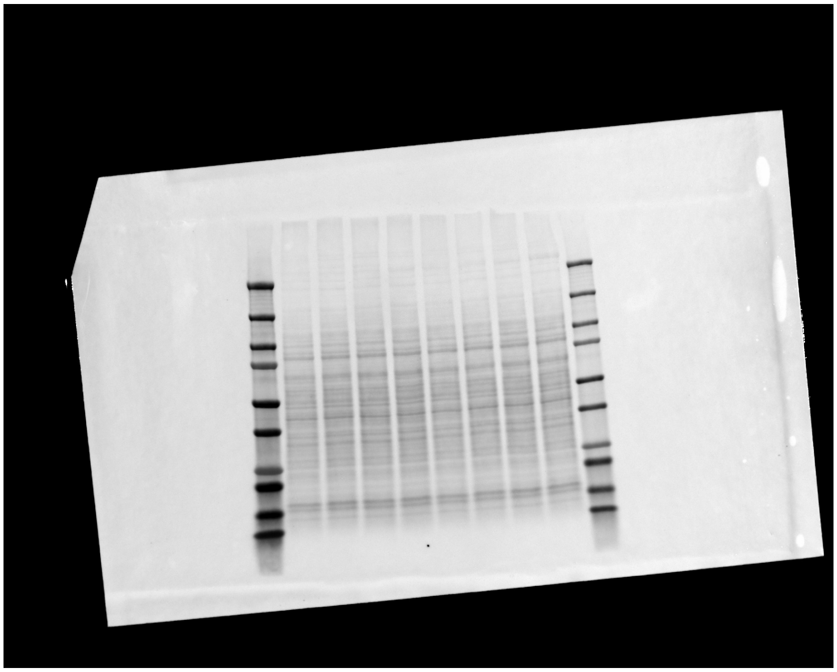

Note: HSP90, HA tag, and FOXB1 were cut from this blot.

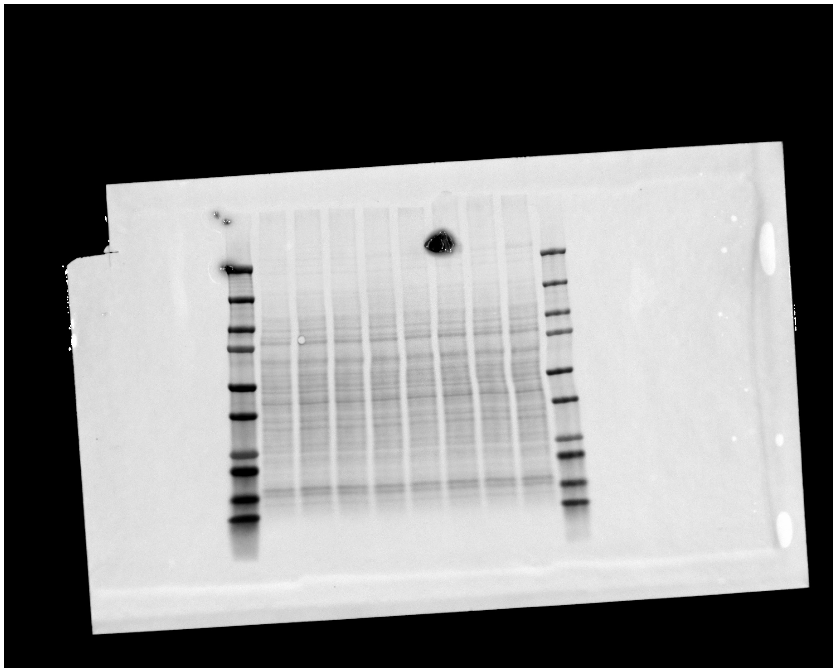

Note: ETV5 was cut from this blot.

Sample loading order, left to right:
7 uL ladder
EV
HA-CIC(ex20)::NUTM1(ex6)
HA-CIC(ex20)::NUTM1(ex6) dTAD
HA-CIC(ex20)::NUTM1(ex6) A
HA-CIC(ex20)::NUTM1(ex6) B
HA-CIC(ex20)::NUTM1(ex6) C
HA-CIC(ex20)::NUTM1(ex6) D
HA-CIC(ex20)::NUTM1(ex6) E
3 uL ladder

### Figure 4 Panel C (continued)

Left = Chemiluminescence, Right = composite of chemi + visual light

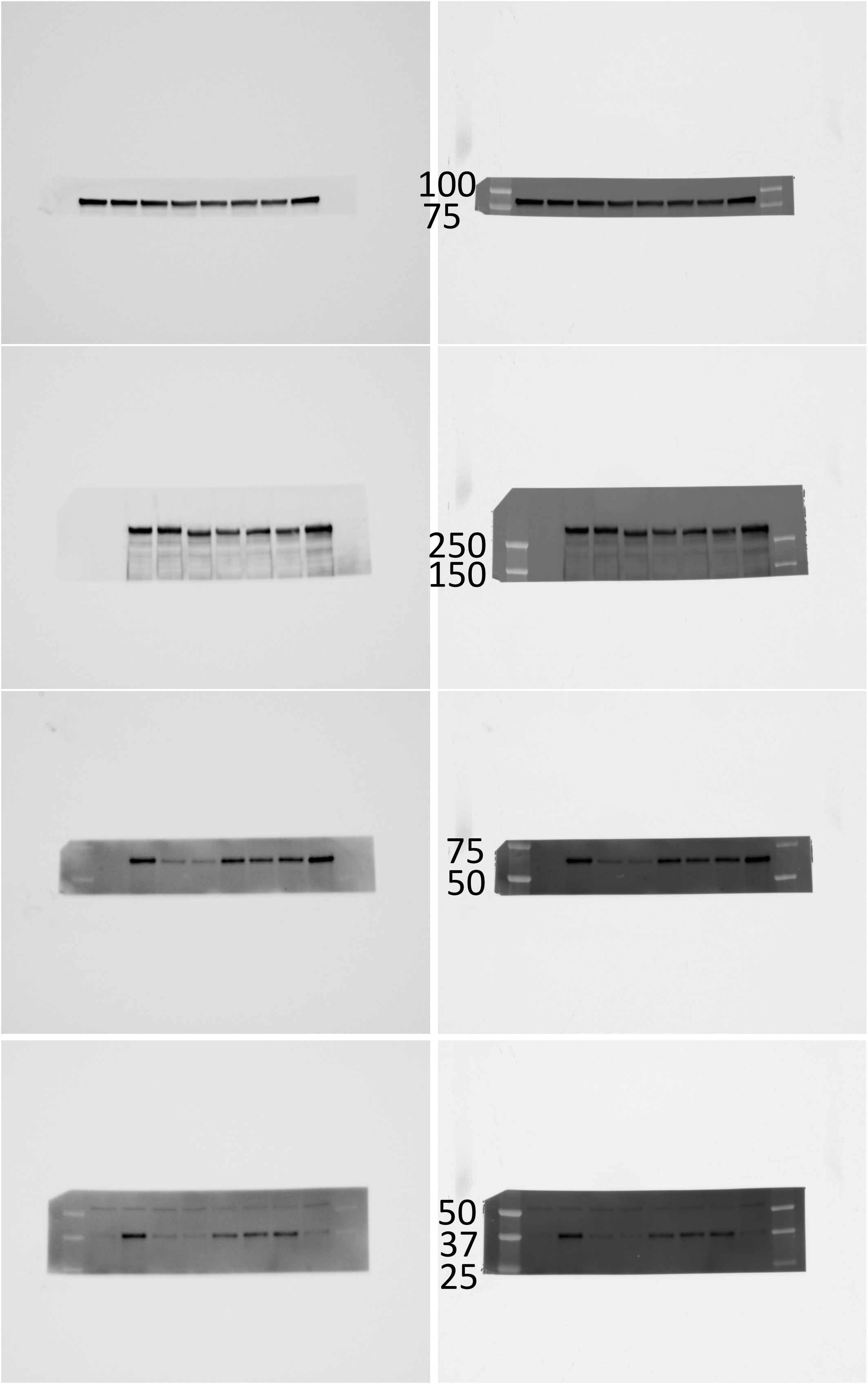


HSP90

HA tag

ETV5

FOXB1

Sample loading order, left to right:
7 uL ladder
EV
HA-CIC(ex20)::NUTM1(ex6)
HA-CIC(ex20)::NUTM1(ex6) dTAD
HA-CIC(ex20)::NUTM1(ex6) A
HA-CIC(ex20)::NUTM1(ex6) B
HA-CIC(ex20)::NUTM1(ex6) C
HA-CIC(ex20)::NUTM1(ex6) D
HA-CIC(ex20)::NUTM1(ex6) E
3 uL ladder

### Figure 4 Panel D

Left = Chemiluminescence, Right = composite of chemi + visual light

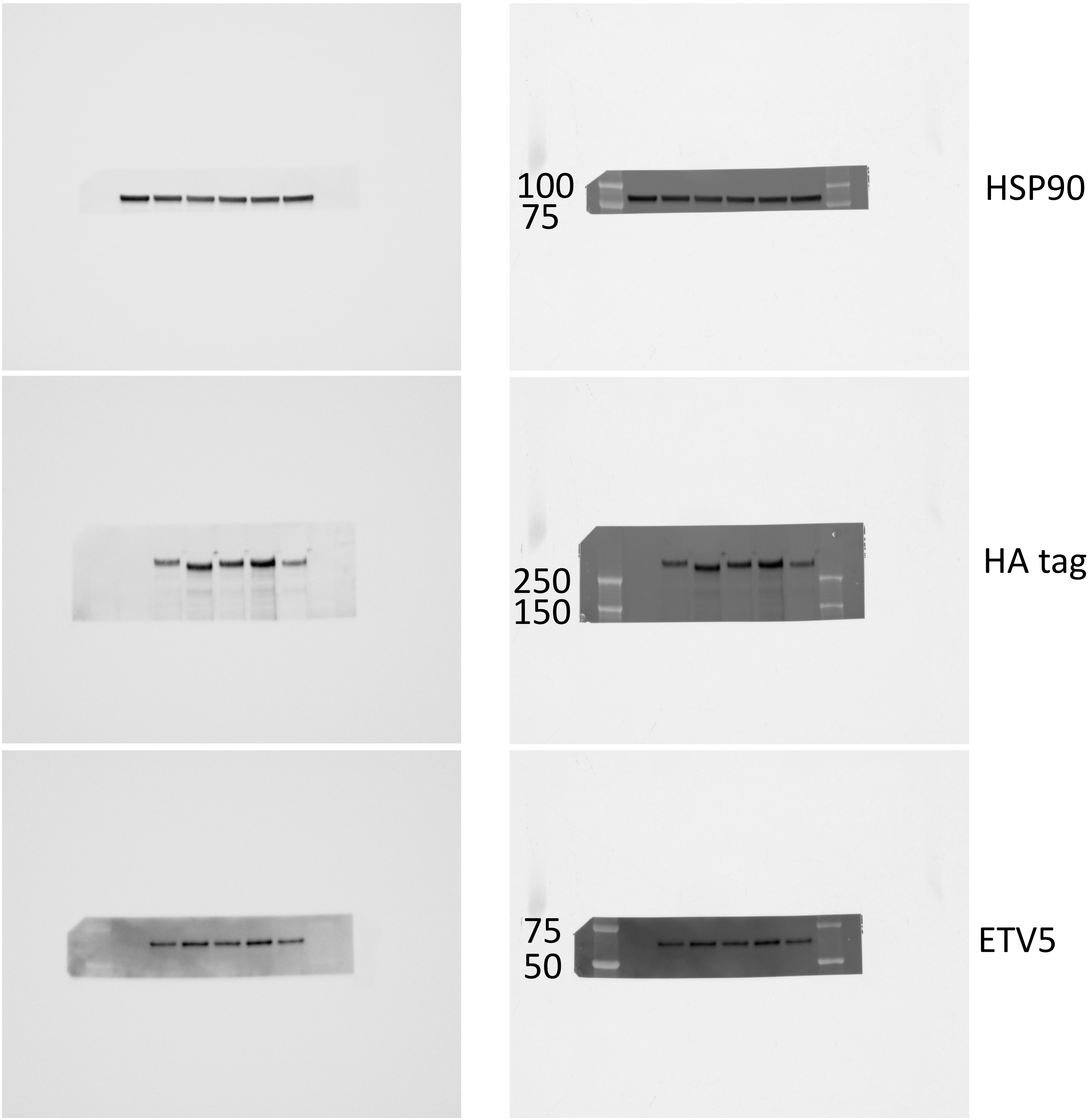


Ponceau S

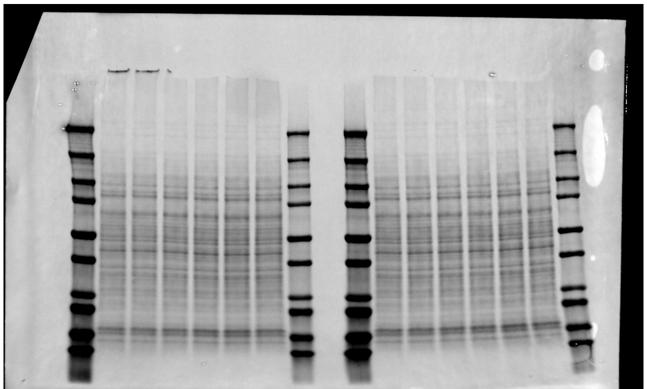

Sample loading order, left to right:
7 uL ladder
EV
HA-CIC(ex20)::NUTM1(ex6)
HA-CIC(ex20)::NUTM1(ex6) E
HA-CIC(ex20)::NUTM1(ex6) E.1
HA-CIC(ex20)::NUTM1(ex6) E.2
HA-CIC(ex20)::NUTM1(ex6) E.3
3 uL ladder

Note: HSP90 and FOXB1 were cut from the left side samples, HA tag and ETV5 from the right.

### Figure 4 Panel D (continued)

Left = Chemiluminescence, Right = composite of chemi + visual light

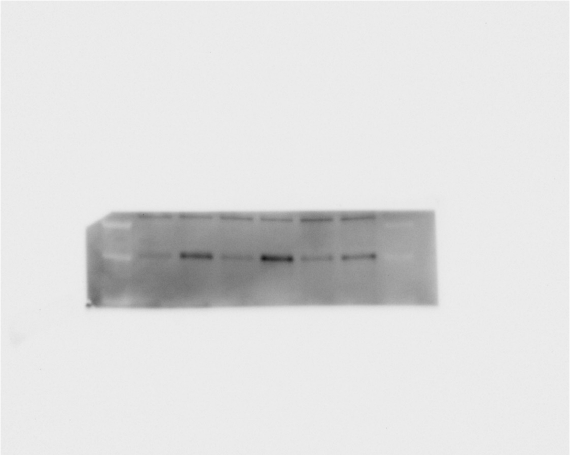

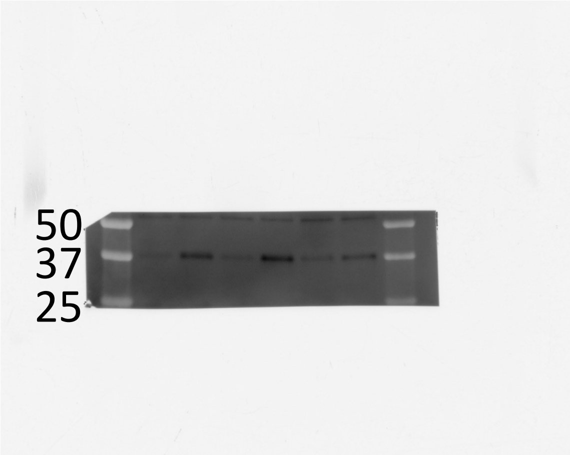


FOXB1

Sample loading order, left to right:
7 uL ladder
EV
HA-CIC(ex20)::NUTM1(ex6)
HA-CIC(ex20)::NUTM1(ex6) E
HA-CIC(ex20)::NUTM1(ex6) E.1
HA-CIC(ex20)::NUTM1(ex6) E.2
HA-CIC(ex20)::NUTM1(ex6) E.3
3 uL ladder

### Figure 4 Panel E

Left = Chemiluminescence, Right = composite of chemi + visual light

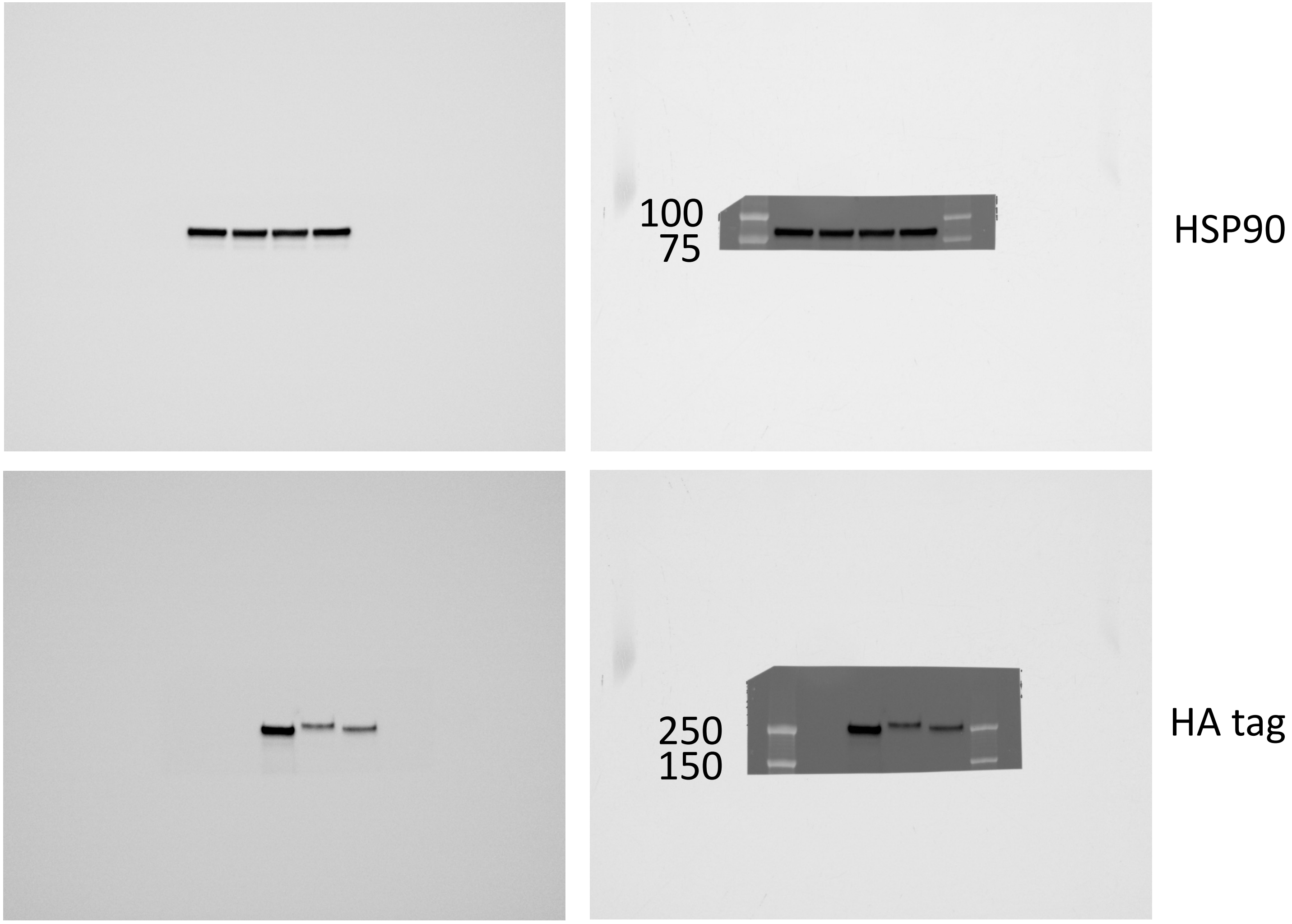


Ponceau S

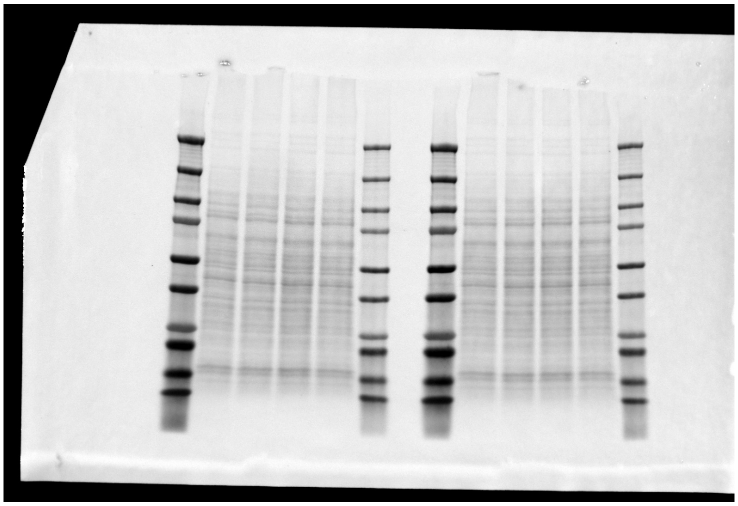

Sample loading order, left to right:
7 uL ladder
EV
HA-CIC::DUX4
HA-CIC::DUX4 +E
HA-CIC::DUX4 +E.2
3 uL ladder

Note: HA tag, FOXB1, and HSP90 were cut from the right side samples, ETV5 from the left.

### Figure 4 Panel E (continued)

Left = Chemiluminescence, Right = composite of chemi + visual light

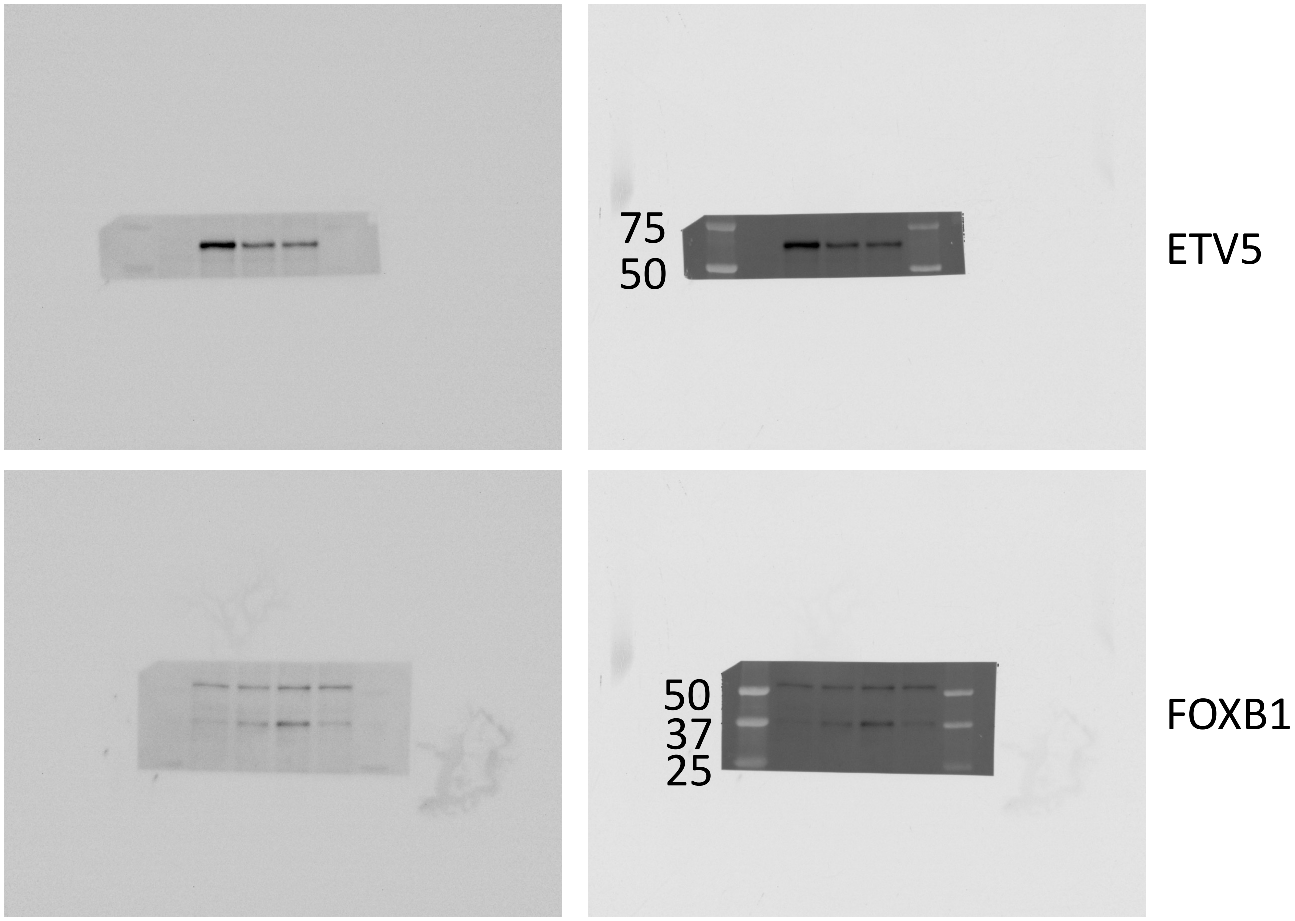


Sample loading order, left to right:
7 uL ladder
EV
HA-CIC::DUX4
HA-CIC::DUX4 +E
HA-CIC::DUX4 +E.2
3 uL ladder

### Supp Figure 3 Panel A

Left = Chemiluminescence, Right = composite of chemi + visual light

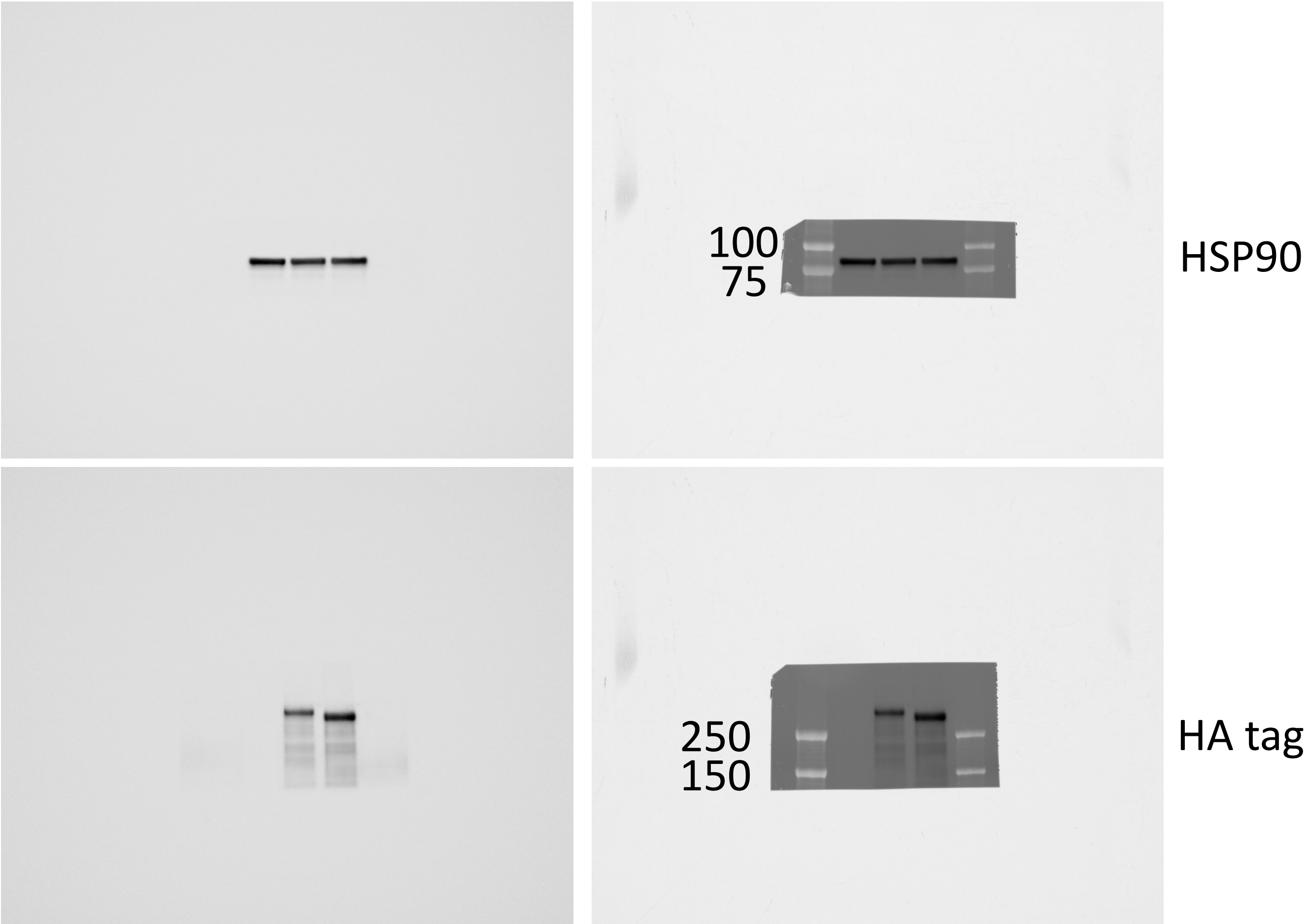


Ponceau S

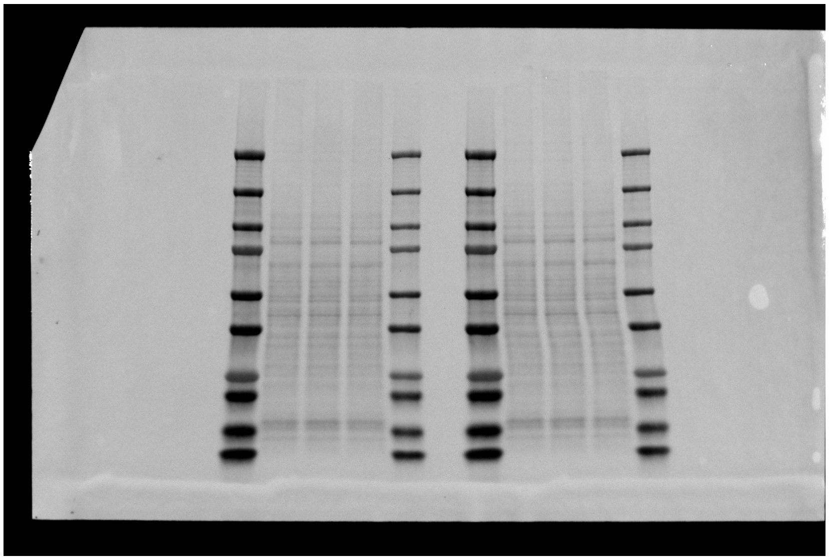

Sample loading order, left to right:
7 uL ladder
EV
HA-CIC(ex18)::NUTM1(ex3)
HA-CIC(ex18)::NUTM1(ex3) E
3 uL ladder

Note: HSP90 and FOXB1 were cut from the left side samples, HA tag and ETV5 from the right.

### Supp Figure 3 Panel A (continued)

Left = Chemiluminescence, Right = composite of chemi + visual light

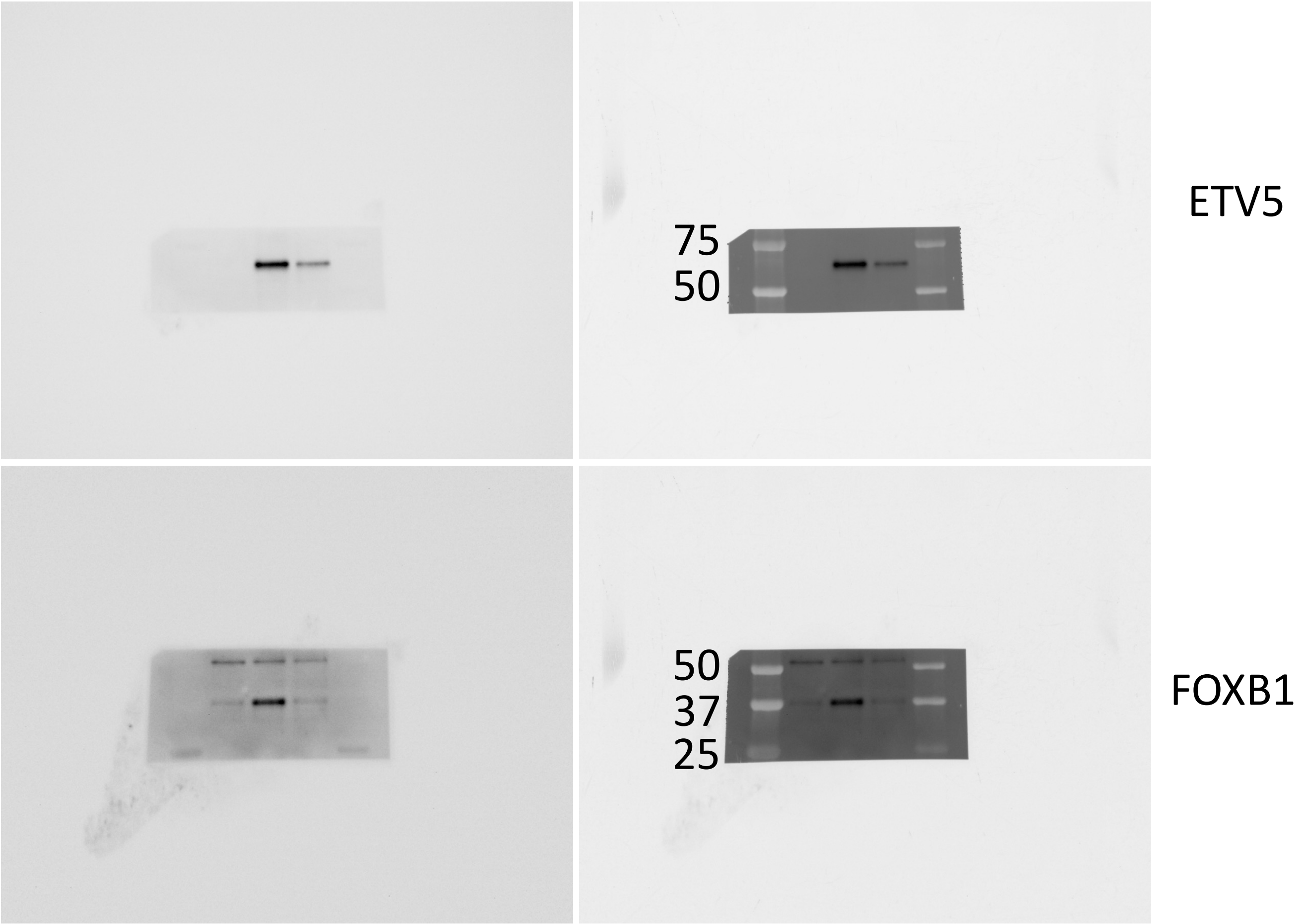


Sample loading order, left to right:
7 uL ladder
EV
HA-CIC(ex18)::NUTM1(ex3)
HA-CIC(ex18)::NUTM1(ex3) E
3 uL ladder

### Supp Figure 3 Panel B

Left = Chemiluminescence, Right = composite of chemi + visual light

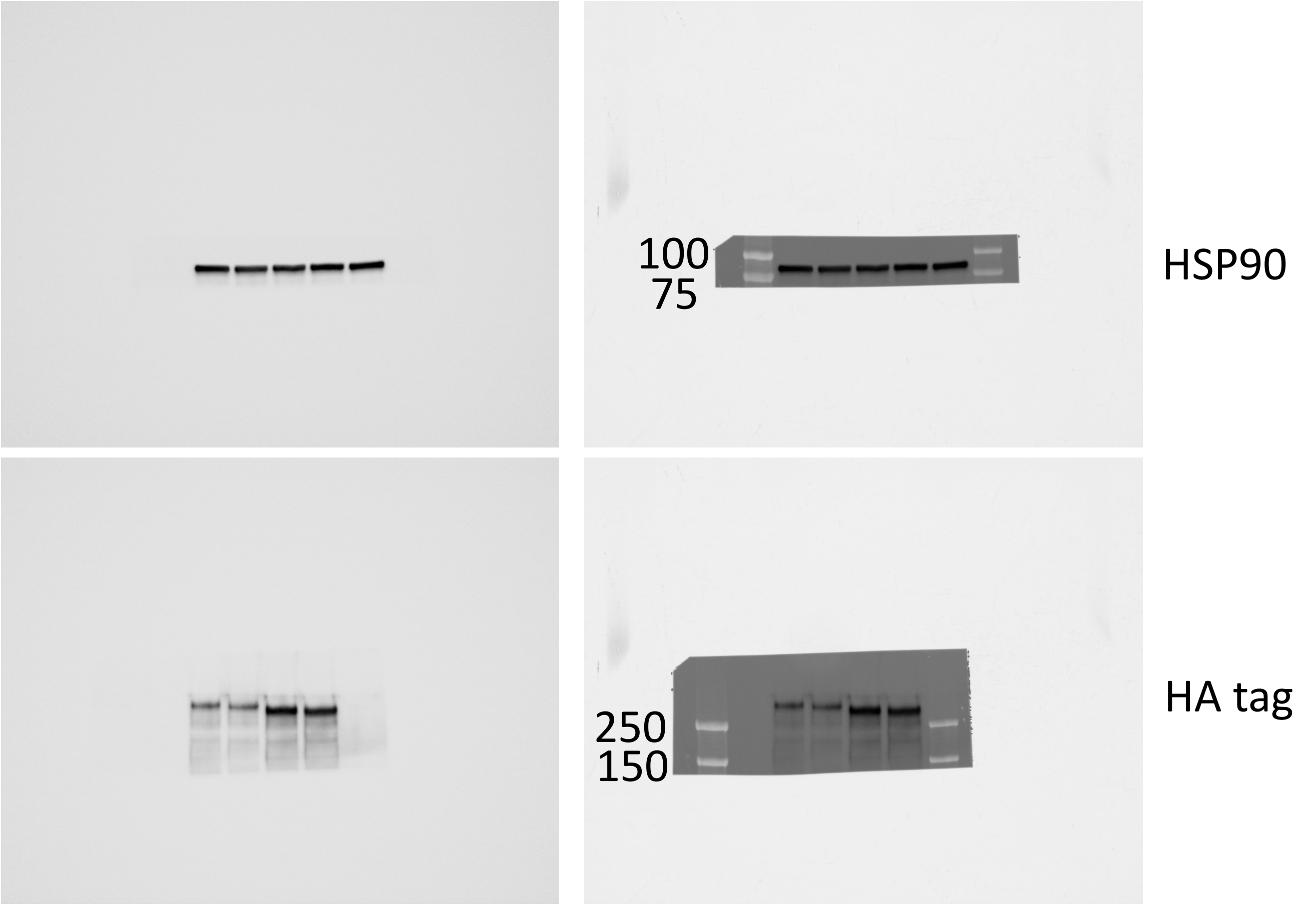


Ponceau S

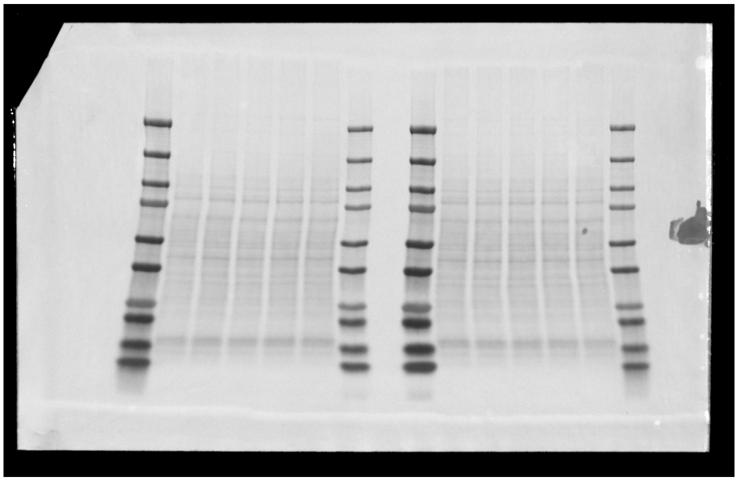

Sample loading order, left to right:
7 uL ladder
EV
HA-CIC(ex20)::NUTM1(ex6)
HA-CIC(ex20)::NUTM1(ex6) dTAD
HA-CIC(ex20)::NUTM1(ex6) E
HA-CIC(ex20)::NUTM1(ex6) dTAD & E
3 uL ladder

Note: HA tag and ETV5 were cut from the left side samples, HSP90 and FOXB1 from the right.

### Supp Figure 3 Panel B (continued)

Left = Chemiluminescence, Right = composite of chemi + visual light

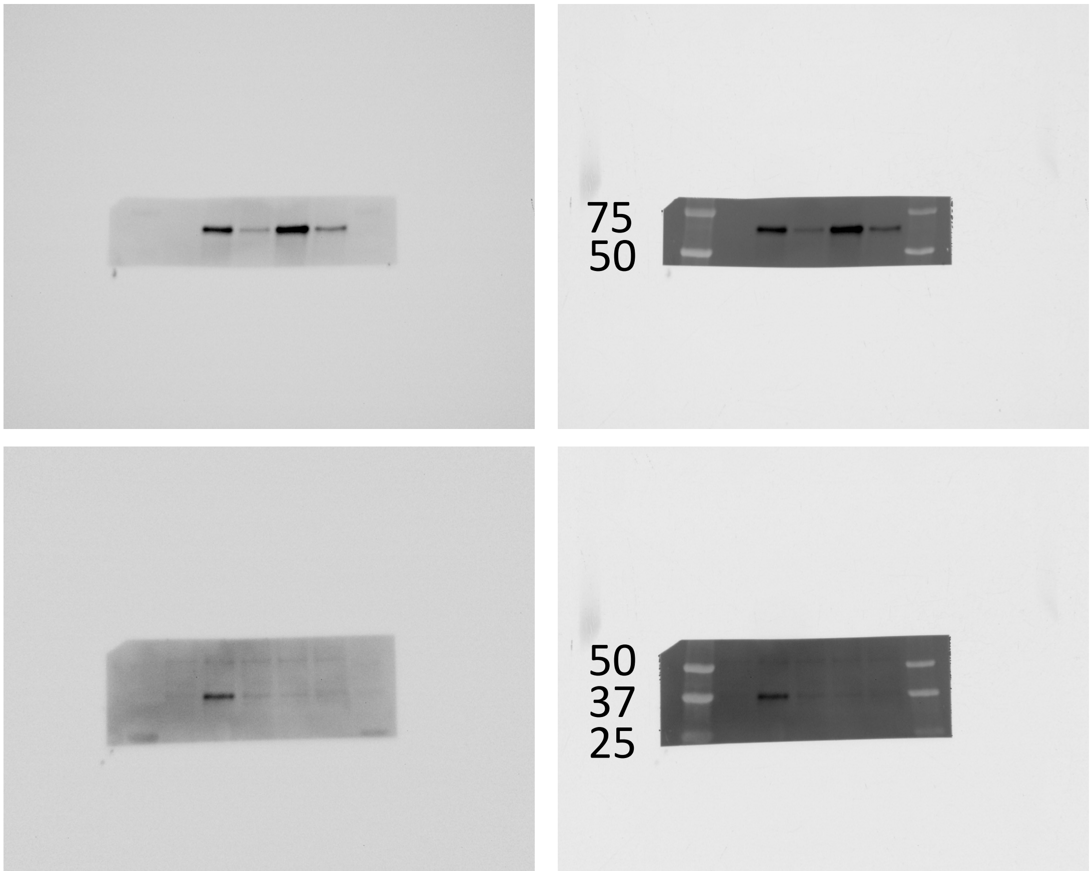


ETV5

FOXB1

Sample loading order, left to right:
7 uL ladder
EV
HA-CIC(ex20)::NUTM1(ex6)
HA-CIC(ex20)::NUTM1(ex6) dTAD
HA-CIC(ex20)::NUTM1(ex6) E
HA-CIC(ex20)::NUTM1(ex6) dTAD & E
3 uL ladder

### Supp Figure 3 Panel C

Left = Chemiluminescence, Right = composite of chemi + visual light

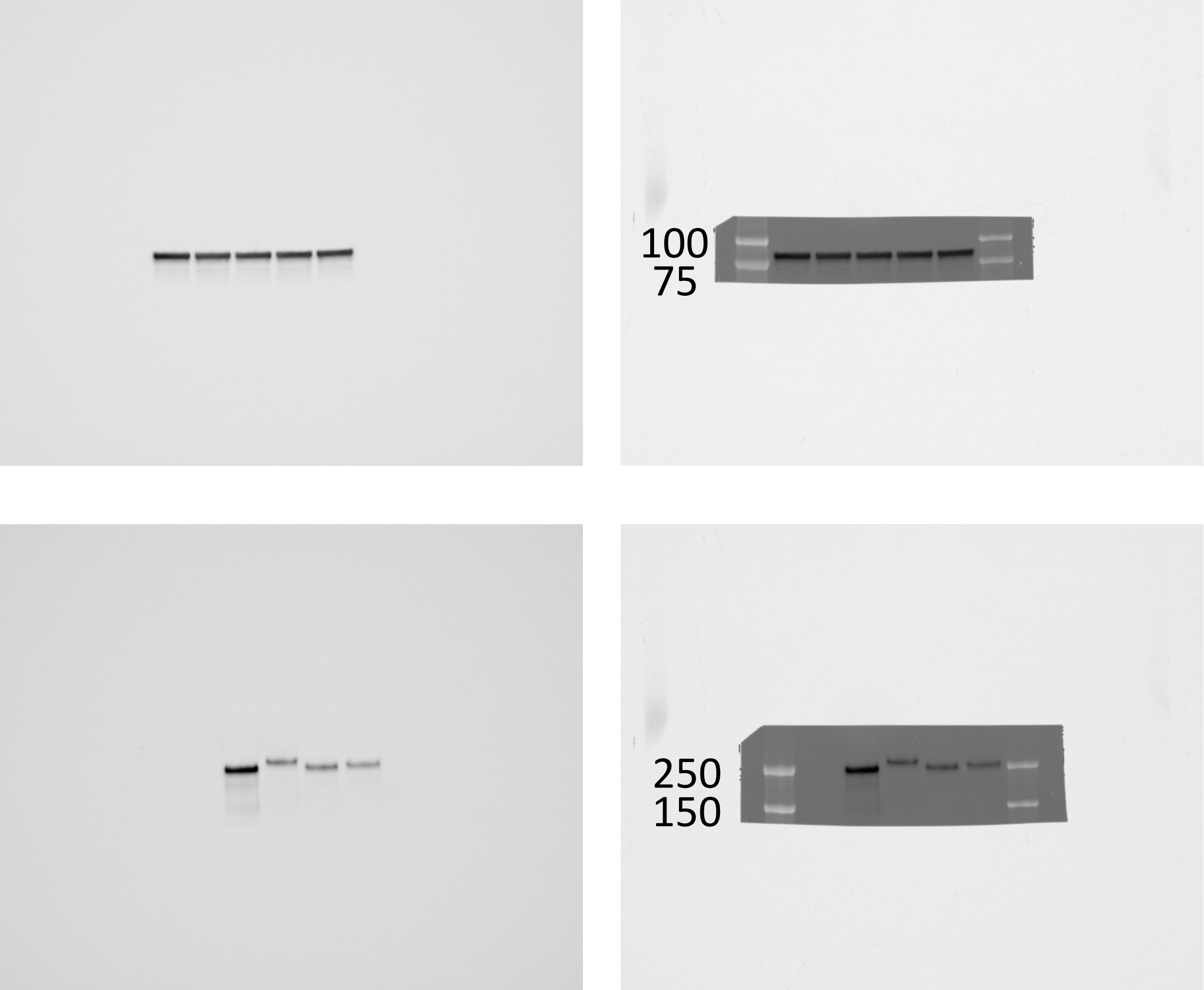


HSP90

HA tag

Ponceau S

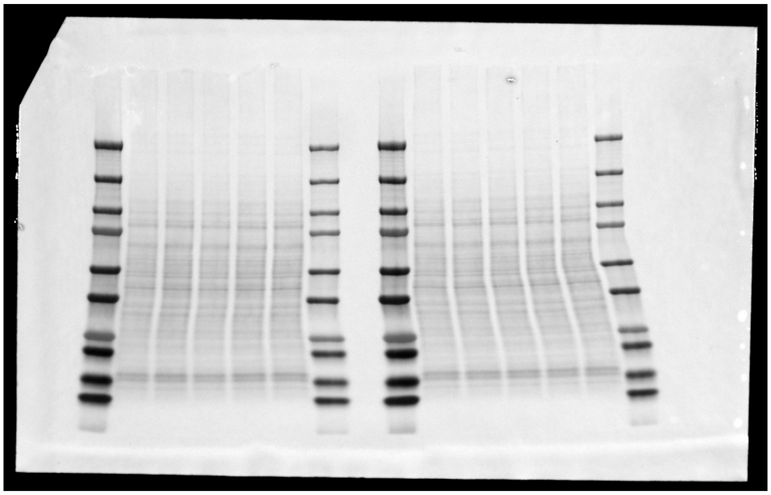

Sample loading order, left to right:
7 uL ladder
EV
HA-CIC::DUX4
HA-CIC::DUX4 +E
HA-CIC::DUX4 +E.2
HA-CIC::DUX4 +linker+E.2
3 uL ladder

Note: HA tag, ETV5, and FOXB1 were cut from the right side samples, HSP90 from the left.

### Supp Figure 3 Panel C (continued)

Left = Chemiluminescence, Right = composite of chemi + visual light

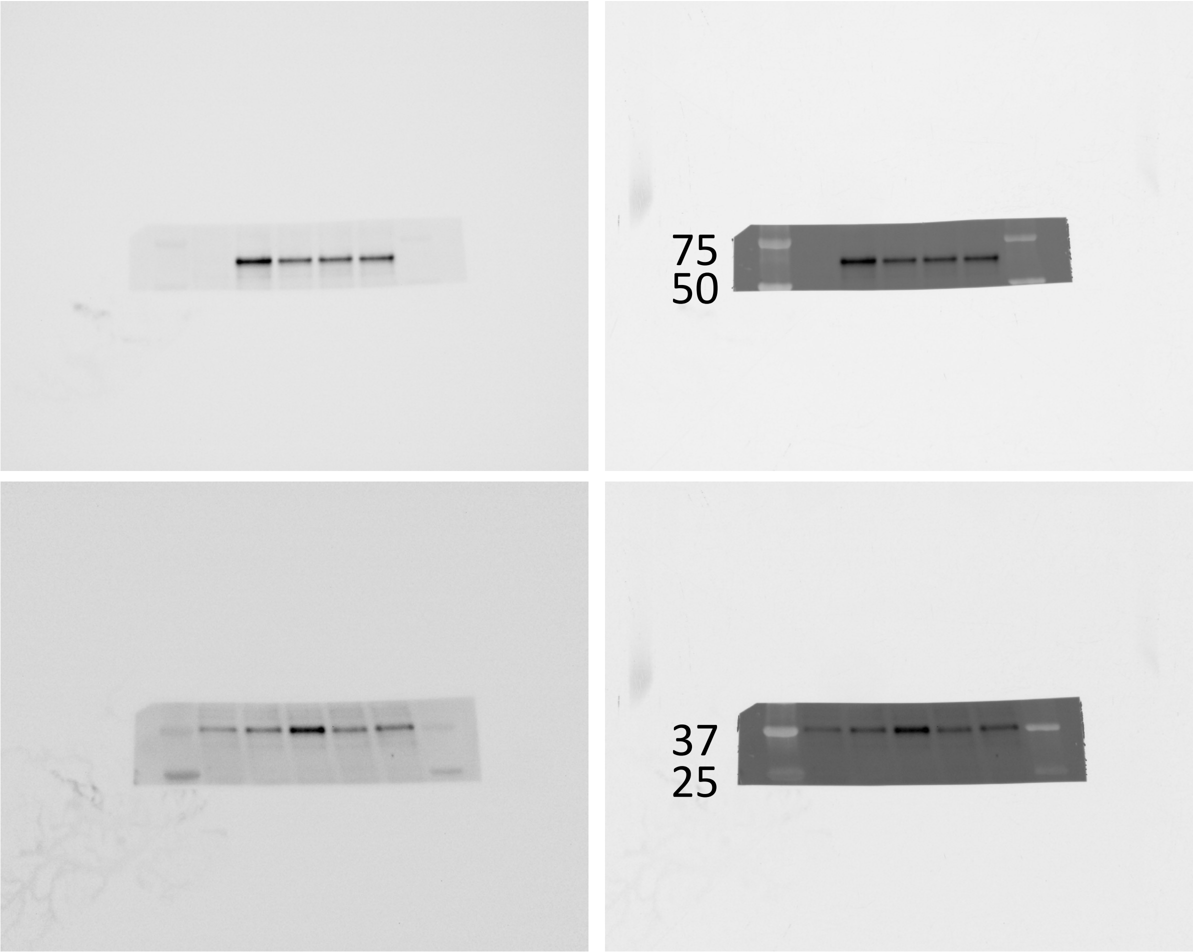


ETV5

FOXB1

Sample loading order, left to right:
7 uL ladder
EV
HA-CIC::DUX4
HA-CIC::DUX4 +E
HA-CIC::DUX4 +E.2
HA-CIC::DUX4 +linker+E.2
3 uL ladder

### Figure 6 Panel A

Left = Chemiluminescence, Right = composite of chemi + visual light

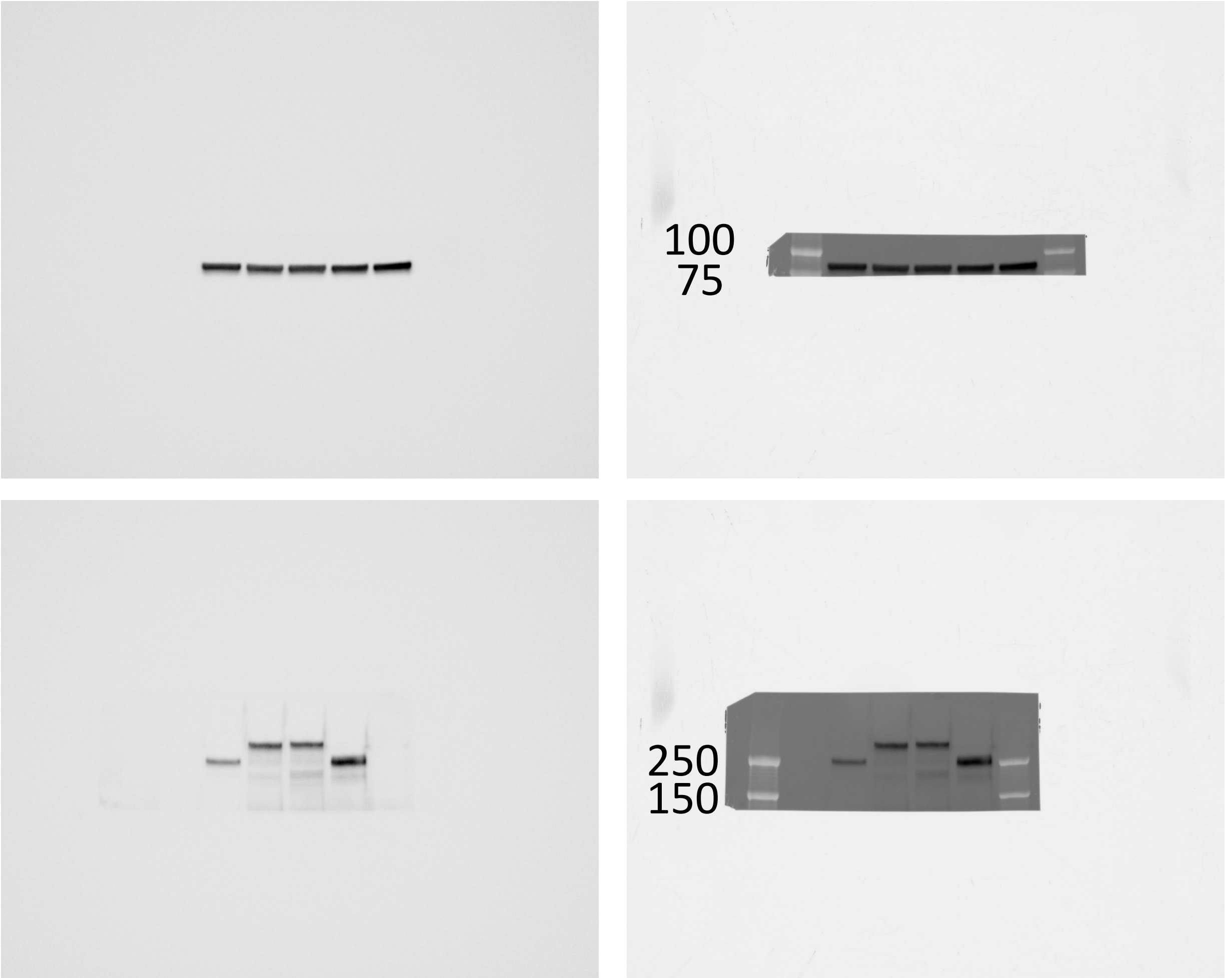


HSP90

HA tag

Ponceau S

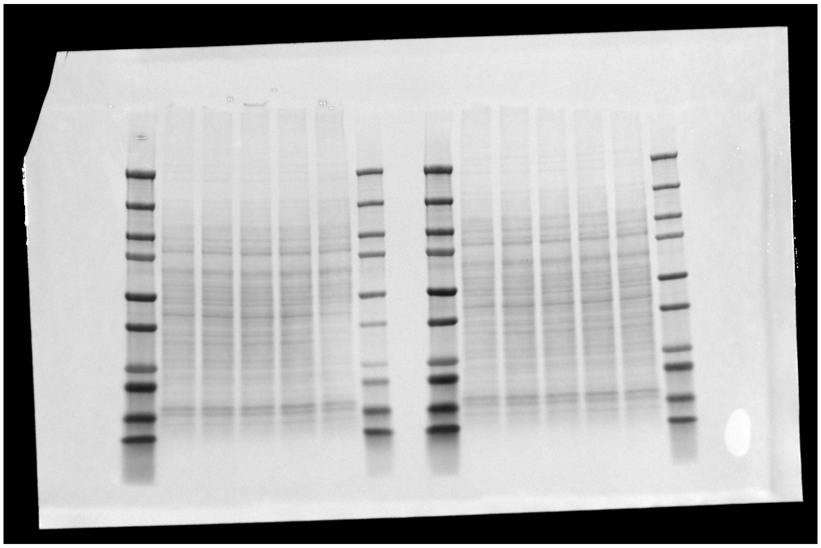

Sample loading order, left to right:
7 uL ladder
EV
HA-CIC::DUX4
HA-CIC(ex20)::NUTM1(ex6)
HA-CIC(ex18)::NUTM1(ex3)
HA-CIC::LEUTX
3 uL ladder

Note: All proteins cut from right set of samples due to clear transfer issue with the left (see small size of far right samples in that set).

### Figure 6 Panel A (continued)

Left = Chemiluminescence, Right = composite of chemi + visual light

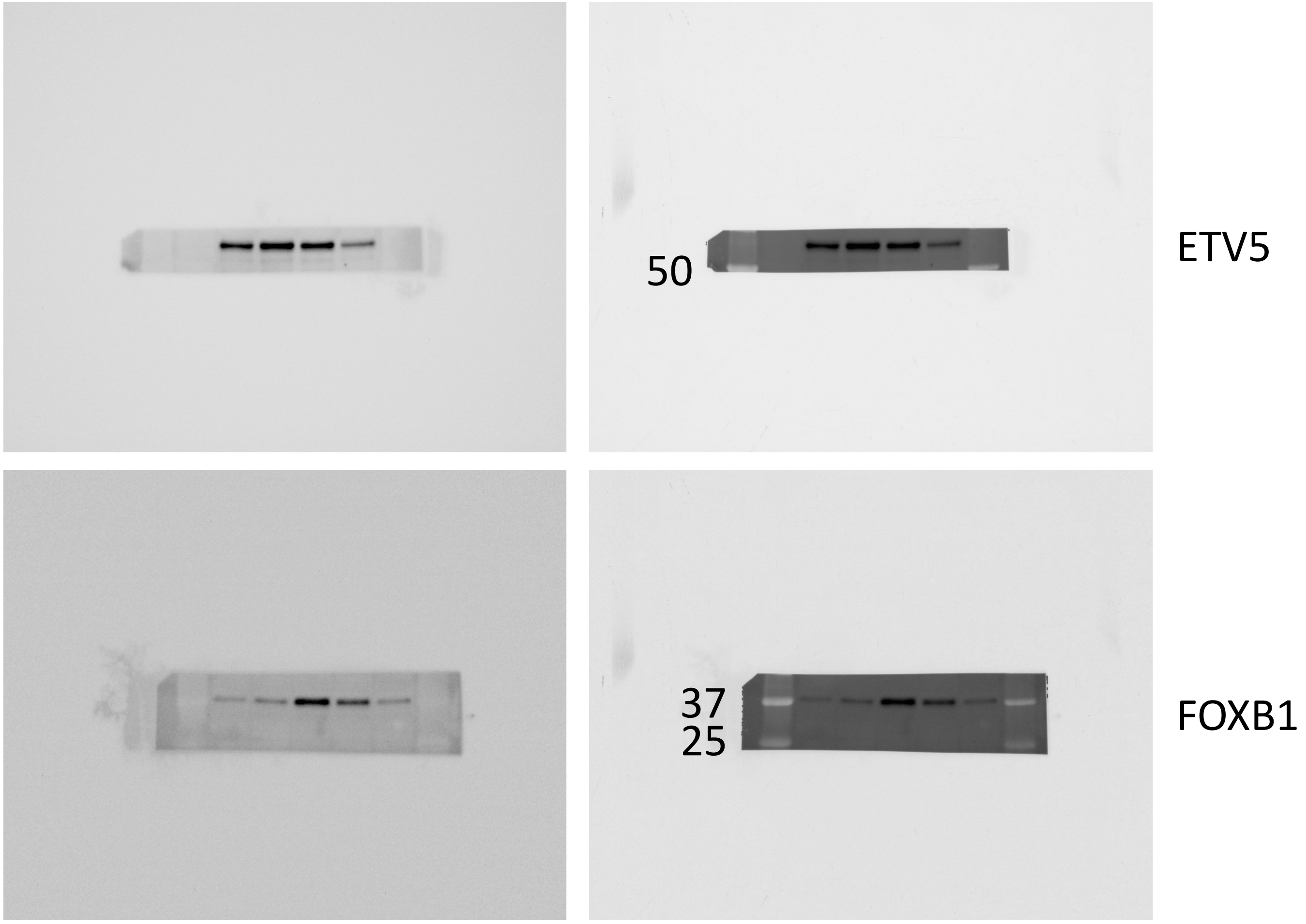


Sample loading order, left to right:
7 uL ladder
EV
HA-CIC::DUX4
HA-CIC(ex20)::NUTM1(ex6)
HA-CIC(ex18)::NUTM1(ex3)
HA-CIC::LEUTX
3 uL ladder

### Figure 6 Panel B

Left = Chemiluminescence, Right = composite of chemi + visual light

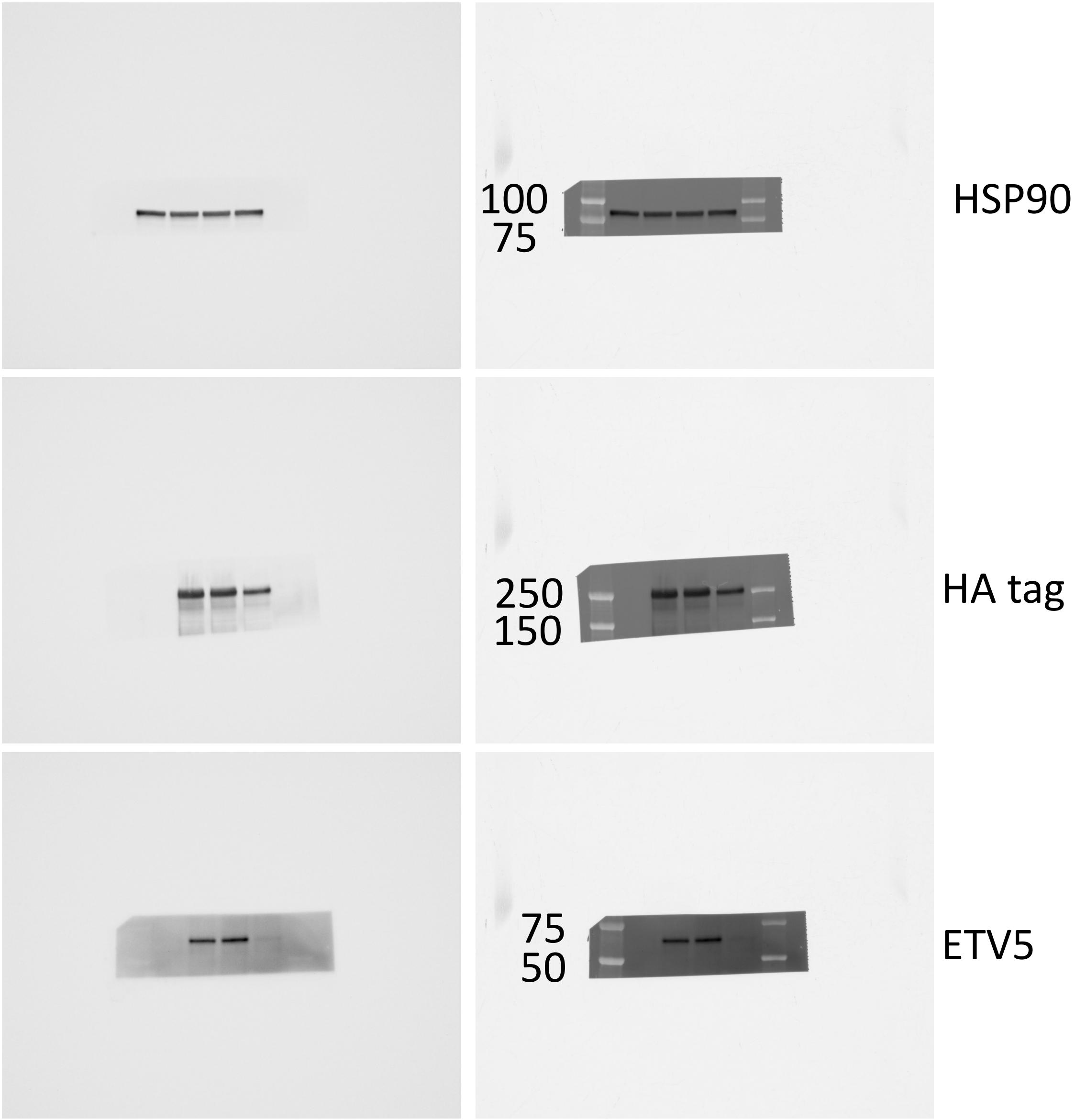


Ponceau S

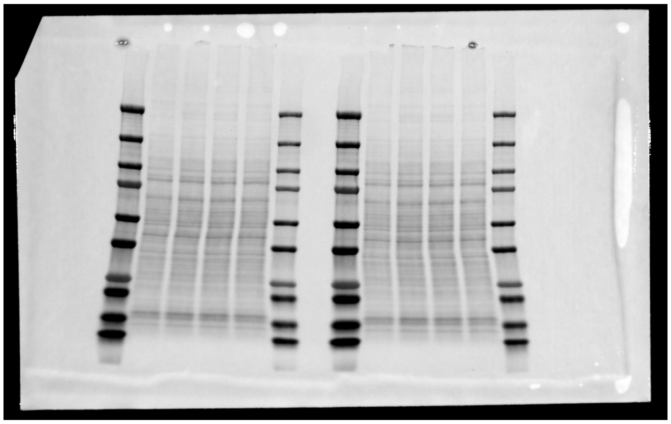

Sample loading order, left to right:
7 uL ladder
EV
HA-CIC::LEUTX
HA-CIC::LEUTX dTAD1
HA-CIC::LEUTX dTAD1&2
3 uL ladder

Note: HA tag and ETV5 were cut from the left side samples, HSP90 from the right.

### Supp Figure 5 Panel A

Left = Chemiluminescence, Right = composite of chemi + visual light

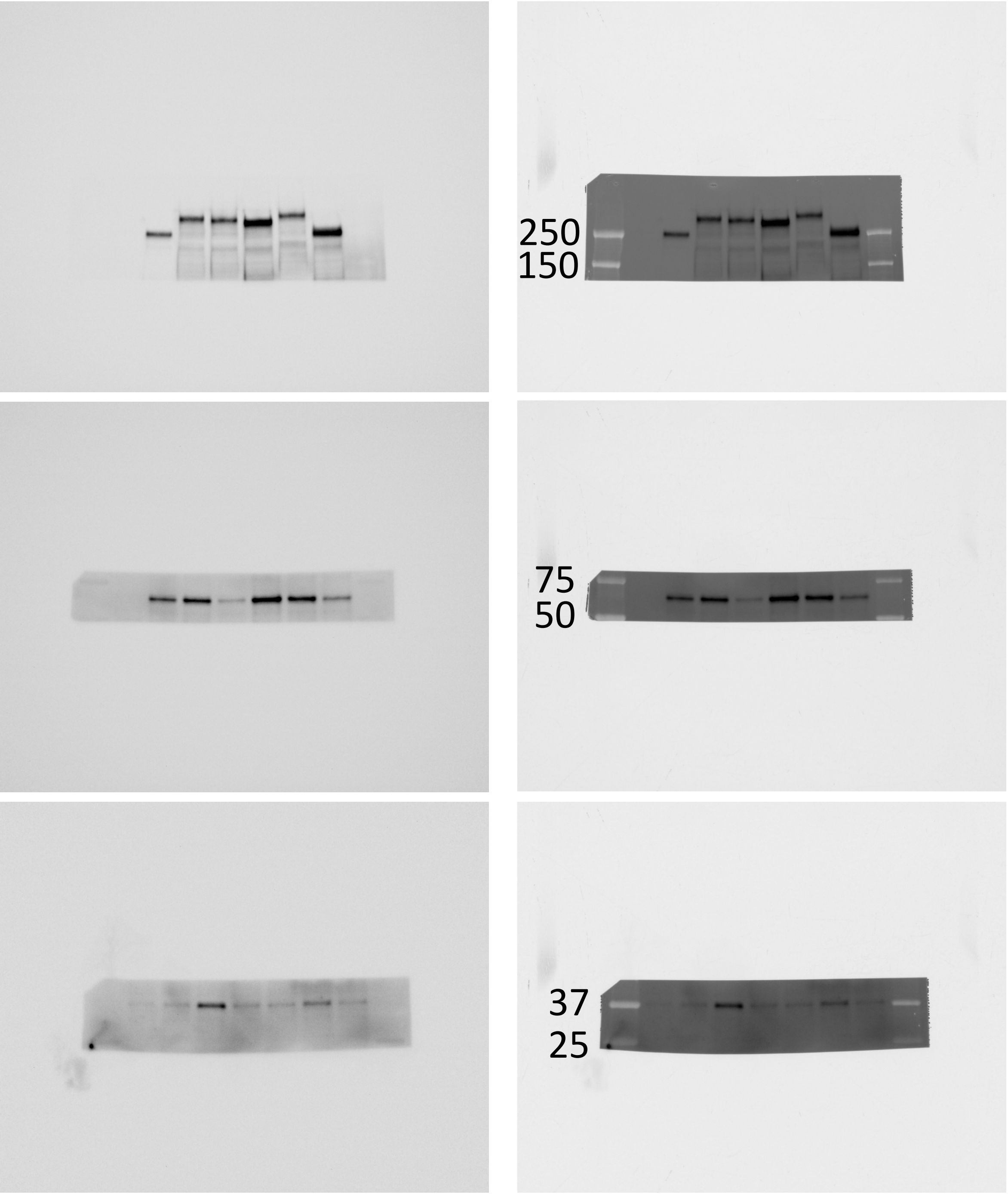


HA tag

ETV5

FOXB1

Ponceau S

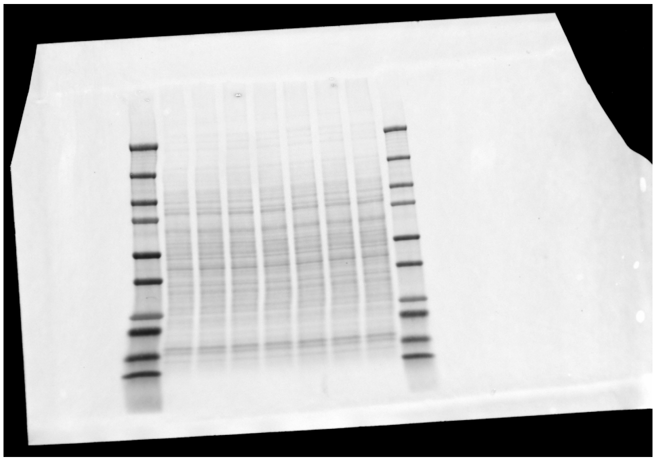

Sample loading order, left to right:

7 uL ladder

EV

HA-CIC::DUX4

HA-CIC(ex20)::NUTM1(ex6)

HA-CIC(ex20)::NUTM1(ex6) dTAD

HA-CIC(ex20)::NUTM1(ex6) E

HA-CIC(ex18)::NUTM1(ex3)

HA-CIC::LEUTX

3 uL ladder

### Figure 7 Panel A

Chemi + composite blots on following slides

Ponceau S

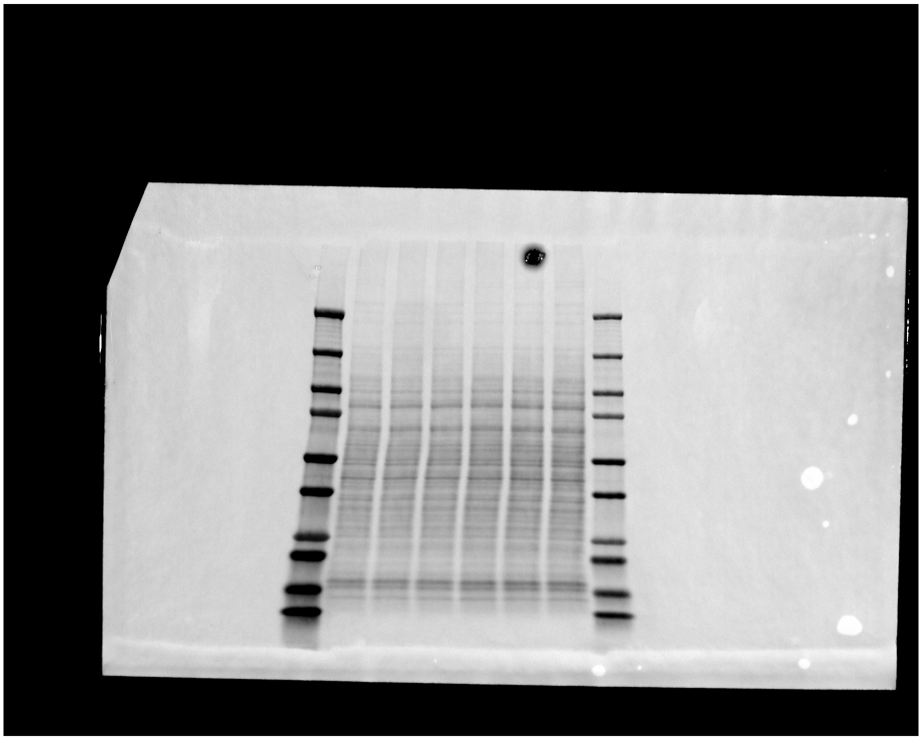

Note: HA tag and ETV5 were cut from these samples.

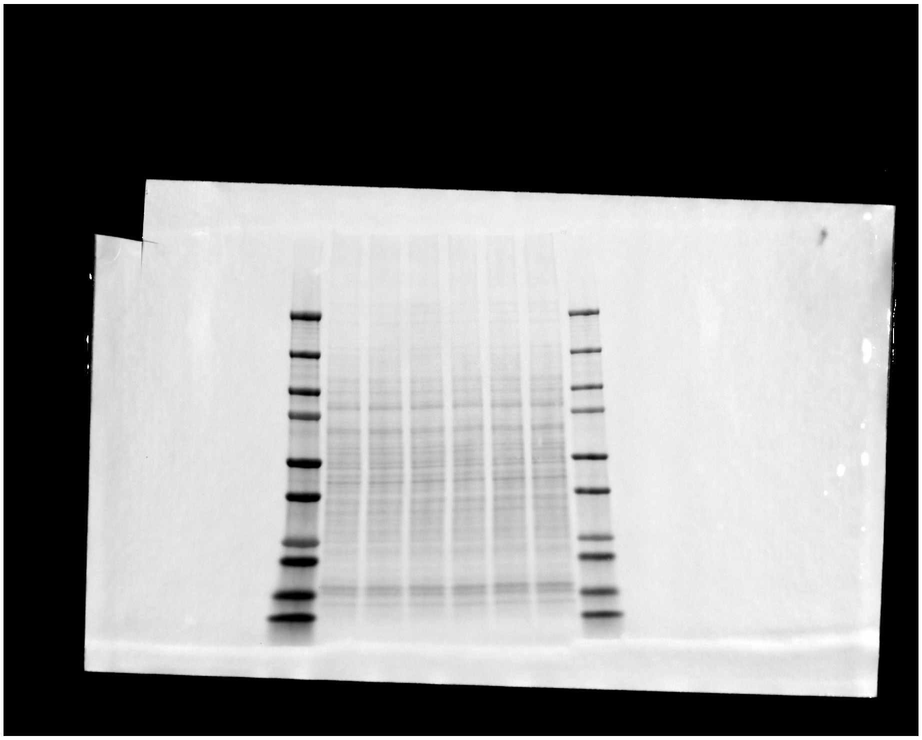

Note: HSP90 was cut from these samples.

Sample loading order, left to right:
7 uL ladder
EV rep 2
HA-ATXN1::DUX4 rep 2
HA-ATXN1::DUX4 delAXH rep 2
EV rep 3
HA-ATXN1::DUX4 rep 3
HA-ATXN1::DUX4 delAXH rep 3
3 uL ladder

### Figure 7 Panel A (continued)

Left = Chemiluminescence, Right = composite of chemi + visual light

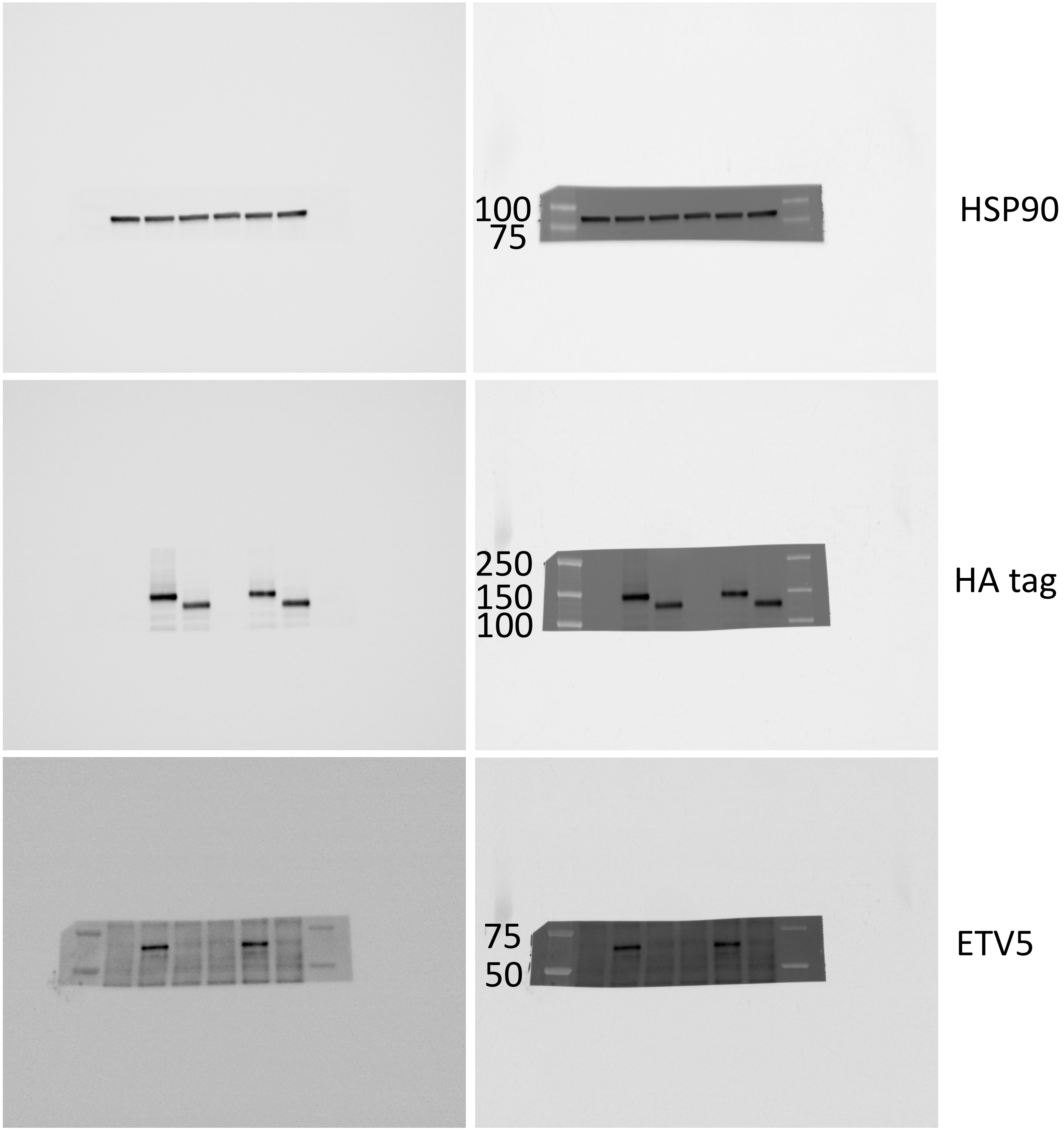


Sample loading order, left to right:
7 uL ladder
EV rep 2
HA-ATXN1::DUX4 rep 2
HA-ATXN1::DUX4 delAXH rep 2
EV rep 3
HA-ATXN1::DUX4 rep 3
HA-ATXN1::DUX4 delAXH rep 3
3 uL ladder
